# Accelerated brain ageing and disability in multiple sclerosis

**DOI:** 10.1101/584888

**Authors:** JH Cole, J Raffel, T Friede, A Eshaghi, W Brownlee, D Chard, N De Stefano, C Enzinger, L Pirpamer, M Filippi, C Gasperini, MA Rocca, A Rovira, S Ruggieri, J Sastre-Garriga, ML Stromillo, BMJ Uitdehaag, H Vrenken, F Barkhof, R Nicholas, O Ciccarelli, on behalf of the MAGNIMS study group

## Abstract

**Background:** Brain atrophy occurs in both normal ageing and in multiple sclerosis (MS), but it occurs at a faster rate in MS, where it is the major driver of disability progression. Here, we employed a neuroimaging biomarker of structural brain ageing to explore how MS influences the brain ageing process.

**Methods:** In a longitudinal, multi-centre sample of 3,565 MRI scans in 1,204 MS/clinically isolated syndrome (CIS) patients and 150 healthy controls (HCs) (mean follow-up time: patients 3⋅41 years, HCs 1⋅97 years) we measured ‘brain-predicted age’ using T1-weighted MRI. Brain-predicted age difference (brain-PAD) was calculated as the difference between the brain-predicted age and chronological age. Positive brain-PAD indicates a brain appears older than its chronological age. We compared brain-PAD between MS/CIS patients and HCs, and between disease subtypes. In patients, the relationship between brain-PAD and Expanded Disability Status Scale (EDSS) at study entry and over time was explored.

**Findings:** Adjusted for age, sex, intracranial volume, cohort and scanner effects MS/CIS patients had markedly older-appearing brains than HCs (mean brain-PAD 11⋅8 years [95% CI 9⋅1—14⋅5] versus −0⋅01 [−3⋅0—3⋅0], p<0⋅0001). All MS subtypes had greater brain-PAD scores than HCs, with the oldest-appearing brains in secondary-progressive MS (mean brain-PAD 18⋅0 years [15⋅4—20⋅5], p<0⋅05). At baseline, higher brain-PAD was associated with a higher EDSS, longer time since diagnosis and a younger age at diagnosis. Brain-PAD at study entry significantly predicted time-to-EDSS progression (hazard ratio 1⋅02 [1⋅01—1⋅03], p<0⋅0001): for every 5 years of additional brain-PAD, the risk of progression increased by 14⋅2%.

**Interpretation:** MS increases brain ageing across all MS subtypes. An older-appearing brain at baseline was associated with more rapid disability progression, suggesting ‘brain-age’ could be an individualised prognostic biomarker from a single, cross-sectional assessment.

**Funding:** UK MS Society; National Institute for Health Research University College London Hospitals Biomedical Research Centre.

## Research in context

### Evidence before this study

We searched Pubmed and Scopus with the terms “multiple sclerosis” and “brain ageing” or “brain age” and “neuroimaging” or “MRI” for studies published before 15^th^ March 2019. This searched return no studies of brain ageing in multiple sclerosis. We also searched the pre-print server for biology, bioRxiv, and found one manuscript deposited, though this study has yet to appear in a peer-reviewed journal. This study found a strong effect of multiple sclerosis on the apparent age of the brain, though was only cross-sectional, was from a single centre, did not consider disease subtypes and did not consider the relevance of clinical characteristics for brain ageing. Therefore, although there is strong prior evidence of the importance of brain atrophy in multiple sclerosis, there was no information on how the nature of this atrophy relates to brain ageing.

### Added value of this study

Here we demonstrate for the first time that the progressive atrophy in multiple sclerosis patients results in an acceleration of age-related changes to brain structure. Using a large multi-centre study, our data strongly support the idea that brain ageing is increased in multiple sclerosis, and that this is apparent across disease subtypes, including those with very early disease - Clinically Isolated Syndrome. Of particular value is the demonstration that baseline brain-age can be used to predict future worsening of disability, suggesting that a general index of age-related brain health could have relevance in clinical practice for predicting which patients will go on to experience a more rapidly progressing disease course.

### Implications of all the available evidence

Combined with the single other available study, this work shows robust evidence for a cross-sectional influence of multiple sclerosis on the apparent age of the brain, under the brain-age paradigm. This paradigm provides a new approach to considering how multiple sclerosis effects the structure of the brain during ageing, suggesting that multiple sclerosis may result in both disease-specific insults (e.g., lesions) alongside changes that are less specific (e.g., atrophy) and seen in ageing and other diseases. Potentially, treatments that improve brain health during normal ageing could be used to benefit patients with multiple sclerosis. Finally, brain-age may also have prognostic clinical value as a sensitive, if non-specific, biomarker of future health outcomes.

## Introduction

In multiple sclerosis (MS), age has been implicated as the dominant driver of disease progression.^1^ Older age increases the risk of progression,^2^ irrespective of disease duration, and once progression starts, disability accrual is independent of previous evolution of the disease (presence or not relapses, or relapse rates.^3–5^ This raises the possibility that MS interacts with some of the neurobiological drivers of brain ageing, leading to acceleration of the process, hastening brain atrophy in some individuals and leading to poorer long-term outcomes.^6^

That diseases may impact rates of biological ageing has been previously mooted outside of the context of MS. Potentially, a disease has both a specific impact but also may trigger a sequence of events which result in an acceleration of the biological processes seen in normal ageing, both systemically^7^ and in the brain.^8,9^

Recently, methods have been developed for measuring the biological ageing of the brain; the so-called ‘brain-age’ paradigm.^10^ Brain-age uses machine-learning analysis to generate a prediction of an individual’s age (their brain-predicted age), based solely on neuroimaging data (most commonly 3D T1-weighted MRI). The comparison of an individual’s brain-predicted age with their chronological age thus gives an index of whether their brain structure appears ‘older’ or ‘younger’ than would be expected for their age. By subtracting chronological age from brain-predicted age one can derive a brain-predicted age difference (brain-PAD); a simple numerical value in the unit years which shows promise as a biomarker of brain ageing. For example, brain-age has been shown to predict the likelihood of conversion from mild cognitive impairment to Alzheimer’s^11,12^ as well as the risk of mortality.^13^ Moreover, there is evidence for increased brain ageing in other neurological conditions contexts: traumatic brain injury,^14^ HIV,^15^ Down’s syndrome,^16^ and epilepsy.^17^

Here we employ brain-age to assess the relationship between MS disease progression and the brain ageing process. Using longitudinal neuroimaging and clinical outcomes in a large cohort of MS patients and healthy controls (HCs), we tested the following hypotheses: (i) MS patients have older-appearing brains than HCs; (ii) In MS patients, there is a relationship between brain-predicted age difference and disability at study entry; (iii) Brain-predicted age difference increases over time as disabilities worsen; and (iv) Brain-predicted age difference at baseline predicts future disability progression.

## Methods

### Participants

This study used data collected from seven European MS centres (MAGNIMS: www.magnims.eu) and Imperial College London on n=1,354 participants (table 1), largely overlapping with our previous work (detailed in the appendix table S1).^18^ Patients had all received a diagnosis of MS according to 2010 McDonald Criteria^19^ or CIS.^20^ MS/CIS patients were scored on the Expanded Disability Status Scale (EDSS).^21^ HCs without history of neurological or psychiatric disorders were also included (n=150). For longitudinal imaging analysis, participants were required to have undergone at least two high-resolution T1-weighted MRI acquired with the same protocol with an interval of ≥1 month.

**Table 1:** Characteristics of MS/CIS patients and healthy controls.

|  | MS/CIS patients | Healthy controls |
| --- | --- | --- |
| N | 1204 | 150 |
| N with follow-up data | 1155 | 111 |
| Female (n, %) | 771 (64) | 82 (55) |
| Number of scans per participant<br>mean ± SD [range] | 2.61 ± 1.01 [1-7] | 2.82 ± 1.90 [1-10] |
| Length of follow-up (years)<br>mean ± SD [range] | 3.41 ± 3.15 [0.2-15] | 1.97 ± 1.38 [0.5-6.0] |
| Age at baseline scan (years) | 39.41 ± 10.76 [15-68] | 37.29 ± 9.96 [23-66] |
| Brain-predicted age at baseline<br>(years) | 50.27 ± 14.90 [7.4-92] | 38.43 ± 11.12 [14.5-70] |
| Baseline subtype (%):<br>CIS/RRMS/SPMM/PPMS | 296 (25) / 677 (56) / 111 (9) / 120 (10) | - |
| Time since **clinical** diagnosis at<br>baseline (years) | 7.26 ± 7.96 [0-48] | - |
| EDSS at baseline<br>mean ± SD [range] | 2.60 ± 1.95 [0-9] | - |
| Disease-modifying treatment at<br>baseline (n, %): yes/no/unknown | 475 (39) / 675 (56) / 54 (5) | - |
| CIS = clinically isolated syndrome, EDSS = Expanded Disability Status Scale, RRMS = relapsing remitting MS, SD = standard deviation, SPMS = secondary progressive MS, PPMS = primary progressive MS. |  |  |

The final protocol for this study was reviewed and approved by the European MAGNIMS collaboration for analysis of pseudo-anonymized scans and the Imperial NHS Trust (London Riverside Research Ethics Committee: 14/LO/0343).

### EDSS progression

Time-to-event, where a progression event was an individual’s progression on the EDSS, was defined as per our previous work^18^: when a patient showed a longitudinal change of: a 1⋅5-point increase in EDSS if the baseline EDSS was 0; a 1-point increase if baseline EDSS was 1 to 6 inclusive; and a 0⋅5-point increase if EDSS was greater than 6.

### Neuroimaging acquisition

Overall, 3,565 T1-weighted MRI scans were used in the study according to local MRI protocols, which used similar acquisition parameters. Thirteen different scanners (Siemens, GE, Philips) were used in patients recruited from 1998 onwards (see appendix table S1).

### Machine-learning brain-predicted age analysis

Brain-predicted age calculation followed our previously established protocol.^15^ In brief, all structural images were pre-processed using SPM12 to generate grey matter (GM), white matter (WM) segmentations. Visual quality control was then conducted to verify segmentation accuracy; all images were included. Segmented GM and WM images were then non-linearly registered to a custom template (based on the training dataset). Finally, images were affine registered to MNI152 space (voxel size = 1⋅5mm^3^), modulated and smoothed (4mm). Summary volumetric measures of GM, WM, cerebrospinal fluid (CSF) and intracranial volume (ICV) were also generated.

Brain-predicted ages were generated using Pattern Recognition for Neuroimaging Toolbox (PRoNTo v2⋅0, www.mlnl.cs.ucl.ac.uk/pronto) software.^22^ First, a model of healthy brain ageing was defined: brain volumetric data (from in a separate training dataset, n=2001 healthy people, aged 18-90; appendix table S2) were used as the independent variables in a Gaussian Processes regression, with age as the dependent variable. This regression model achieved a mean absolute error (MAE) of 5⋅02 years, assessed using ten-fold cross-validation, which explained 88% of the variance in chronological age.

Next, the coefficients from the full historical training model (n=2001) were applied to the current test data (i.e., MS/CIS patients and HCs), to generate brain-predicted ages. These values were adjusted to remove age-related variance, by subtracting 3⋅33 and then dividing by 0⋅91 (the intercept and slope of a linear regression of brain-predicted age on chronological age in the training dataset).

Finally, brain-PAD scores were calculated by subtracting chronological age from brain-predicted age and used for subsequent analysis. A positive brain-PAD score indicates that the individual’s brain is predicted to be ‘older’ than their chronological age.

### Statistical analysis

Using brain-PAD values further statistical analysis was carried out to test our hypotheses, using R v3⋅4⋅3. A full list of R packages and versions is included in the accompanying R Notebook (appendix). We used linear mixed effects models, enabling incorporation of fixed and random effects predictors to model each given outcome measure. In these models, brain-PAD was used as the outcome variable. Each model included fixed effects of group (e.g., MS/CIS patient versus HCs; MS subtype [CIS, RRMS, SPMS, PPMS]), age, sex and ICV and random effects of MRI scanner field-strength and original study cohort (modelling intercept). Estimated marginal means and confidence intervals from linear models were calculated. This analysis was repeated using data from a single cohort from a single centre (UCL, London), where all MS subtypes were present.

A random effects meta-analysis was conducted to explore the heterogeneity of the group effects on brain-PAD across different study cohorts. Only cohorts that included HCs and MS or CIS patients were included in this analysis.

To establish whether brain volume measurements were driving the variability in brain-PAD, we performed a linear regression with hierarchical partitioning of variance, with brain-PAD as the outcome variable and age, sex, GM, WM and CSF volume as predictors.

Subsequent analyses were conducted to test for fixed-effect influences of EDSS score (MS and CIS patients), and time since clinical diagnosis and age at clinical diagnosis (MS patients only). Model fits were considered using F-tests and post-hoc pairwise comparisons using t-tests or Tukey tests where appropriate.

We explored how longitudinal changes in brain-PAD related to changes in disability over time in two ways: (i) by correlating annualised change in brain-PAD (i.e., the difference between first measured brain-PAD and last brain-PAD, divided by the interval in years) with the annualised change in EDSS score; (ii) by using linear mixed effects models to investigate group (MS/CIS vs., HCs; MS subtype) by time interactions. These analyses included a random effect of participant (modelling slope and intercept), alongside age, sex, ICV scanner and cohort effects.

Survival analysis, using a Cox proportional hazards regression, was used to test whether baseline brain-PAD predicted time-to-EDSS progression, including age at baseline MRI and sex as covariates.

We investigated the impact of MS lesions on brain-PAD in MS. Using cross-sectional data from a subset of n=575 MS/CIS patients, for which manually-annotated lesion maps were available, we explored the relationship between MS lesions and measurements of brain-PAD, using the FSL lesion-filling algorithm,^23^ by artificially removing lesions from T1-weighted MRI scans. Both ‘lesion-filled’ and ‘unfilled’ scans were run through the brain-age prediction procedure, then resulting brain-PAD scores compared.

### Role of the funding source

The funder of the study had no role in study design, data collection, data analysis, data interpretation, or writing of the report. The corresponding authors had full access to all the data in the study and had final responsibility for the decision to submit for publication.

## Results

### Multiple sclerosis is associated with older appearing brains

The MAGNIMS sample forms part of a well-characterised population (table 1). The combined cohort involves patients from six countries with a mean follow-up of 3⋅41 years in patients.

Patients with MS/CIS had markedly greater brain-PAD scores at time of initial MRI scan compared to HCs (estimated marginal means 11⋅8 years, [95% CI 9⋅1–14⋅5] versus −0⋅01 [95% CI −3⋅0–3⋅0]). When adjusted for the age, sex, intracranial volume, cohort and scanner effects, there was a statistically significant group mean difference in brain-PAD of 11⋅8 years (95% CI 9⋅9–13⋅8, p<0⋅0001).

Though there is considerable heterogeneity between the study cohorts, due to clinical characteristics and technical factors (e.g., MRI scanner system), the difference between MS/CIS and HCs was robust in a random-effects meta-analysis of a subset of the data; six London cohorts that included both MS/CIS patients and HCs (figure 1A). The heterogeneity in the group differences was substantial (I^2^ = 59%, [95% CI 3–91%]).

**Figure 1.**
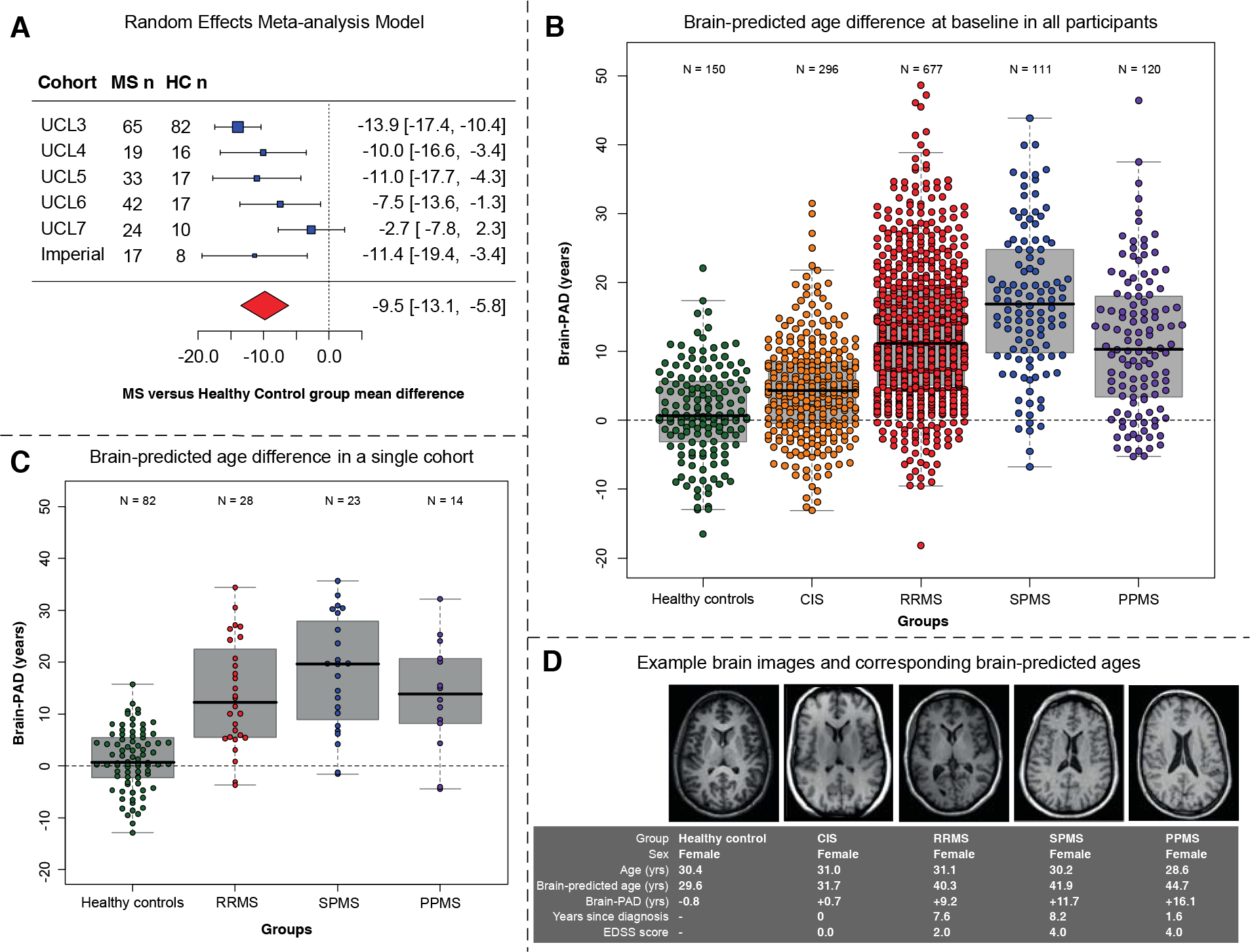
Brain-predicted age difference (Brain-PAD) for MS/CIS patients and healthy controls at baseline. A) A random-effects meta-analysis of the six cohorts that included both MS/CIS patients and HCs found the pooled effect of MS/CIS on brain-PAD compared to HCs was 9⋅45 years (95% CI 13⋅11– 5⋅80), across a total of n=200 MS/CIS patients and n=15HC. Heterogeneity was estimated at I^2^ = 59% [3–91%]. B) Grouped scatterplot depicting the distributions of brain-PAD at baseline, in years. Black lines represent the group median, shaded boxes show the inter-quartile range and whiskers 1⋅5 times the inter-quartile range from the median. C) Data from cohort “UCL3”, where all MS subtypes were present, confirms a similar result to the total cohort. D) Examples of how brain structure relates to brain-PAD, with axial slice from T1-weighted MRI from one healthy control and four individuals with CIS or MS, all females of a similar age. A control brain from a 30-year-old female with a brain-PAD of −0⋅8 years can be compared to a 31-year-old female with CIS, EDSS of 0⋅0 and a brain-PAD of +0⋅7 years, and 31-year-old with RRMS, EDSS of 2⋅0 and a brain-PAD of +9⋅2 years. In addition, we illustrate a 30-year-old with SPMS, EDSS of 4⋅0 and a brain-PAD of +11⋅7 years and a 28-year-old with PPMS, EDSS of 4⋅0 and a brain-PAD of +16⋅7 years. CIS = clinically isolated syndrome, RRMS = relapsing remitting MS, SPMS = secondary progressive MS, PPMS = primary progressive MS.

MS subtype (CIS, RRMS, SPMS, PPMS) significantly influenced brain-PAD (F_3,802⋅25_ = 29⋅9, p<0⋅0001, figure 1B). Estimated marginal mean brain-PAD per subtype were: CIS 6⋅3 years [95% CI 3⋅9–8⋅8], RRMS 12⋅4 years [95% CI 10⋅3–14⋅5], SPMS 18⋅0 years [95% CI 15⋅4–20⋅5], and PPMS 12⋅4 years [95% CI 9⋅7–15⋅2]. Post-hoc pairwise group comparison (appendix table S3) showed statistically significant differences (p<0⋅05) in brain-PAD between each subtype and HCs, and between CIS patients and each of the three MS groups (RRMS, SPMS, PPMS). SPMS patients showed significantly greater brain-PAD compared to both RRMS and PPMS patients. The difference in brain-PAD between PPMS and RRMS was not statistically significant (p=0⋅62). The findings of differences in brain-PAD between MS subtypes were replicated in a single cohort from a single centre, where all subtypes were present (cohort UCL3, figure 1C). Brain-PAD scores and corresponding T1-weighted MRI scans of individual female participants with similar ages, but with different subtypes of MS, are illustrated in figure 1D.

### The relationship between lesions, brain volume and brain-PAD

We considered the impact of lesions of brain-PAD, by comparing brain-PAD values on a single MRI scan from n=575 patients with both a lesion-filled and unfilled version of the same image. The correlation between brain-predicted age using filled and unfilled scans was r=0⋅99, p<0⋅0001 (appendix figure S1A) suggesting that the presence of lesions did not overly influence the brain-PAD values used throughout the study (which were unfilled). A Bland-Altman plot showed a mean difference between filled and unfilled scans was −0⋅28 ±1⋅29 years with no systemic bias caused by lesion filling evident, though there was increased variability between ages 60-70 years (appendix figure S1B).

When we examined whether brain volume measurements were driving the variability in brain-PAD, we found that the combination of chronological age, sex, GM, WM and CSF volume explained about half of the variation in brain-PAD (adjusted R^2^=0⋅48) (appendix table S4). Age (9% variance explained), GM (15%) and CSF (20%) volume were major contributors to variance in brain-PAD.

### Brain-PAD at baseline is associated with disability, age at diagnosis, and time since clinical diagnosis

At baseline, a higher brain-PAD was associated with higher disability, as measured by the EDSS, when adjusting for age, sex, ICV, scanner and cohort: for every 1⋅74 years increase in brain-PAD, the EDSS increased by one point (95% CI 1⋅39–2⋅09], p<0⋅0001). This effect was consistent across the MS subtypes with no statistically significant interaction between subtype and EDSS score (F_3,1159⋅6_ = 1⋅12, p=0⋅34; figure 2A). With the same adjustments, a higher brain-PAD was associated with both younger age at diagnosis and longer time since diagnosis: for every year increase in brain-PAD, the age at diagnosis was reduced by 0⋅45 years (95% CI −0⋅55–−0⋅36], p<0⋅0001); for every 0⋅48 year increase in brain-PAD, the time since diagnosis increased by one year (95% CI 0⋅40–0⋅57, p<0⋅0001). There was an interaction between subtype (RRMS, PPMS and SPMS) and age at diagnosis (F_2,883⋅9_ = 3⋅20, p=0⋅041; figure 2B), driven by the presence of stronger relationships between brain-PAD and age at diagnosis in PPMS (slope beta −0⋅51) and SPMS (beta−0⋅57) compared to RRMS (beta −0⋅36), though all were significant (p<0⋅001). For time since diagnosis, the interaction was also significant (F_2,690⋅5_ = 3⋅61, p=0⋅028; figure 2C), driven by the presence of relationships in RRMS (beta 0⋅48, p<0⋅0001) and SPMS (beta 0⋅26, p=0⋅01), not observed in PPMS (beta 0⋅12, p=0⋅47).

**Figure 2.**
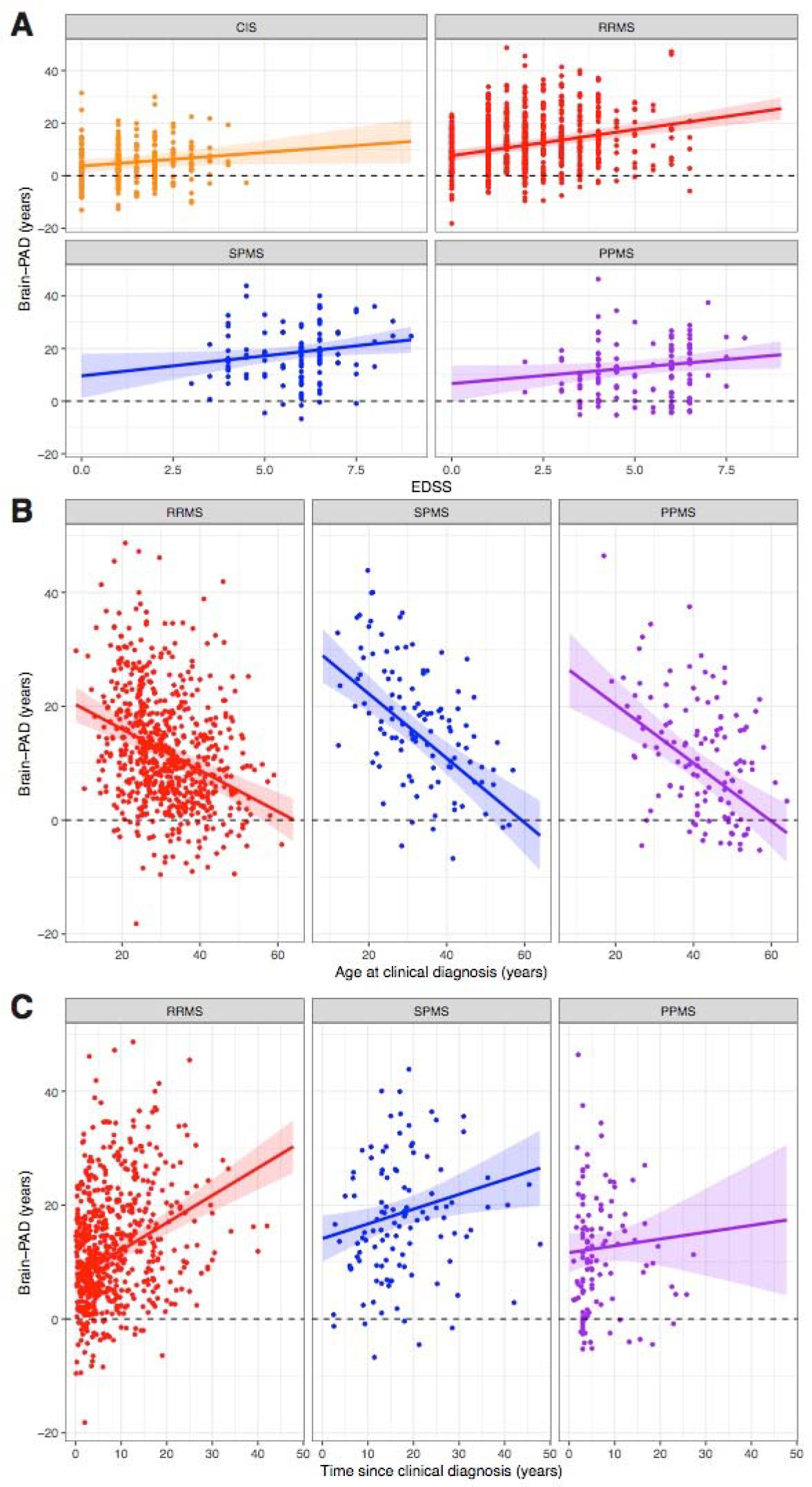
Scatterplot of brain-predicted age difference by age at diagnosis, time since diagnosis and EDSS score. A) Baseline EDSS score (x-axis) and concurrent brain-PAD (y-axis). B) Age at clinical diagnosis at first scan (x-axis) and concurrent brain-PAD (y-axis). C) Time since diagnosis at baseline (x-axis) and concurrent brain-PAD (y-axis). Panels show patients with RRMS, SPMS and PPMS separately. Panels show patients with RRMS, SPMS and PPMS separately. Panels show patients with CIS, RRMS, SPMS and PPMS separately. Lines represented the linear regression lines calculated per group, and shaded areas are the 95% confidence intervals. CIS = clinically isolated syndrome, RRMS = relapsing remitting MS, SPMS = secondary progressive MS, PPMS = primary progressive MS.

### Brain-PAD increase over time correlates with EDSS worsening

In patients who had two or more scans (n=1155), there was a significant positive correlation between annualised change in brain-PAD and annualised change in EDSS (Pearson’s r=0⋅26, p<0⋅0001). There was a significant interaction between EDSS change and disease subtype, when predicting brain-PAD change in linear model (F_4,1092_ = 24⋅5, p=0⋅009). The slopes were positive in CIS (beta 0⋅84, p=0⋅0001) and RRMS (beta 1⋅25, p<0⋅0001), though flatter in PPMS (beta 0⋅59, p=0⋅090) and negative (though not significant) in SPMS (beta −0⋅70, p=0⋅29; figure 3). To explore the latter finding post-hoc, correlated baseline brain-PAD with the number of follow-up scans completed. This showed a significant inverse correlation (n=104, Spearman’s rho=−0⋅29, p=0⋅0028).

**Figure 3.**
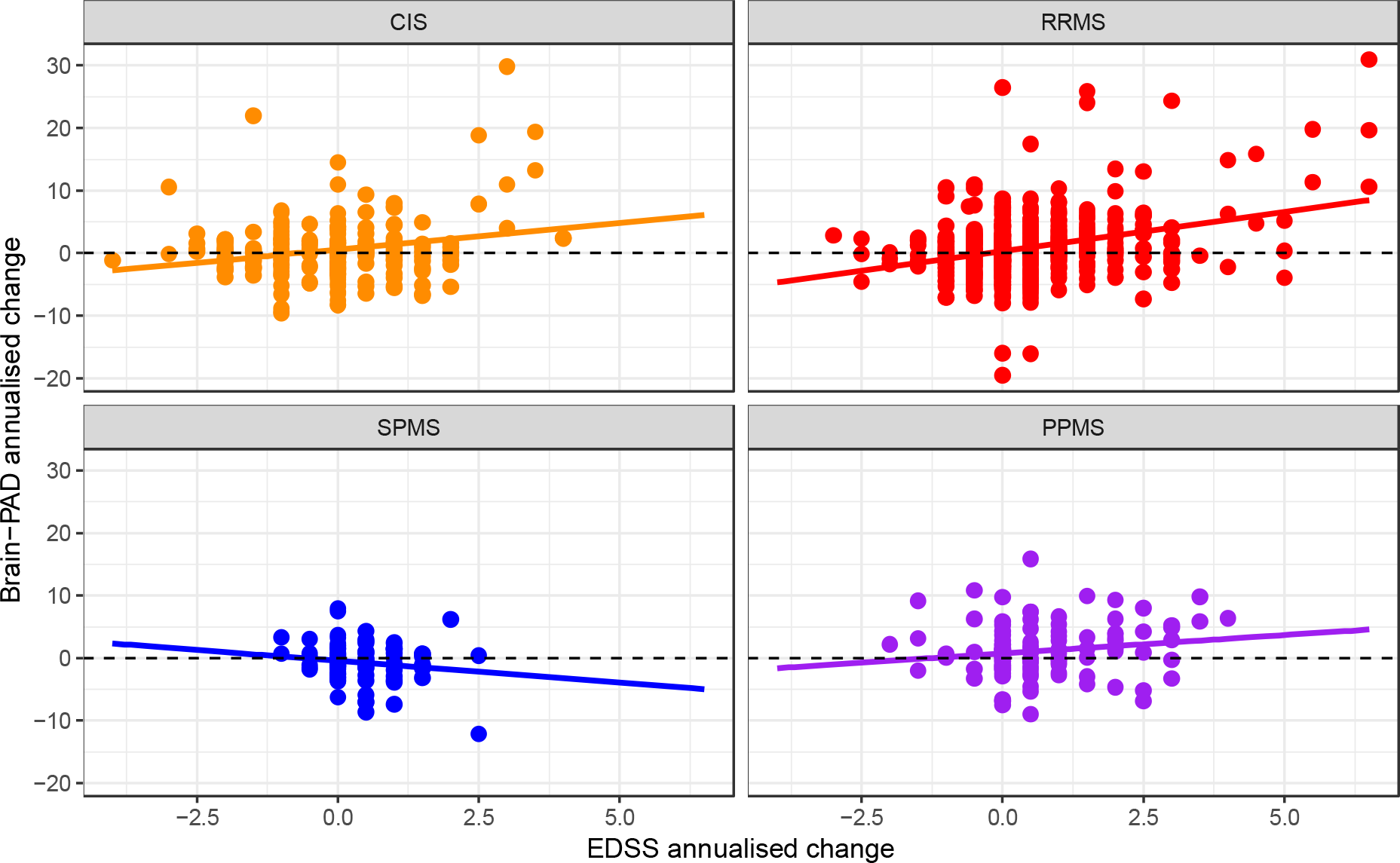
Scatterplot of annualised changed in EDSS score and brain-predicted age difference. Panels show patients with CIS, RRMS, SPMS and PPMS separately, with annualised change in EDSS score between baseline and final follow-up (x-axis) and annualised change in brain-PAD between baseline and final follow-up (y-axis). Lines represented the linear regression lines calculated per group, and shaded areas are the 95% confidence intervals. CIS = clinically isolated syndrome, RRMS = relapsing remitting MS, SPMS = secondary progressive MS, PPMS = primary progressive MS.

### Brain-predicted age difference at first scan predicts EDSS worsening

In patients who had EDSS assessed at ≥2 time-points (n=1147), baseline brain-PAD significantly predicted EDSS worsening. Of these patients, 303 (26⋅5%) experienced EDSS worsening during the follow-up period. Using a Cox proportional-hazards regression model, adjusted for age and sex, the hazard ratio for brain-PAD was 1⋅027 (95% CI 1⋅016–1⋅038, p<0⋅0001). In other words, for every 5 years of additional brain-PAD, there was a 14⋅1% increased chance of EDSS progression during follow-up. Survival curves grouped by a median split of baseline brain-PAD illustrate the differing rates of ‘survival’ prior to EDSS progression (figure 4).

**Figure 4.**
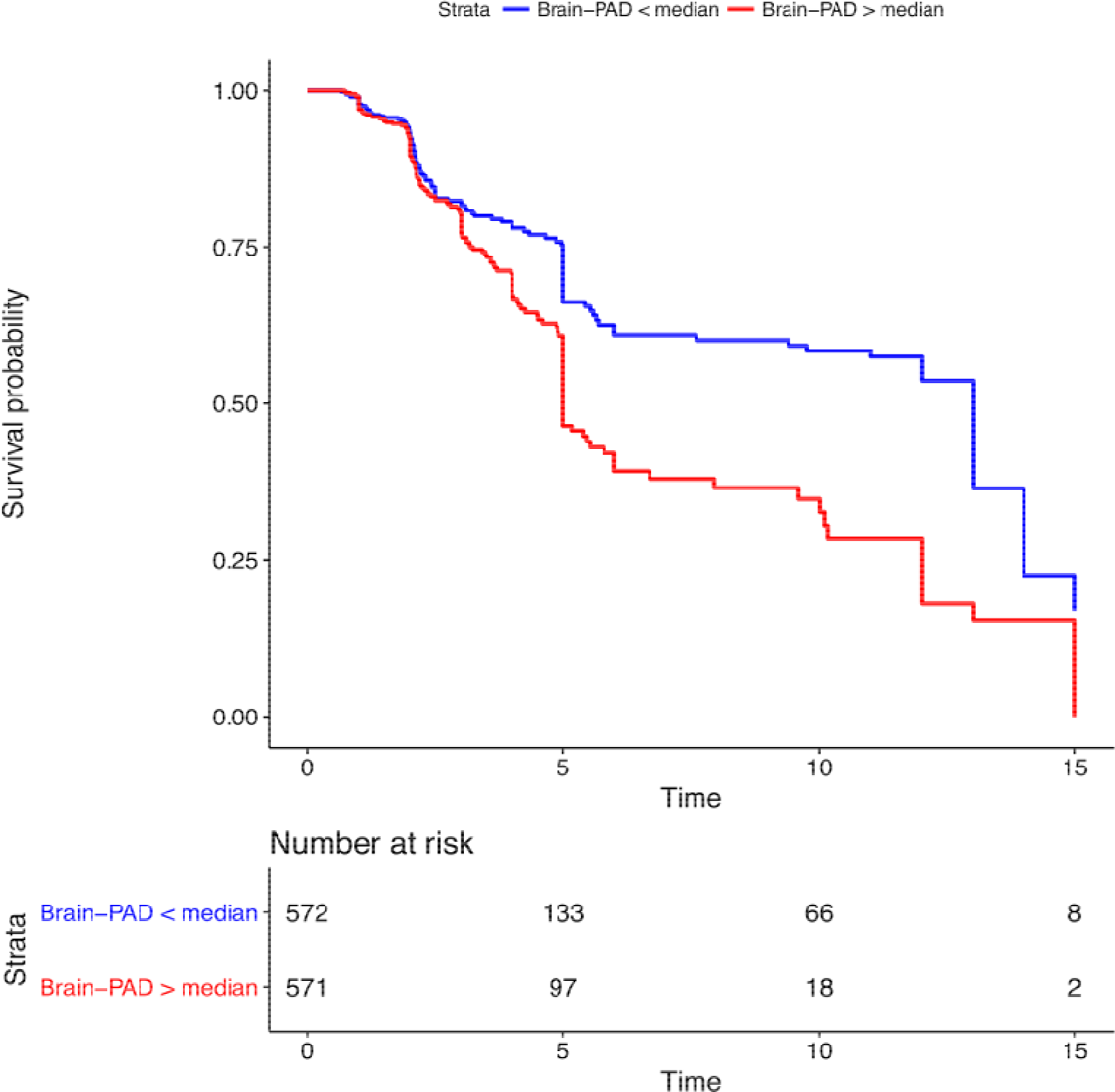
Time-to-EDSS progression survival curves based on baseline brain-PAD. Kaplan-Meier plot illustrating the relationship between brain-PAD at first scan and survival prior to an EDSS progression “event”. Based on a median split of brain-PAD within MS/CIS patients (median brain-PAD = +9⋅68 years).

### MS accelerates longitudinal increase in brain-PAD

A total of 1266 participants had two or more MRI scans (MS/CIS=1155, HCs=111). This included 573 with three or more scans (MS/CIS=509, HCs=64). When using these data, we found a significant interaction between group and time (F_1,1325⋅6_ = 5⋅37, p=0⋅021) and between MS subtypes and time (F_4,938⋅25_ = 5⋅35, p<0⋅0001), when adjusting for age, gender, ICV, cohort and scanner status (figure 5). This indicated that the annual rate of increase in brain-PAD over time was faster in MS/CIS than in HCs, and significantly different between MS subtypes. The estimate marginal mean annualised rates of increase in brain-PAD per group was as follows: HCs −0⋅98 [95% CI −2⋅03–0⋅07], CIS −0⋅14 [95% CI −1⋅07–0⋅78], RRMS 0⋅93 [95% CI 0⋅21–1⋅66], SPMS 0⋅34 [95% CI −0⋅69–1⋅37], PPMS 1⋅21 [95% CI 0⋅16–2⋅25], all CIS/MIS 0⋅70 [95% CI 0⋅01–1⋅39].

**Figure 5.**
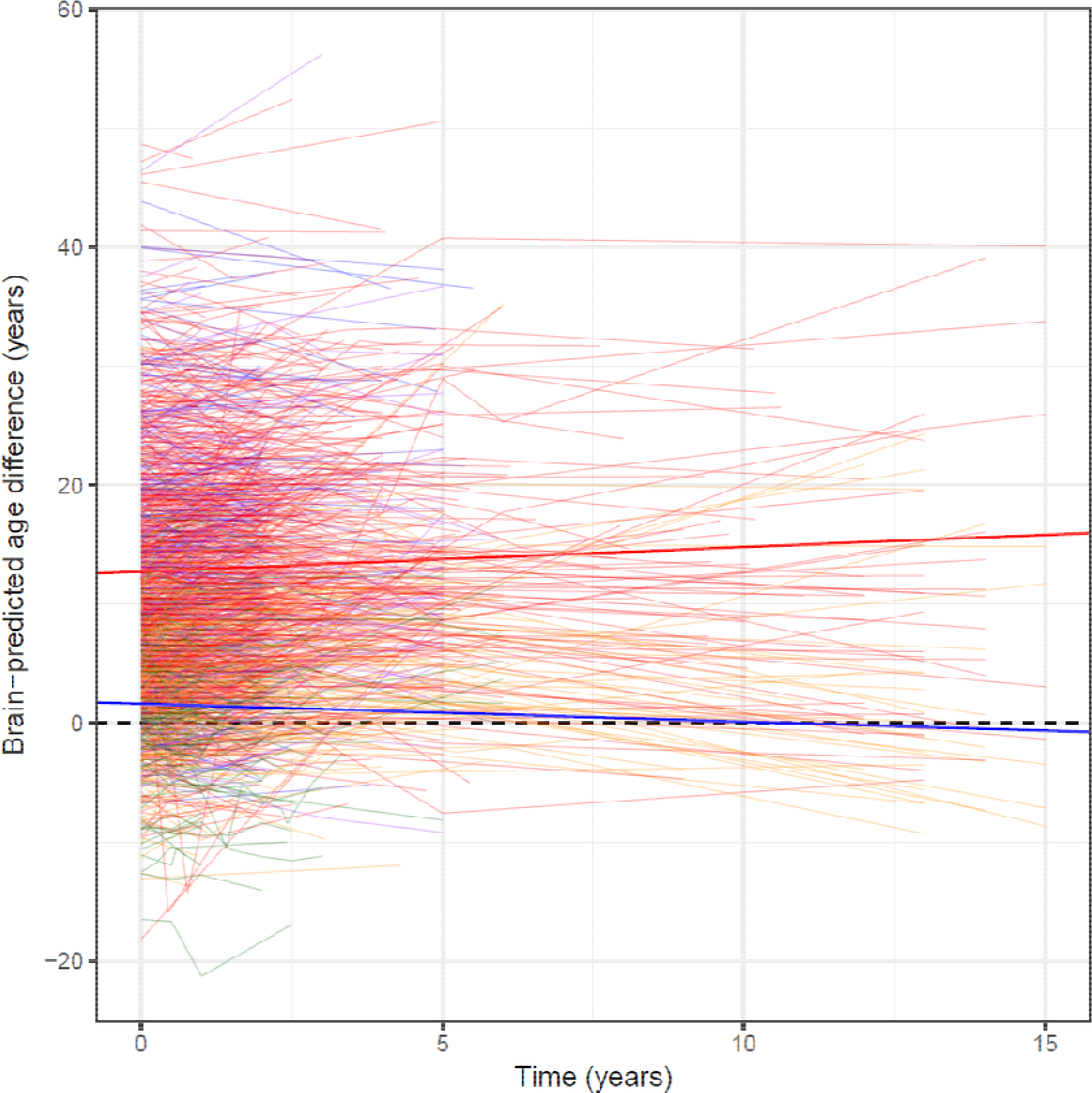
Individual trajectories of brain-predicted age difference by time from baseline. Lines show individual trajectories of brain-PAD scores over the longitudinal study period, coloured according to group (HC = green, CIS = orange, RRMS = red, SPMS = blue, PPMS = purple) Time (from baseline scan) in years (x-axis) and brain-PAD (y-axis). The solid lines represent the average longitudinal slopes for HCs (blue) and all MS/CIS patients (red). The dashed line shows brain-PAD = 0 years.

## Discussion

By assessing the relationship between MS disease progression and the normal brain ageing process, we have found that patients with MS have an older appearing brain (i.e., higher brain-PAD) compared to controls. As the disease develops from a clinically isolated episode to relapsing and then secondary progressive MS, brain-PAD increases. A single baseline brain-PAD was independently associated with higher disability (measured by EDSS), younger age at diagnosis and longer time since diagnosis, irrespectively of disease phenotype. Using scans performed at multiple sites in different scanners we observed that longitudinal brain-PAD increases correlate with worsening disability and that measures of brain-PAD at baseline predict future disability accumulation. In the whole cohort, we show that measures of brain-PAD over time increase with respect to chronological age, implying an accelerated ageing process, particularly in RRMS and PPMS.

In a life-long disease, the accumulation of neurological disability is the main clinical and societal burden,^24^ estimated to cost $10⋅6 billion/year in the USA.^25^ Tracking disease evolution is hampered by the lack of a simple and powerful outcome measure. MRI-assessed brain atrophy is a surrogate outcome for this process, but the need for precise longitudinal assessments, usually over at least a 12-month interval, reduces the feasibility of use. Here, we demonstrate that with a single T1-weighted MRI, brain-PAD values can index elements of MS disease progression. Firstly, we show that a single point estimate can place a patient’s disease and disability in context of their age. This has been lacking with current techniques but is achieved because brain-PAD measures change relative to a model of the healthy ageing process. Our results suggest that the ‘brain-age’ framework can provide informative data without the need for longitudinal scans.^26^ Secondly, we demonstrate that a single measure can give prognostic value for disability accumulation. This can allow us to better contextualise the impact of the disease on an individual, measured at a single time point, and then chart different pathways of neurodegeneration in MS. Brain atrophy has undoubted utility in capturing elements of disease progression, but is currently difficult to utilise in clinical practice.^27^ Here we demonstrate that machine learning technique provides a biomarker of structural brain ageing that enables prediction of disability worsening. Thus, the ability to make prognostic predictions from cross-sectional data could prove highly valuable to facilitate early use of therapy to prevent future disability accumulation.^28^

The brain-age paradigm has been applied widely in neuropsychiatric diseases,^10^ though only recently in MS.^29^ Kaufmann and colleague’s analysis (n=254) showed a strong effect of MS on brain-age (mean increase 5⋅6 years), though was only cross-sectional and did not explore subtypes separately. Here we go further, utilising serial MRI scans that were carried out over 15 years in a wide range of settings – different countries, institutions and scanners. The mean magnitude of the apparent brain ageing we observed MS (11.8 years) is greater than has been reported in dementia (9 years),^11^ epilepsy (4⋅5 years)^17^ or after a traumatic brain injury (4⋅7 years).^14^ We show that brain-PAD increases faster than chronological age in MS/CIS patients, suggesting an accelerating neurodegenerative process. Interestingly, brain-PAD did not increase longitudinally in SPMS patients; potentially due to a survivor bias or a floor effect in this group, whereby those patients with rapidly deteriorating disease did not return for longitudinal follow-up. Evidence for this comes from the inverse correlation between brain-PAD at baseline and the number of follow-up scans acquired in SPMS patients.

We addressed some potential issues with the use of a non-specific ageing biomarker like brain age for the assessment of MS. Brain lesions, the overt MRI marker of MS disease activity, had minimal impact of the brain-PAD measurement in MS. Brain volumes, perhaps unsurprisingly, were strongly correlated with brain-PAD; GM, WM and CSF volume measures combined explained ~49% of the variance in brain-PAD. Evidently, a substantial proportion of variation in brain-PAD is not explained by demographic and MRI characteristics and might be unique to ‘brain-age’. In particular, ‘brain-age’ incorporates voxelwise MRI data in the statistical model, thereby capturing more information than when using summary statistics. This means that more widespread and distributed patterns of features (i.e., voxelwise GM and WM volumes) can contribute to the age-prediction model, capturing elements of cortical thinning, sulcal widening and ventricular enlargement, alongside more macroscopic loss of tissue volume.

Our study has some strengths and weaknesses. The sample size for both training and test sets is relatively large but one potential limitation is the multiple sources of training data, though previous work has shown high between-scanner reliability.^30^ Thus, if it is to be used as a single value this would need to be in the context of individual scanner performance. Comprehensive biomedical data were not available on all these individuals in the training dataset, meaning some may have had undetected health conditions. However, individuals in this sample were screened according to various criteria to ensure the absence of manifest neurological, psychiatric or major medical health issues. We were not able to determine the impact of therapy in this study as it was not a randomised trial and worsening disease drives use of therapy, the effectiveness of which is challenging to determine. However, the majority of the current study sample were on not receiving therapy at baseline, thus therapeutic effects are unlikely to have confounded our results.

This work supports the use of the ‘brain-age’ paradigm in MS. We propose that brain-predicted age has potential value for: 1) MS disease monitoring; potentially capturing the progressive processes that start early on in all disease phenotypes including CIS. 2) Integrating MRI measures of brain injury in MS in a wide range of centres and different scanners. 3) Conveying complex neuroanatomical information in a conceptually simple and intuitive manner. 4) Assessing both current brain health and prognosis. 5) Aiding clinical trial design, by stratifying enrolment based on high brain-PAD, or using brain-PAD as a surrogate outcome measure, reflecting age-associated neurodegeneration. Further work is needed to determine its utility in larger clinical cohorts, but its ease of use makes it an exciting candidate for such cohorts. Further work is needed to improve the anatomical interpretability of brain-age, both in general and specifically to MS. Ultimately, this may offer insight into an individual’s disease course, in line with the move towards precision medicine in the treatment of MS.

## Acknowledgements

JC is funded by a UKRI/MRC Innovation Fellowship. OC, RN, FB and DC acknowledge the National Institute for Health Research University College London Hospitals Biomedical Research Centre. RN acknowledges the National Institute for Health Research Imperial College London Hospitals Biomedical Research Centre. AE received the McDonald Fellowship from Multiple Sclerosis International Federation (MSIF, http://www.msif.org) and ECTRIMS-MAGNIMS fellowship.

---

**Table S1.**
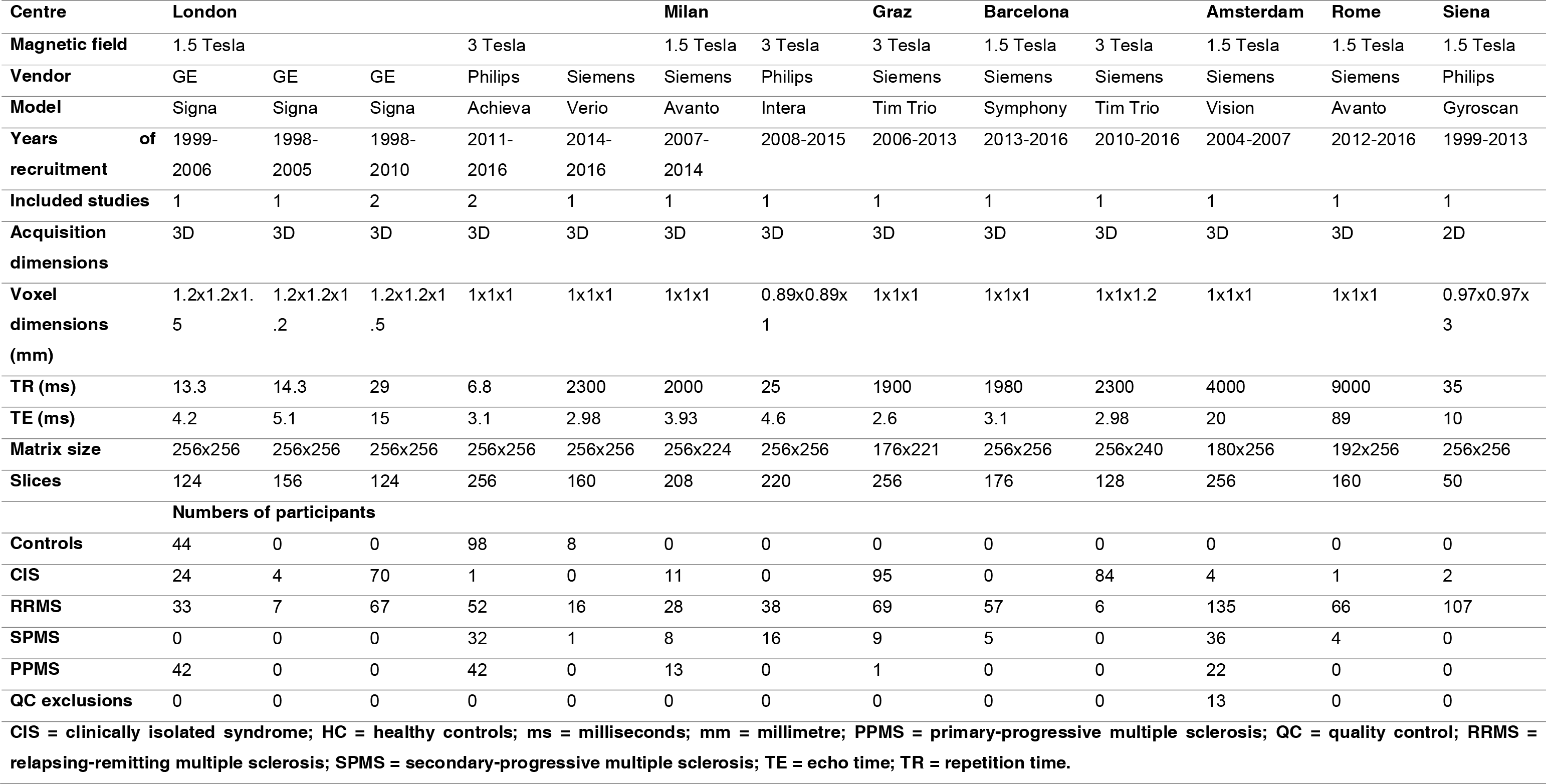
MRI acquisition protocols

**Table S2.**
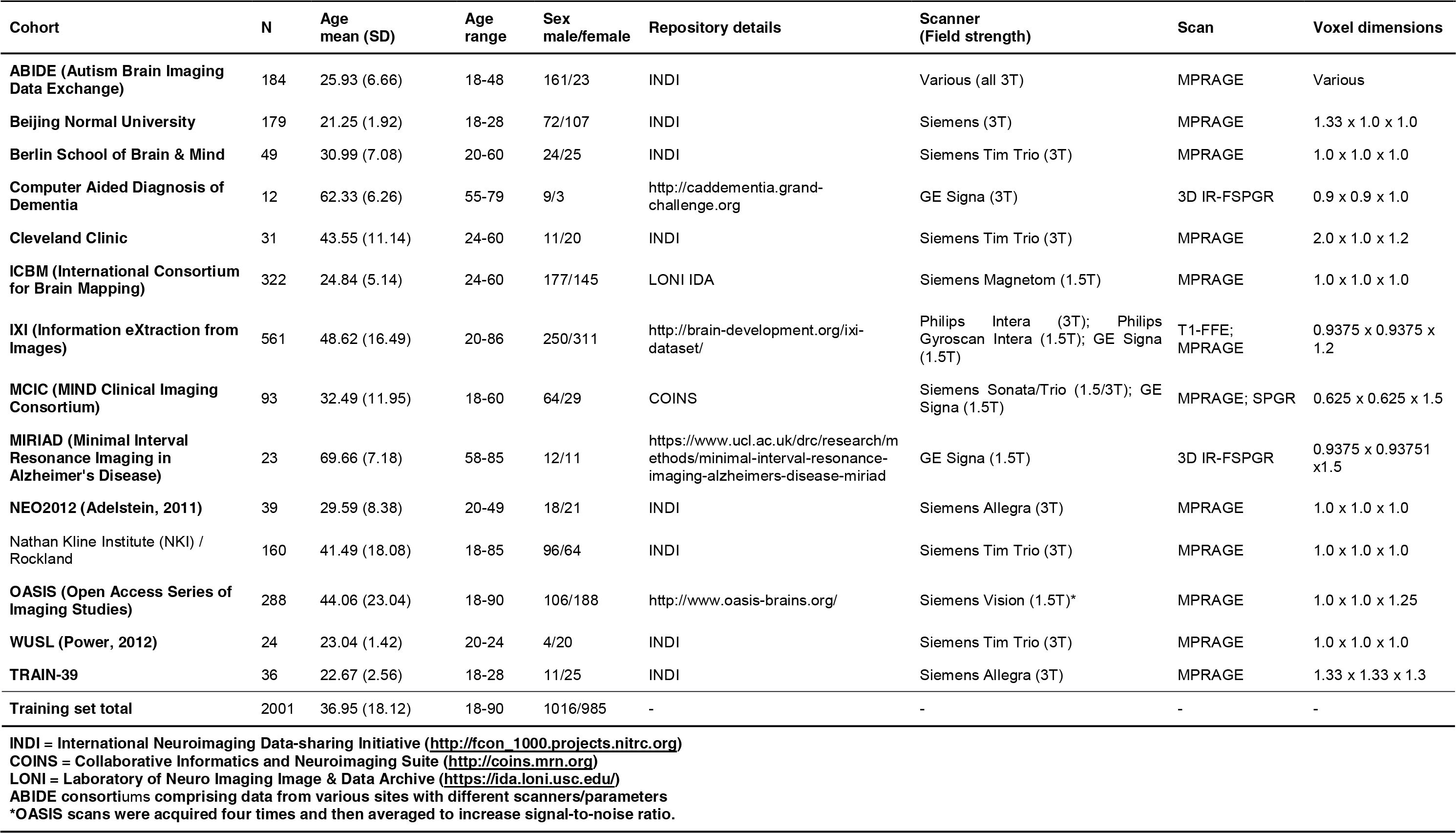
Data sources for healthy brain age training sample

**Table S3.** Post-hoc comparison of differences in brain-PAD between subtypes

| Group comparison | Mean difference<br>(brain-PAD years) | 95% Confidence<br>intervals | P-value |
| --- | --- | --- | --- |
| **CIS-HC** | 3.46 | 0.93, 5.98 | 0.0018 |
| **RRMS-HC** | 11.38 | 9.11, 13.65 | <0.0001 |
| **SPMS-HC** | 16.12 | 12.96, 19.27 | <0.0001 |
| **PPMS-HC** | 10.09 | 7.00, 13.17 | <0.0001 |
| **RRMS-CIS** | 7.92 | 6.17, 9.68 | <0.0001 |
| **SPMS-CIS** | 12.66 | 9.85, 15.46 | <0.0001 |
| **PPMS-CIS** | 6.63 | 3.90, 9.36 | <0.0001 |
| **SPMS-RRMS** | 4.74 | 2.16, 7.32 | <0.0001 |
| **PPMS-RRMS** | -1.29 | -3.79, 1.20 | 0.62 |
| **PPMS-SPMS** | -6.03 | -9.35, -2.71 | <0.0001 |
| CIS = clinically isolated syndrome; HC = healthy controls; PPMS = primary-progressive multiple sclerosis; RRMS = relapsing-remitting multiple sclerosis; SPMS = secondary-progressive multiple sclerosis. |  |  |  |

**Table S4.** Variance explained in brain-predicted age difference by different predictors from hierarchical partitioning of variance.

| Predictor variable | Unique R<sup>2</sup> | Percentage of model variance explained |
| --- | --- | --- |
| **Chronological age** | 0.090 | 18.4% |
| **Sex** | 0.005 | 1.1% |
| **GM volume/ICV** | 0.152 | 31.2% |
| **WM volume/ICV** | 0.046 | 9.4% |
| **CSF volume/ICV** | 0.195 | 39.9% |
| **Total** | 0.488 | 100% |
| CSF = cerebrospinal fluid; GM = grey matter; ICV = Intracranial volume; WM = white matter. |  |  |

**Figure S1.**
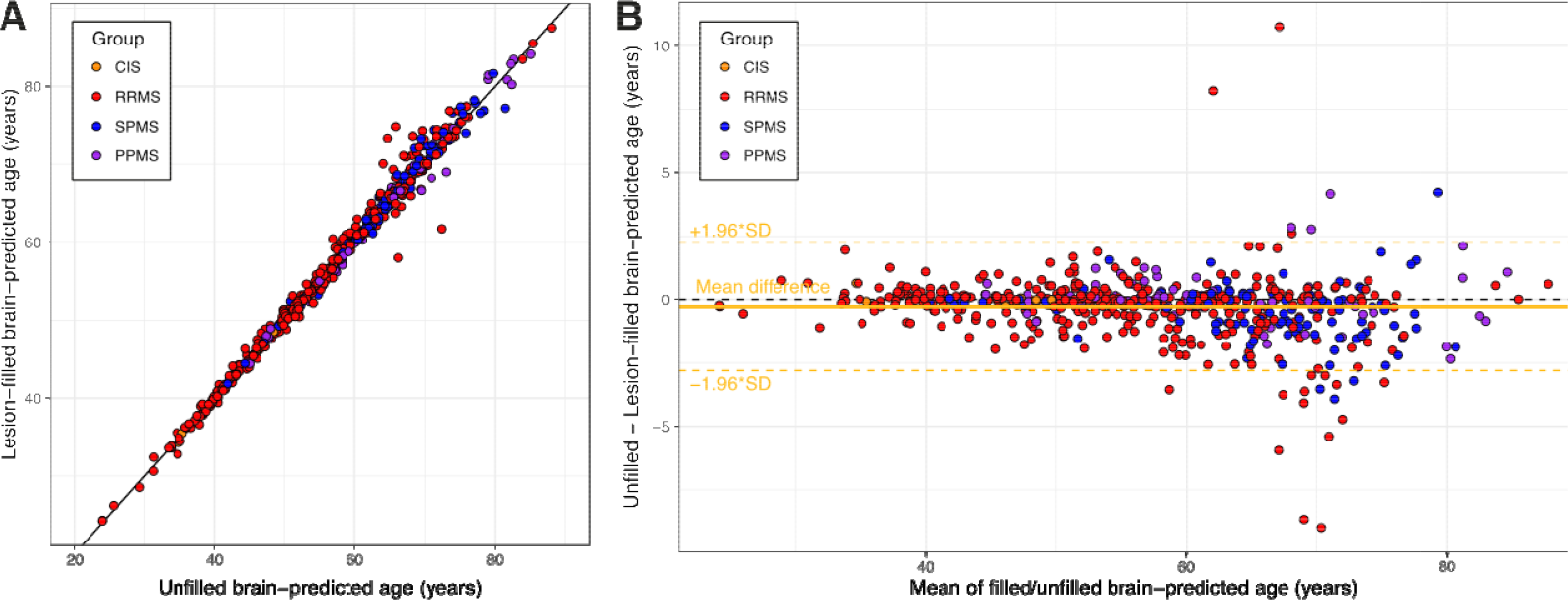
Impact of lesion filling procedure on brain-PAD. A) Scatterplot showing brain-predicted age derived from original, ‘unfilled’ T1-weighted MRI scans (x-axis), plotted against brain-predicted ages generated from T1-weighted MRIs that had undergone the automated lesion ‘filling’ procedure. Points are coloured according to MS subtype or CIS B) Bland-Altman plot of brain-predicted age from unfilled T1-weighted MRI scans and brain-predicted ages generated from ‘filled’ T1-weighted MRIs. The plot shows the mean value from the two measures for each participant (x-axis) and the difference between the two measures (y-axis). The blac dashed line is the line of equality. The mean difference line is plotted in darkgoldenrod1 (mean difference = −0.28 years), along with the limits of agreement (±1.96 * standard deviation of differences, dashed). Total n = 575, with CIS = 8, RRMS = 382, SPMS = 119, PPMS = 66.

## Appendix

### MAGNIMS Study Group: Steering Committee Members

Alex Rovira (co-chair): MR Unit and Section of Neuroradiology, Department of Radiology, Hospital Universitari Vall d’Hebron, Universitat Autònoma de Barcelona, Barcelona, Spain

Christian Enzinger (co-chair): Department of Neurology, Medical University of Graz, Graz, Austria

Frederik Barkhof: Queen Square Multiple Sclerosis Centre, UCL Institute of Neurology, University College London, London, UK

Olga Ciccarelli: Queen Square Multiple Sclerosis Centre, UCL Institute of Neurology, University College London, London, UK

Massimo Filippi: Neuroimaging Research Unit, Institute of Experimental Neurology, Division of Neuroscience, San Raffaele Scientific Institute, Vita-Salute San Raffaele University, Milan, Italy

Nicola De Stefano: Department of Medicine, Surgery and Neuroscience, University of Siena, Siena, Italy

Ludwig Kappos: Department of Neurology, University Hospital, Kantonsspital, Basel, Switzerland

Jette Frederiksen: The MS Clinic, Department of Neurology, University of Copenhagen, Glostrup Hospital, Denmark

Jaqueline Palace: Centre for Functional Magnetic Resonance Imaging of the Brain, University of Oxford, UK

Maria A Rocca: Neuroimaging Research Unit, Institute of Experimental Neurology, Division of Neuroscience, San Raffaele Scientific Institute, Vita-Salute San Raffaele University, Milan, Italy

Jaume Sastre-Garriga: Department of Neurology/Neuroimmunology, Multiple Sclerosis Centre of Catalonia (CEMCAT), Hospital Universitari Vall d’Hebron, Universitat Autònoma de Barcelona, Barcelona, Spain

Hugo Vrenken: Department of Radiology and Nuclear Medicine, MS Center Amsterdam, Amsterdam, The Netherlands

Tarek Yousry: NMR Research Unit, Institute of Neurology, University College London, London, UK

Claudio Gasperini: Department of Neurology and Psychiatry, University of Rome Sapienza, Rome, Italy.

### R Notebook used for statistical analysis

James Cole - March 2019. Built with R version 3.5.2

This is Notebook contains the final brain age analysis of MS patient data and controls from the UCL cohort, the MAGNIMS consortium and the Imperial College London PET study (n=25). The analysis uses brain-predicted age difference (brain-PAD) to look at brain ageing in the context of MS. The brain-PAD values were generated in PRONTO, using an independent healthy (n=2001) training dataset, and the values were corrected for proportional bias using the intercept and slope of the age by brain-predicted age regression in the training dataset.

#### Initial set up of analysis

Clear workspace, load libraries, set colour palette

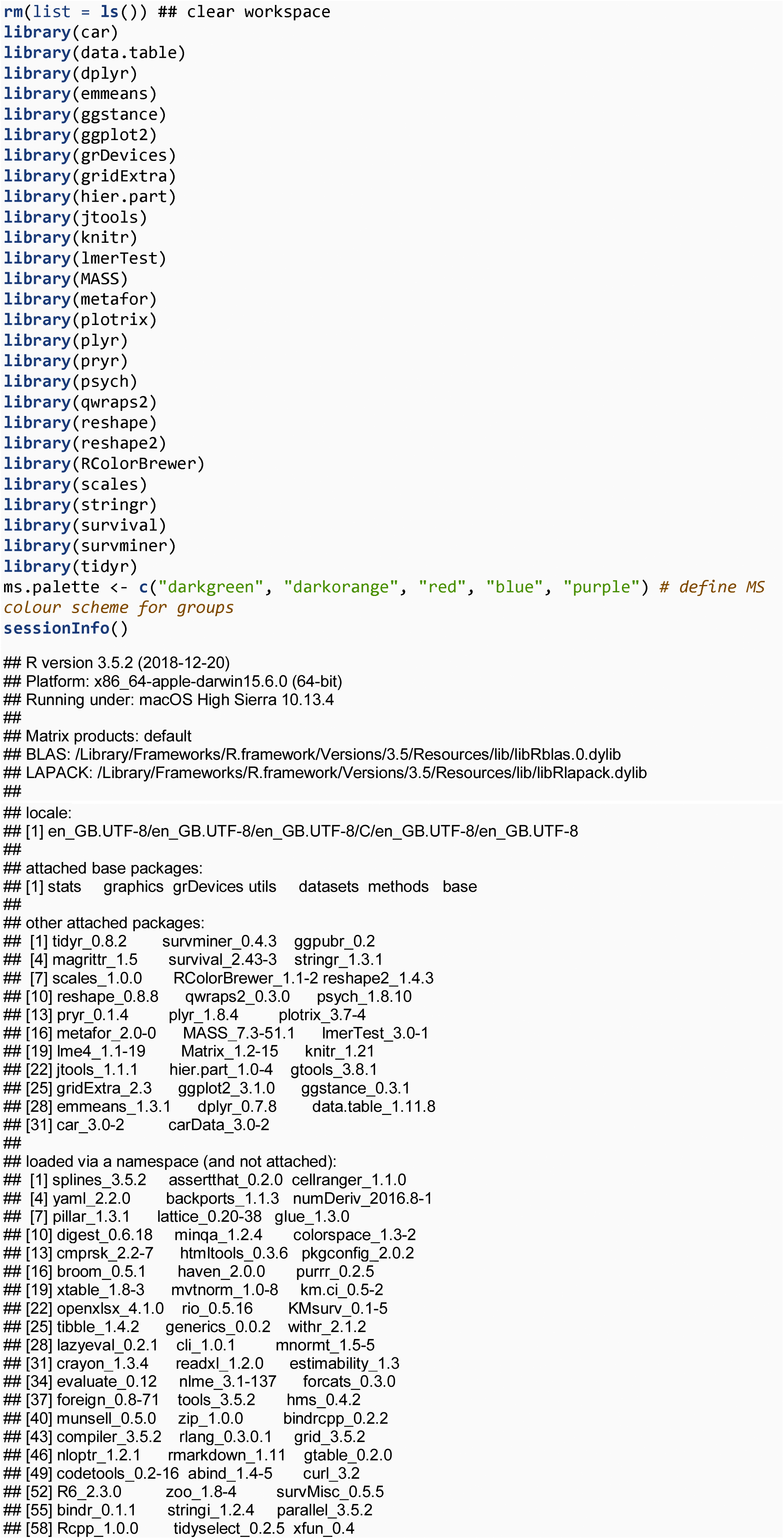

**Get data from CSV and define longitudinal data.frames**

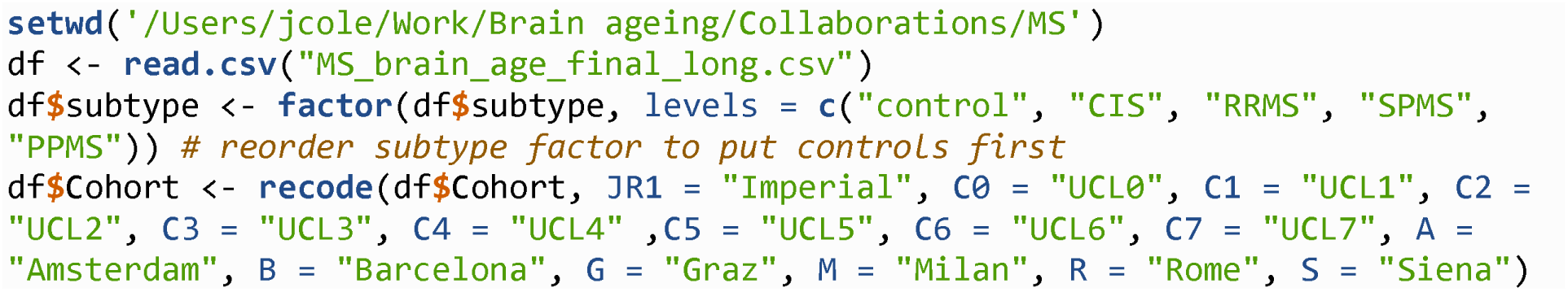

**Exclude participants with errors in the database & correct time since diagnosis errors**

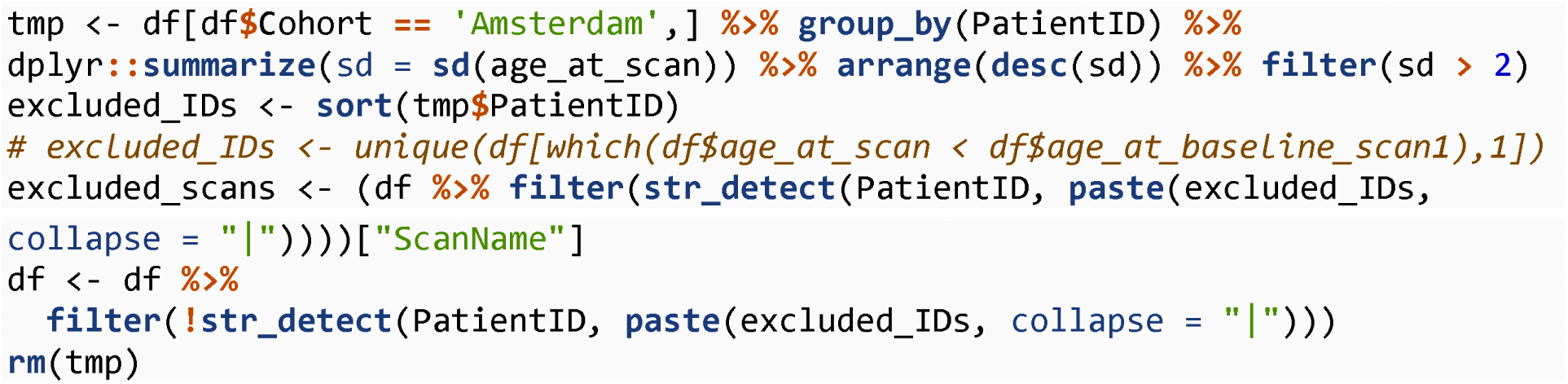

There were 13 subjects with 38 scans excluded in total. Data entry errors in original spreadsheet; age at baseline not consistent within subject.

Load data for lesion filling analysis

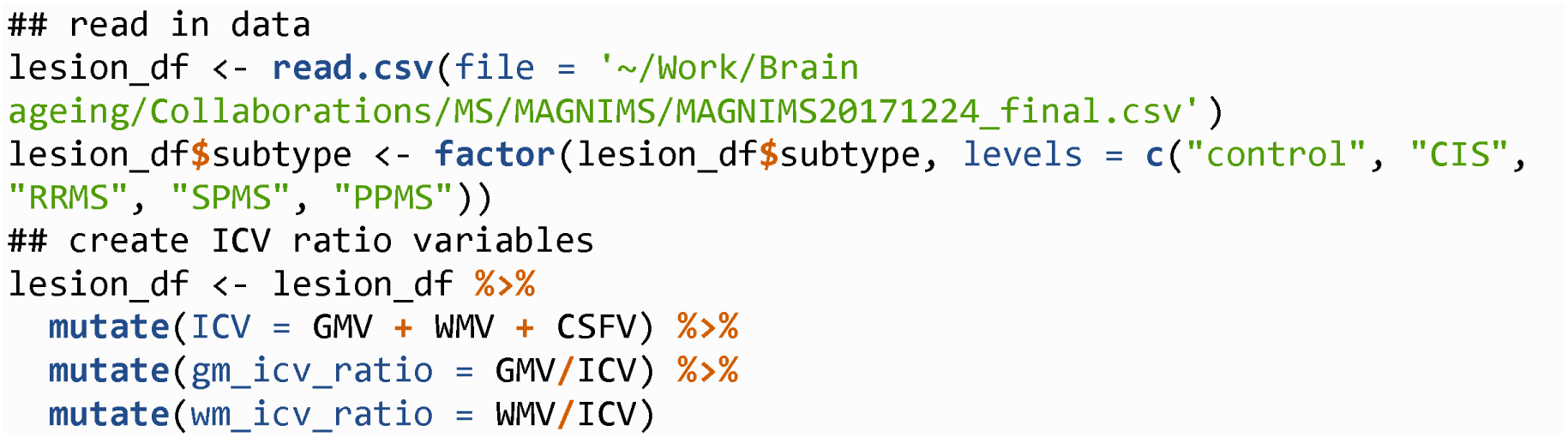

**Generate baseline only data.frame and show data frame structure**

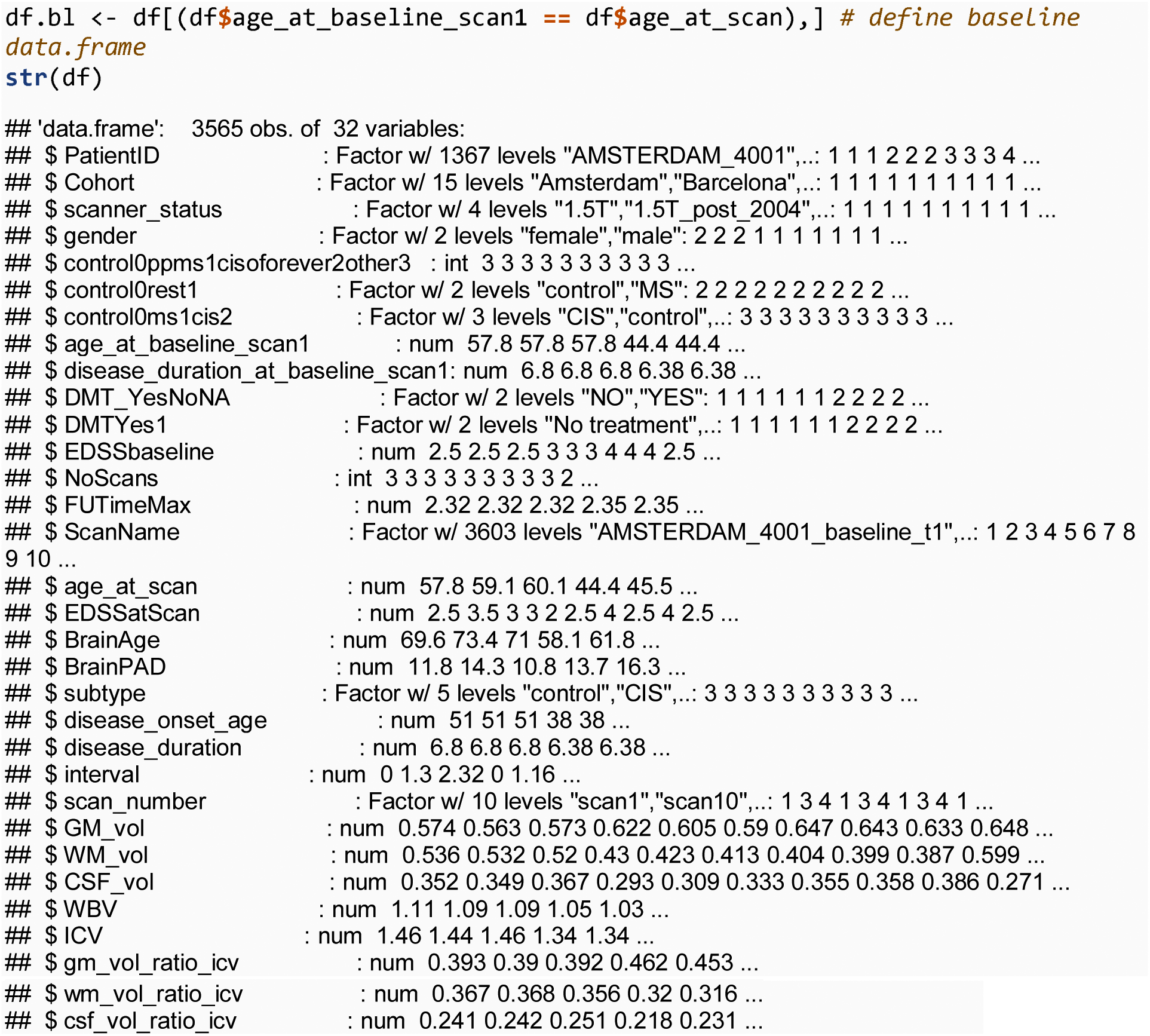

#### Basic stats

Total number of subjects, and by group

The total number of subjects included was n = 1354

The total number of MS patients (including CIS) was n = 1204 and healthy controls was n = 150

**Number of scans in total and per group**

Total number of scans = 3565

**Number of people with 2 or more scans**

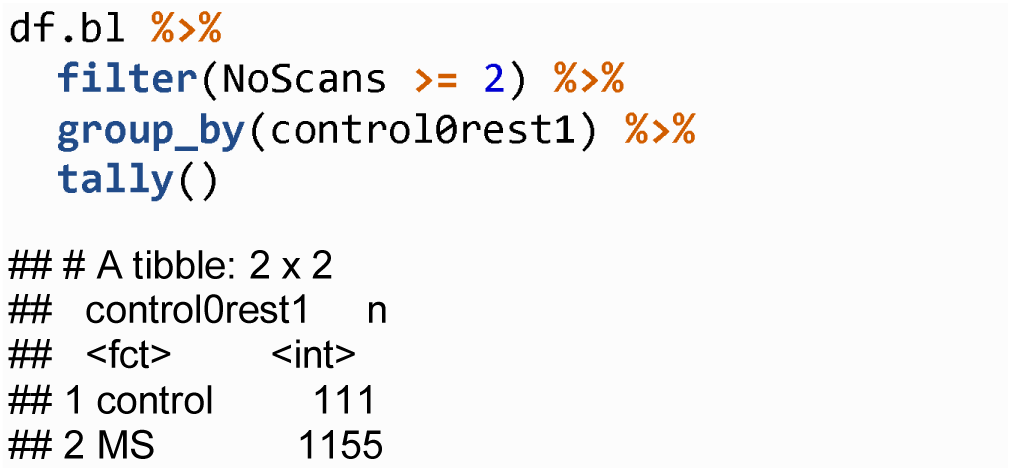

Number of people with 3 or more scans

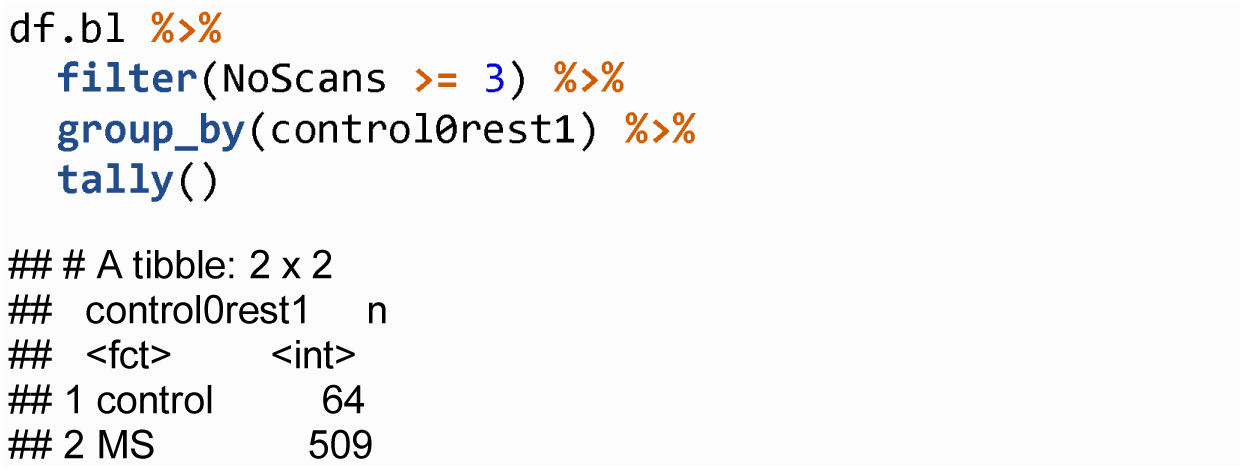

Generate demographics table using qwarps2

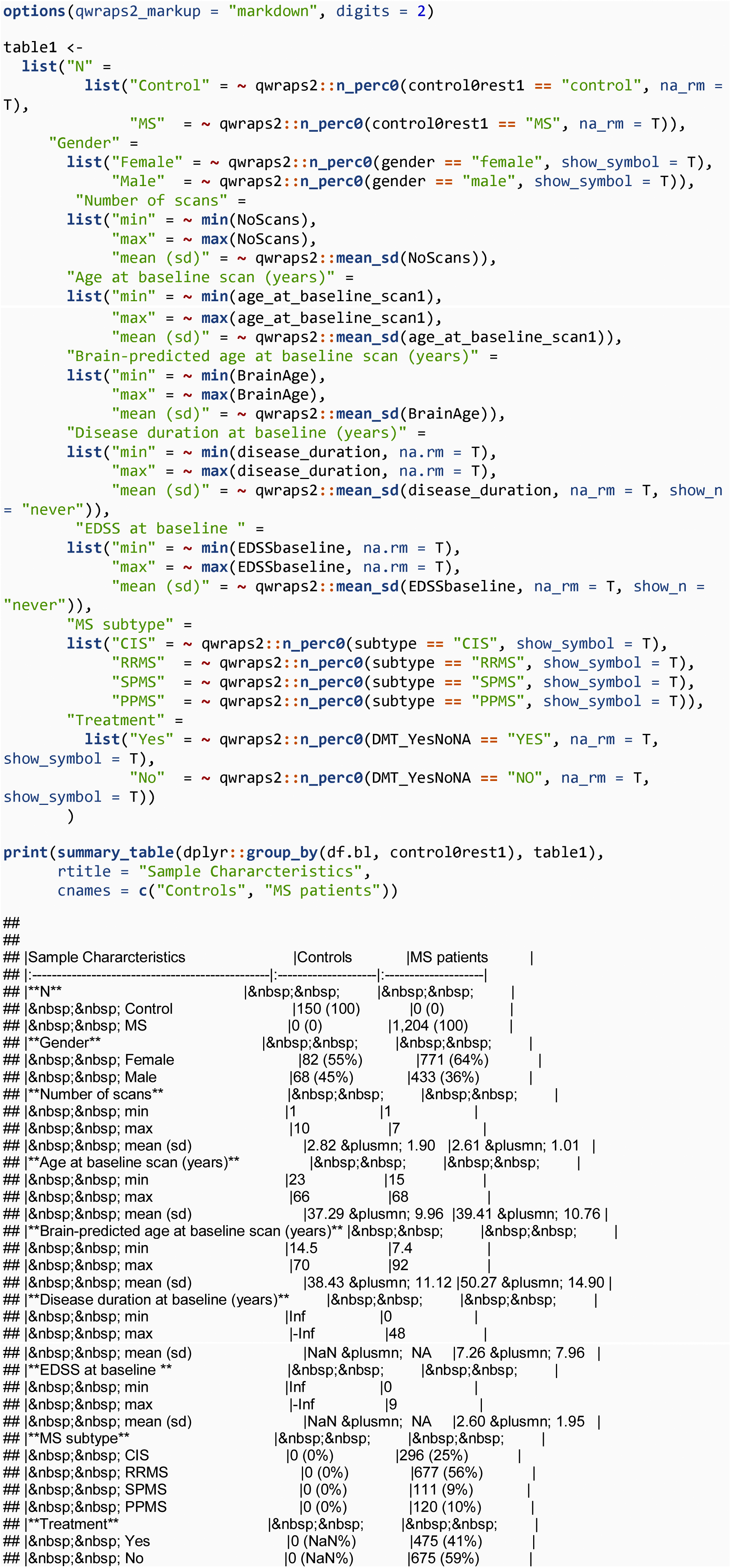

**Need to get treatment NAs using table()**

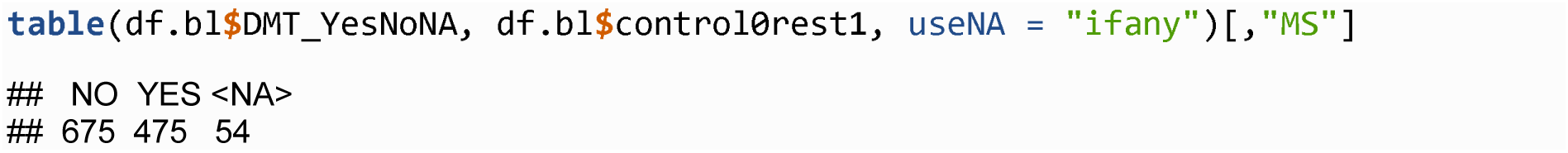

Need to get length of follow-up from longitudinal database

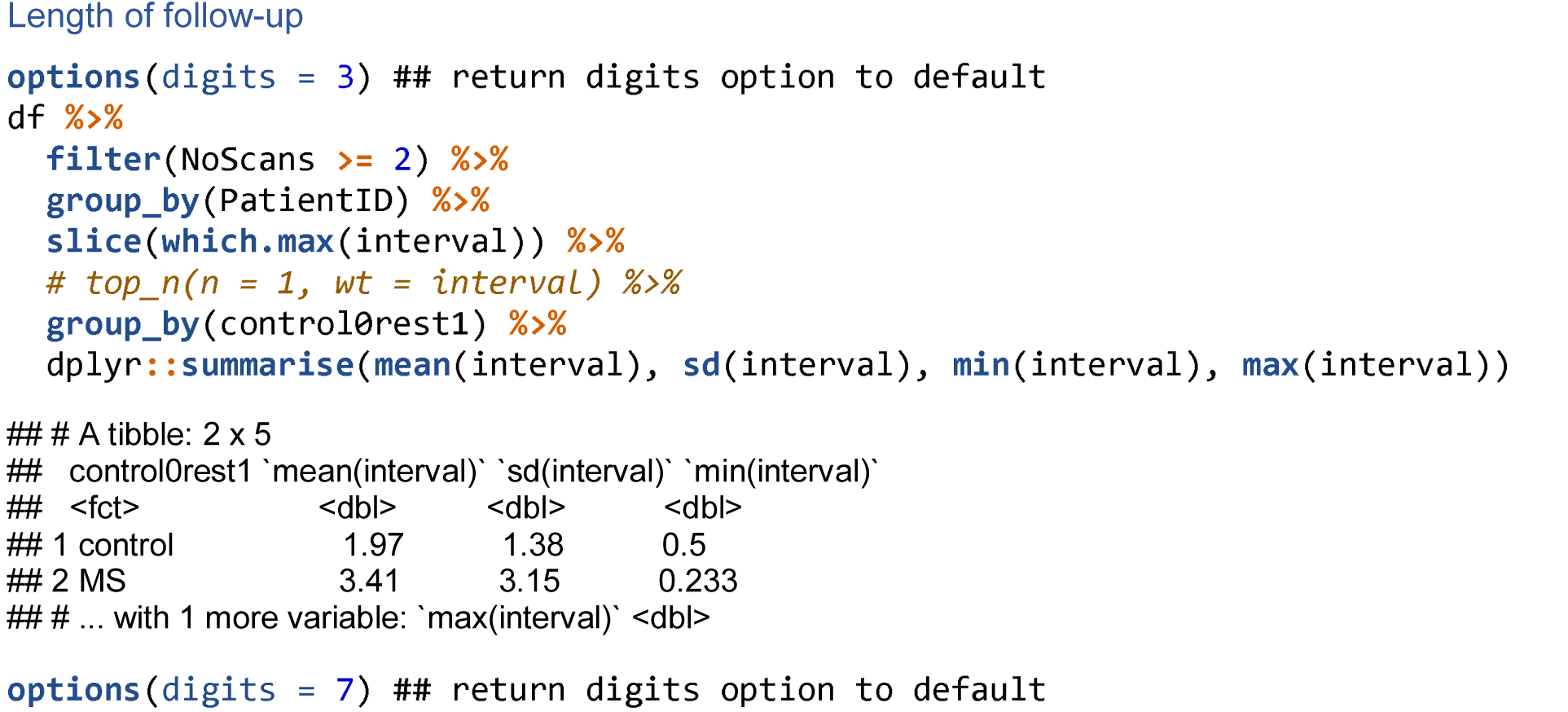

#### Baseline brain-age analysis

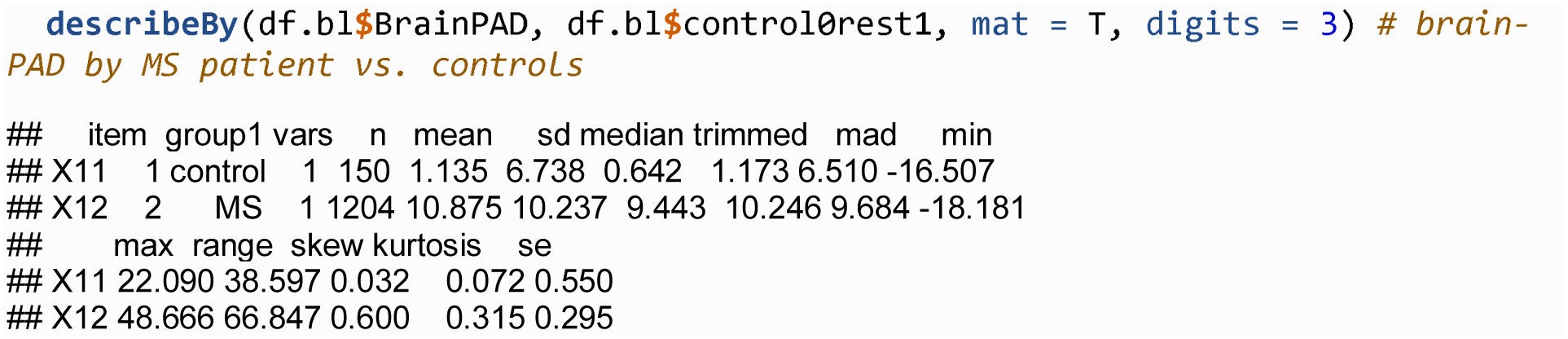

**Estimated marginal means**

Generate EMMs for all MS/CIS and healthy controls. LME adjusting for age, gender, ICV, cohort and scanner status.

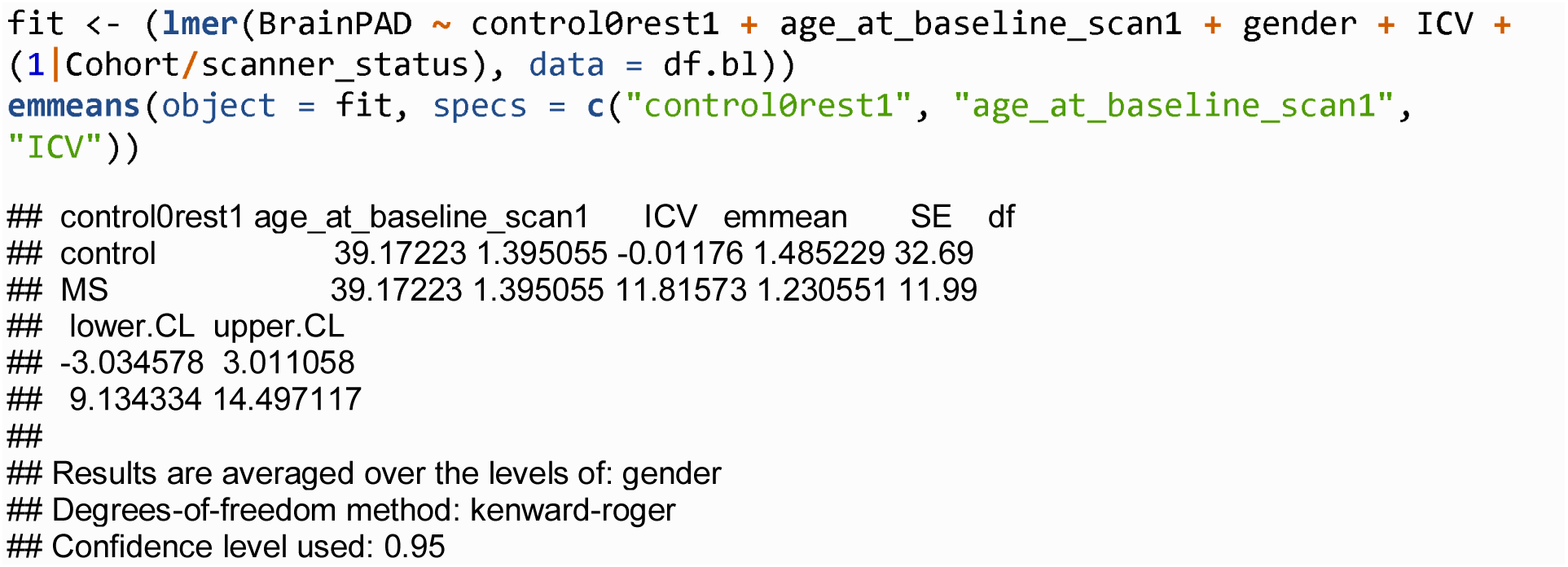

**Effects of cohort and scanner status on brain-PAD.**

The significant interaction motivates nesting in the LME.

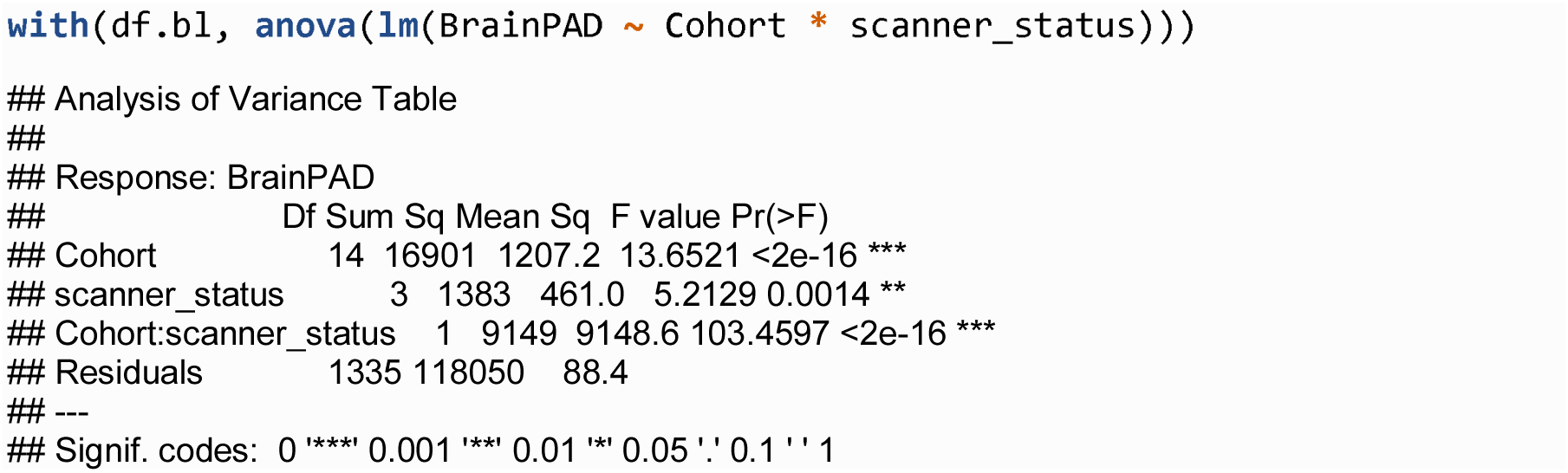

**Meta-analysis looking at all the separate cohorts with MS/CIS patients and controls**

Check which cohorts contain healthy controls and patients.

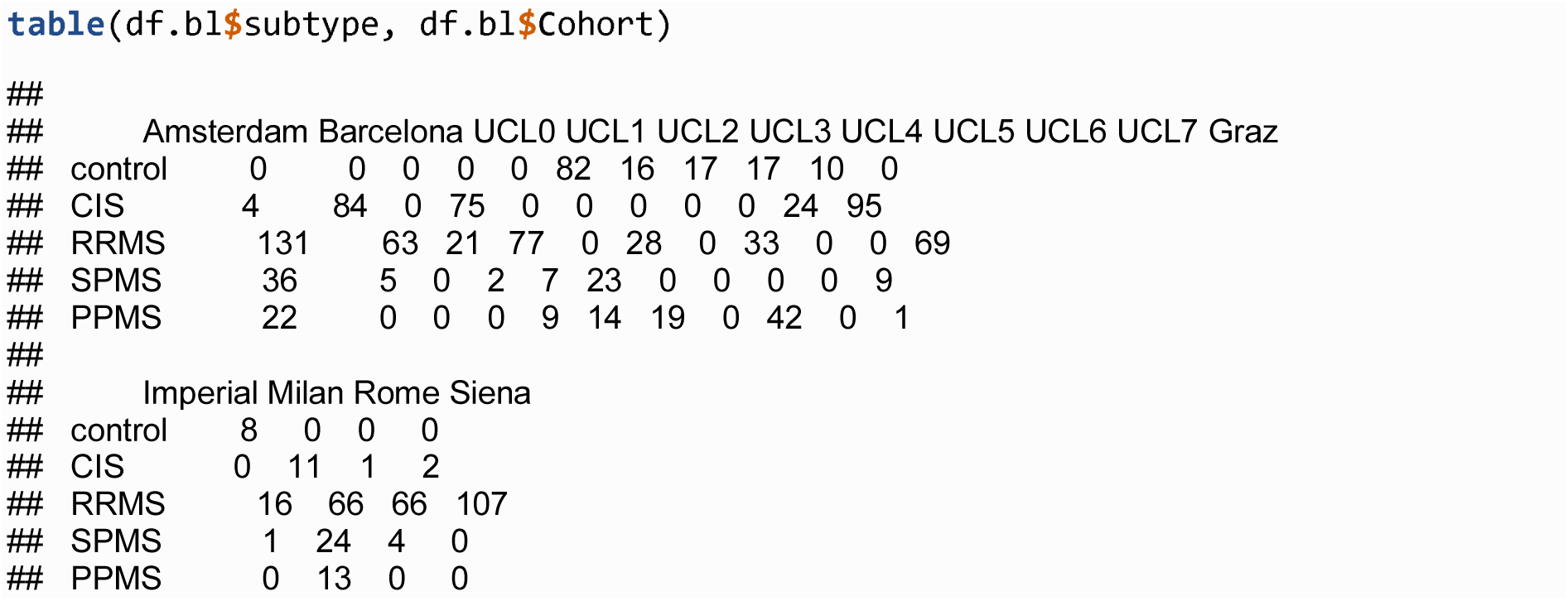

Create data.frame with summary data appropriate for meta-analysis.

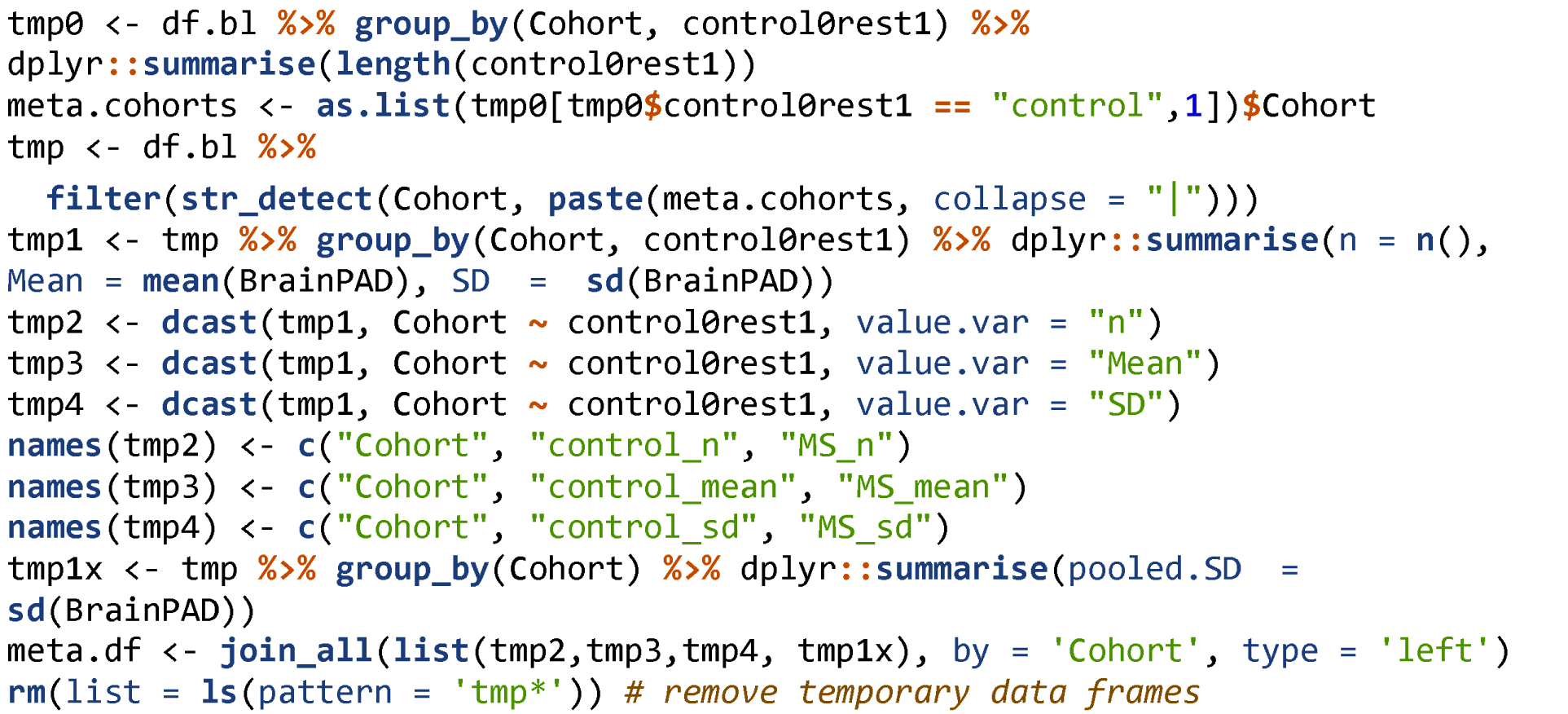

Run meta-analysis using the metafor package, to fit a random-effects meta-analysis using REML.

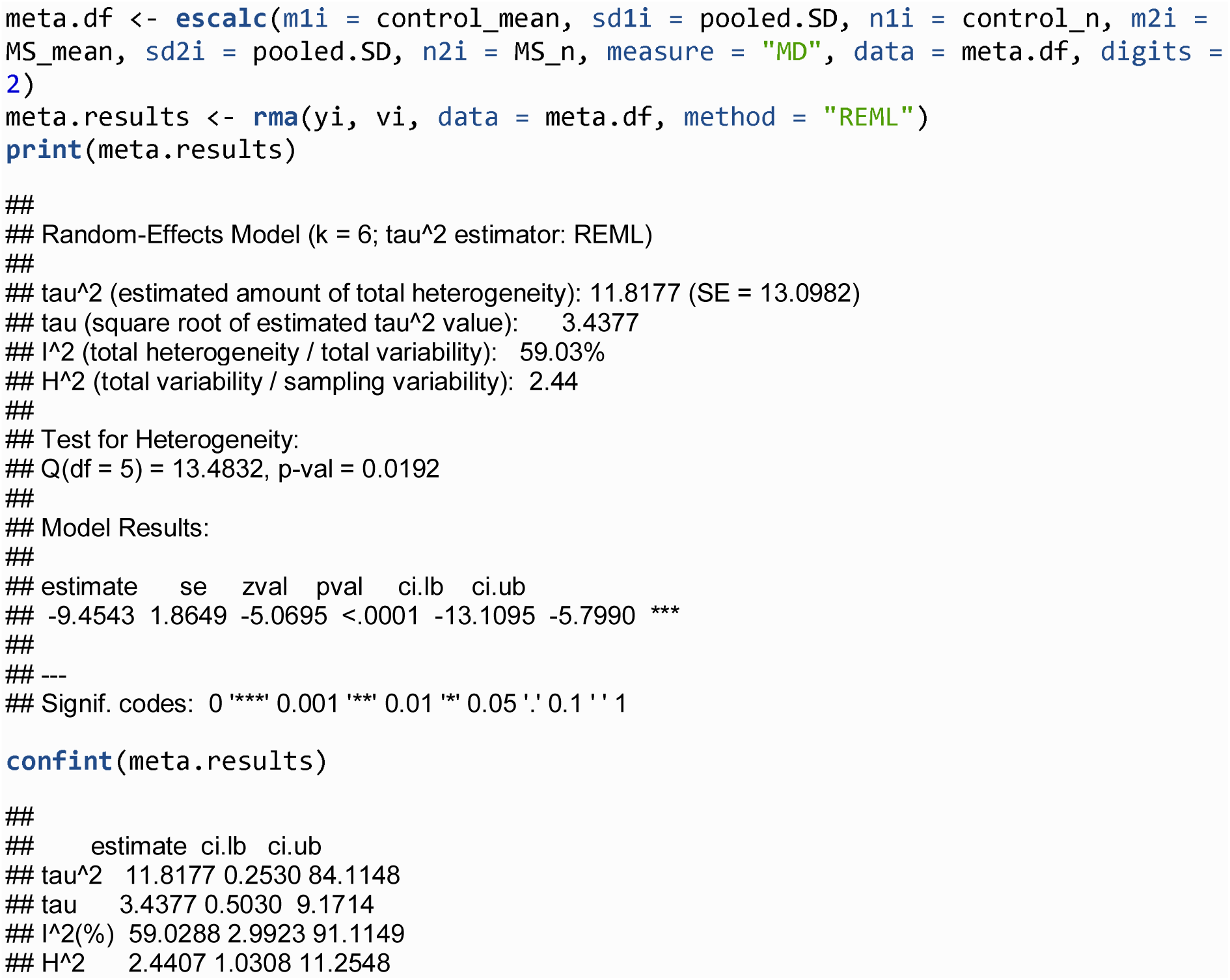

Forest plot of results

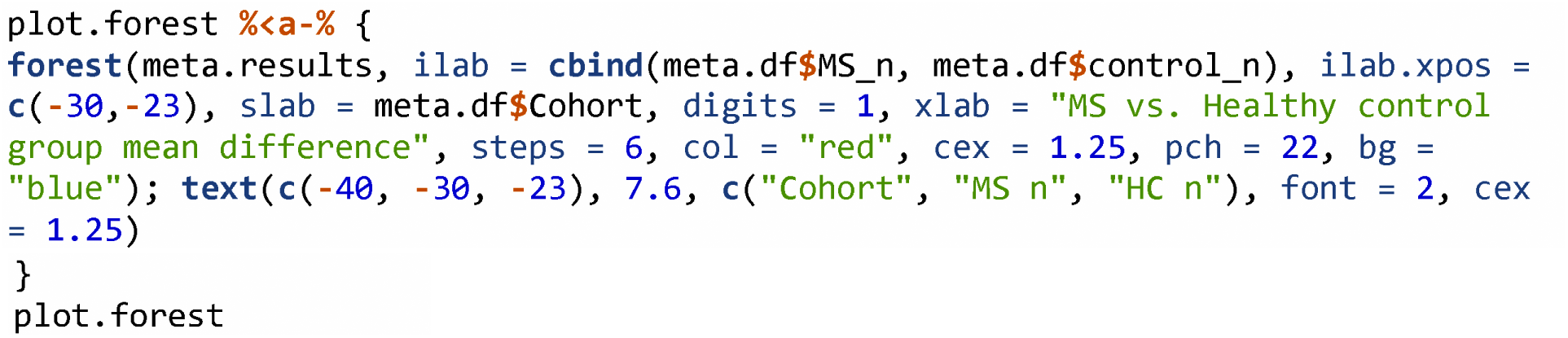

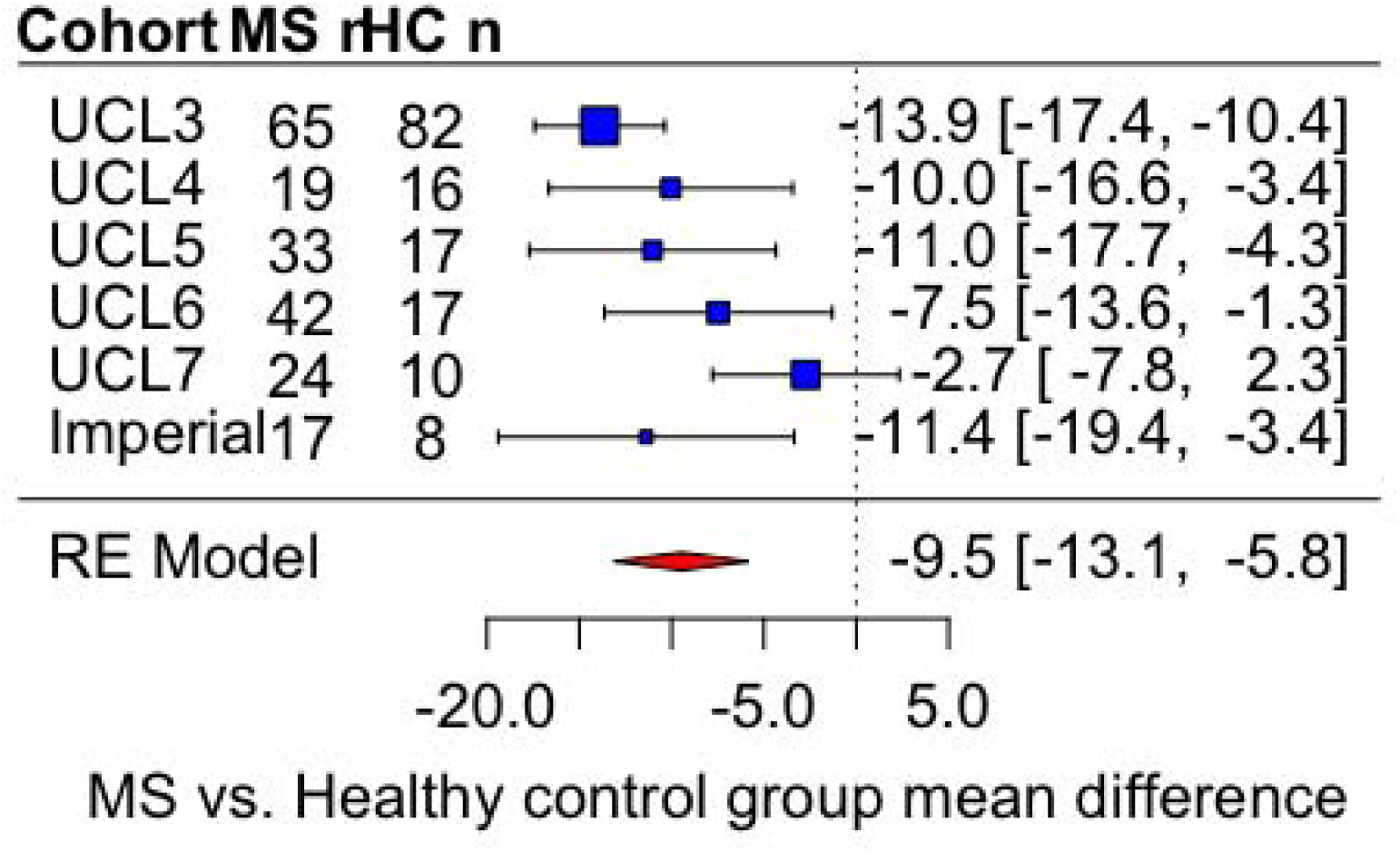

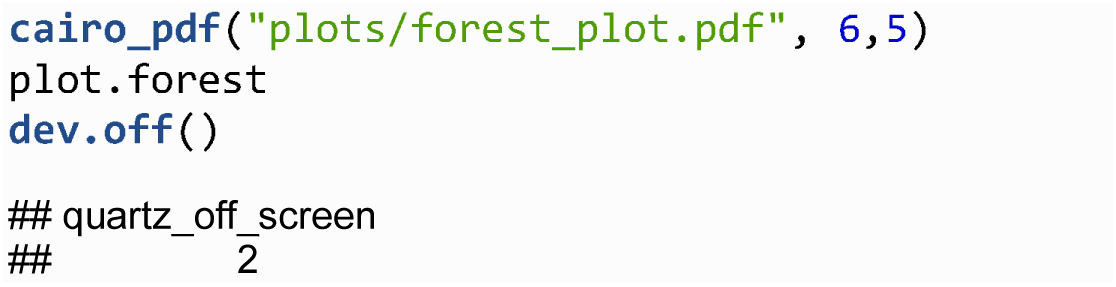

Linear regression analysis restricted to cohort UCL3, includes covariates: age, gender, intracranial volume.

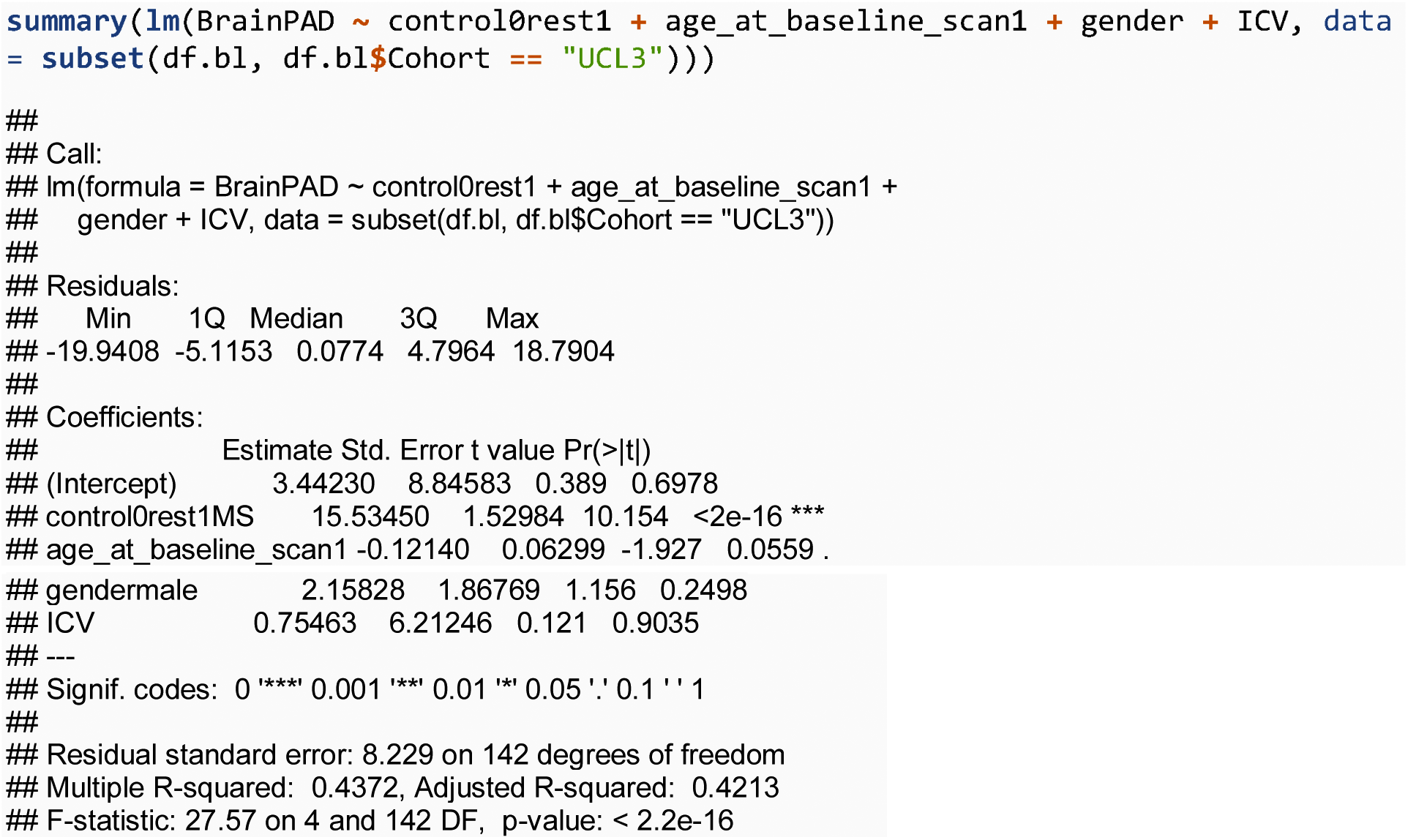

### Main result

#### LME model to predict brain-PAD based on group

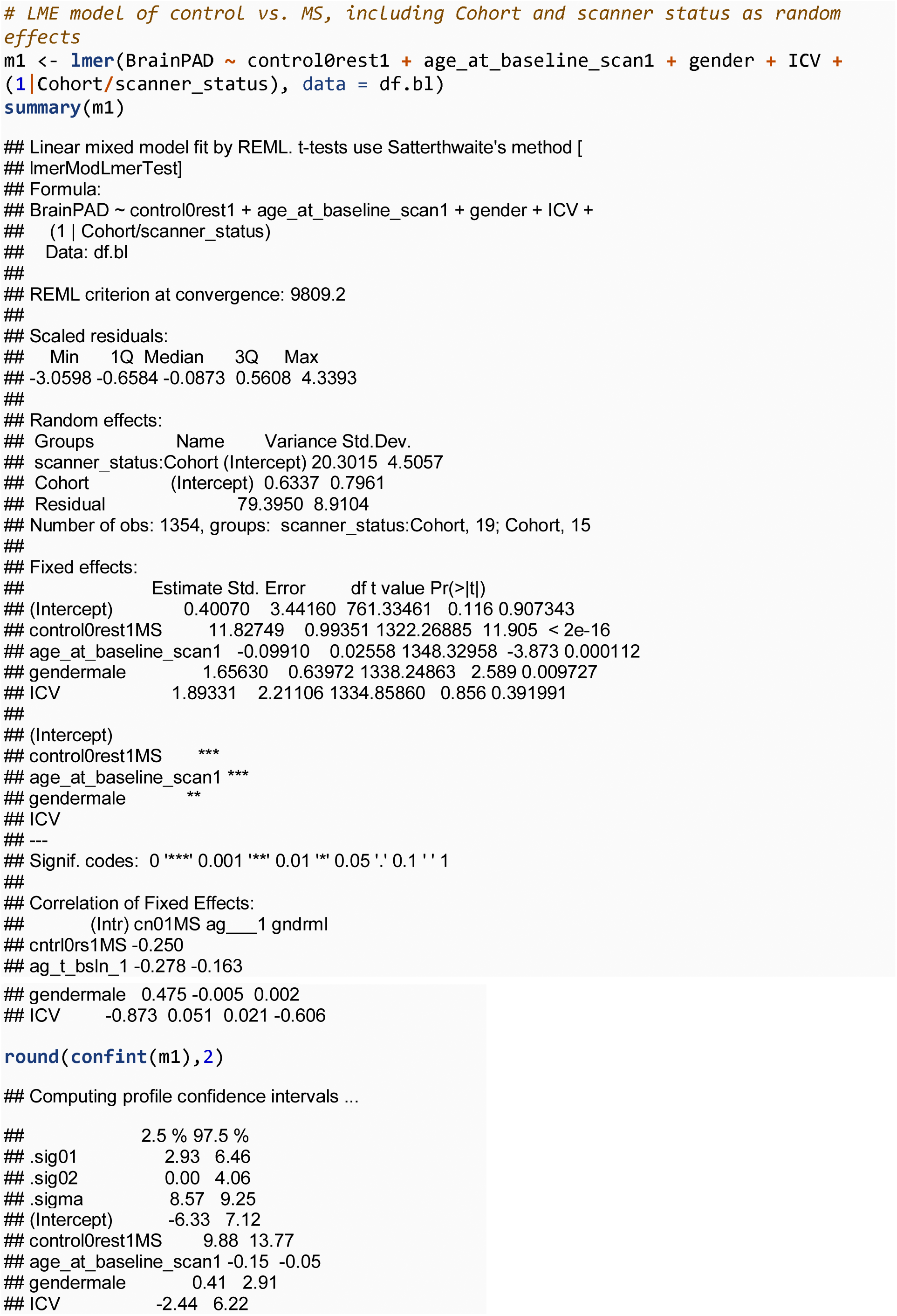

#### Lesion filling

To establish whether using the FSL lesion filling software influences brain-predicted age values. This analysis was conducted only in UCL patients.

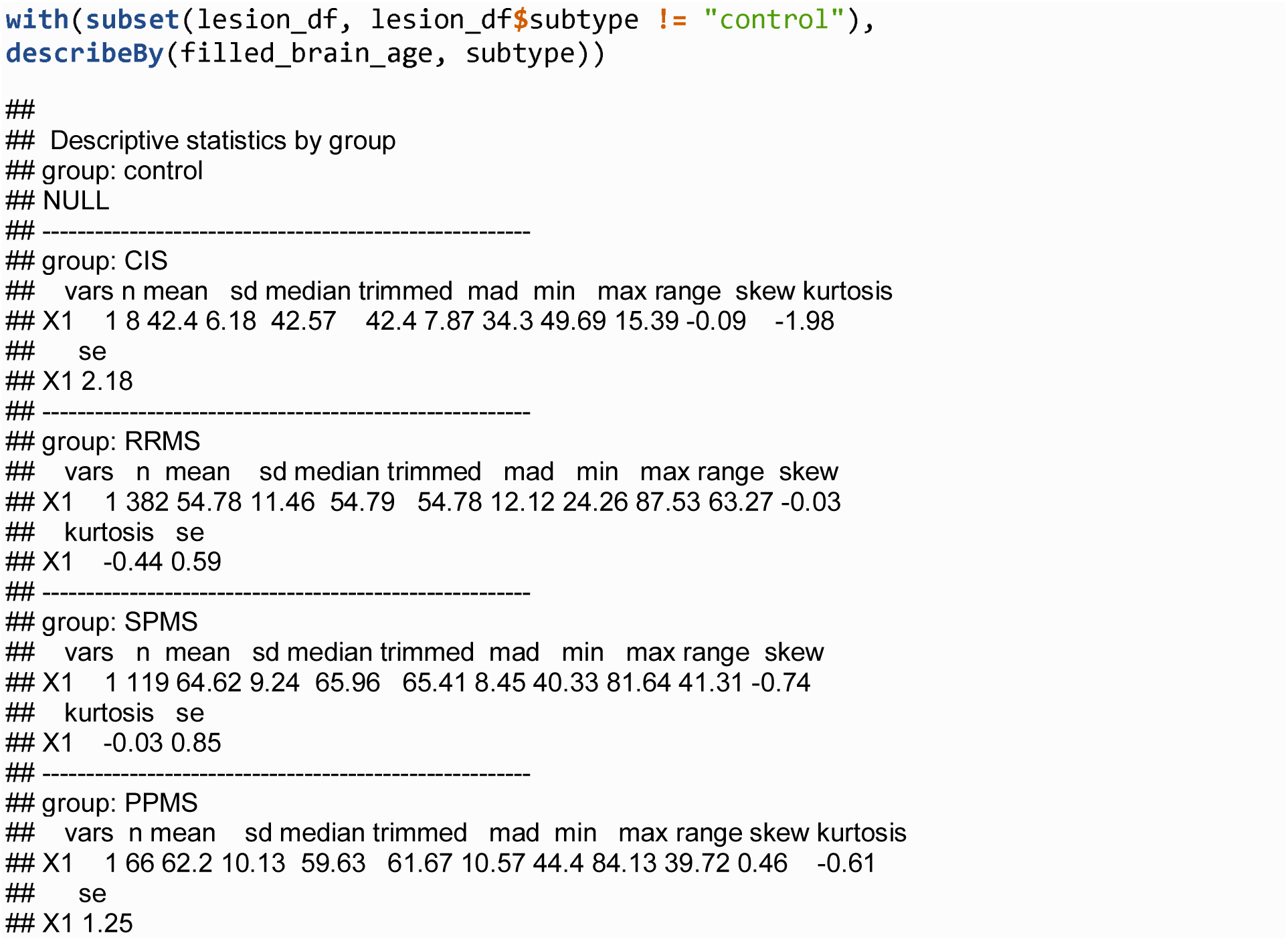

Correlation between brain-predicted age from filled and unfilled images: Pearson’s r = 0.994. Median absolute error (MAE) between brain-predicted age from filled and unfilled images = 0.3717 years. Mean difference between brain-predicted age from filled and unfilled images = 0.28 years.

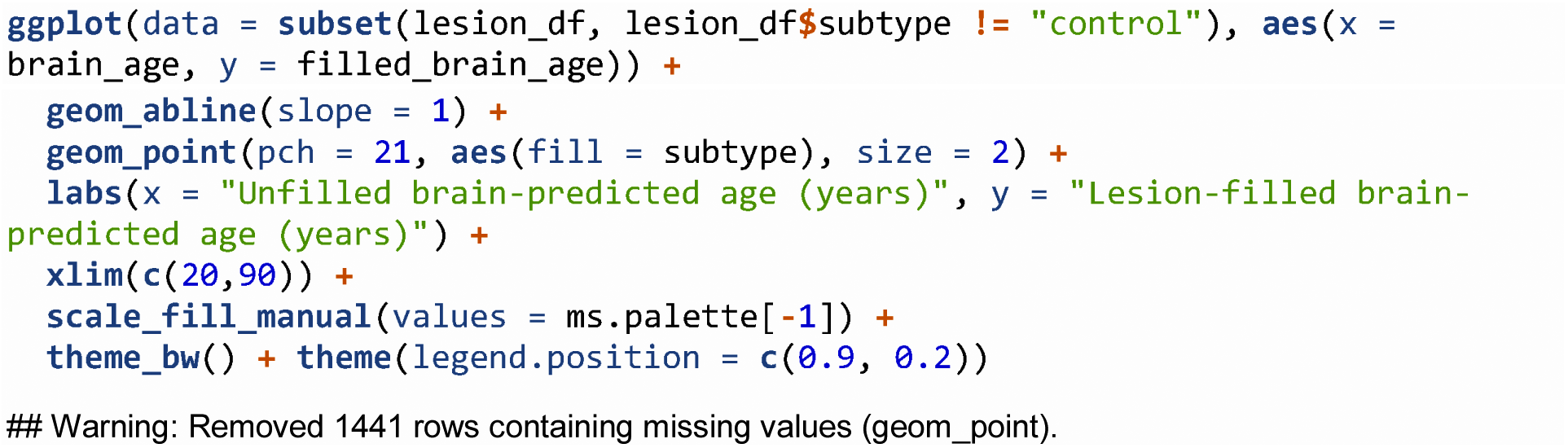

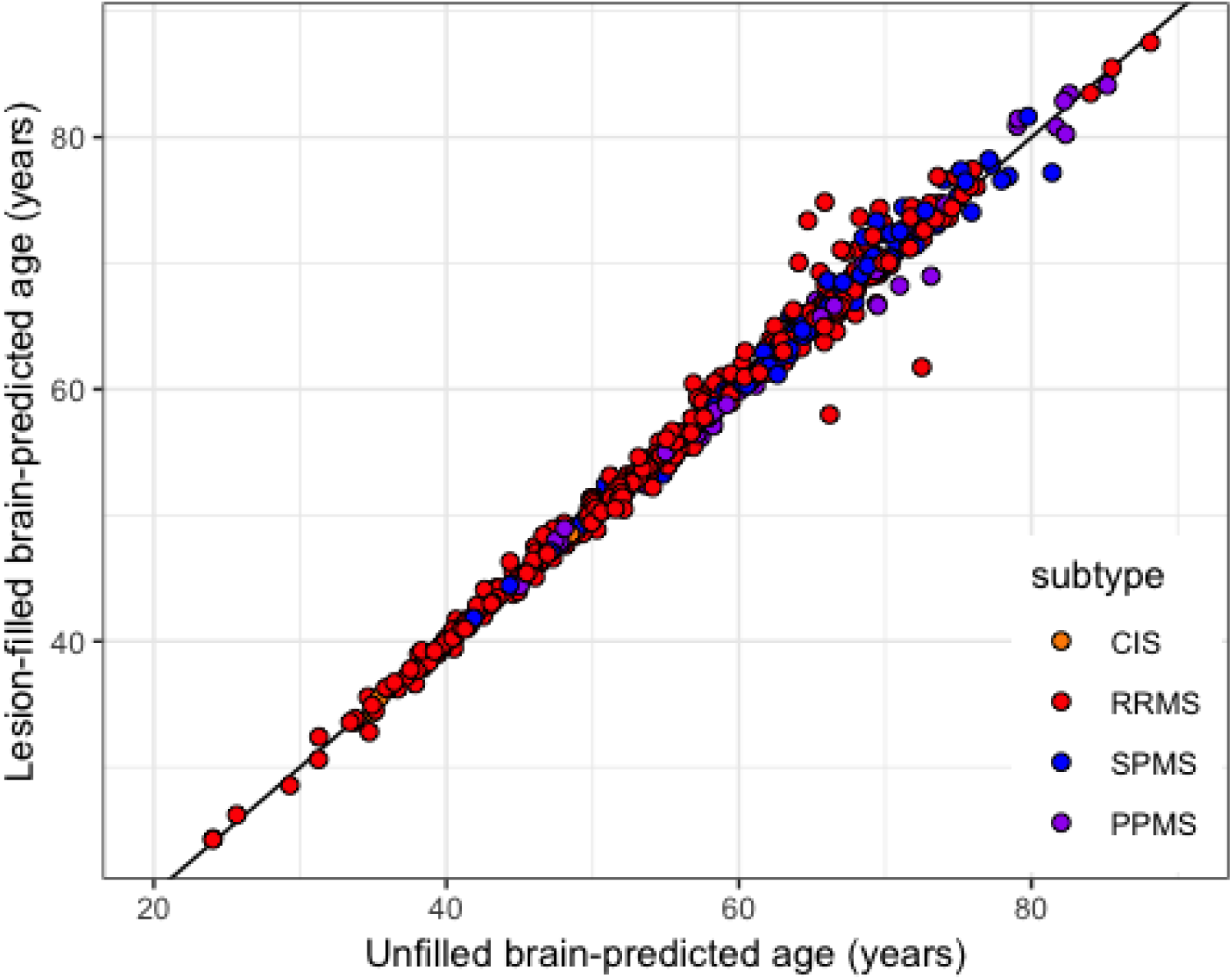

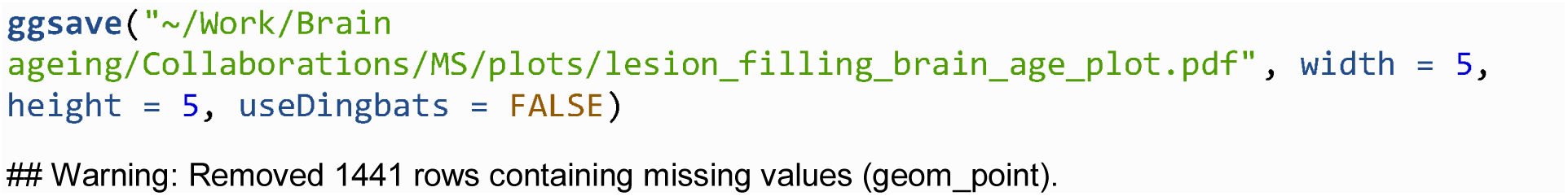

#### Bland-Altman plot

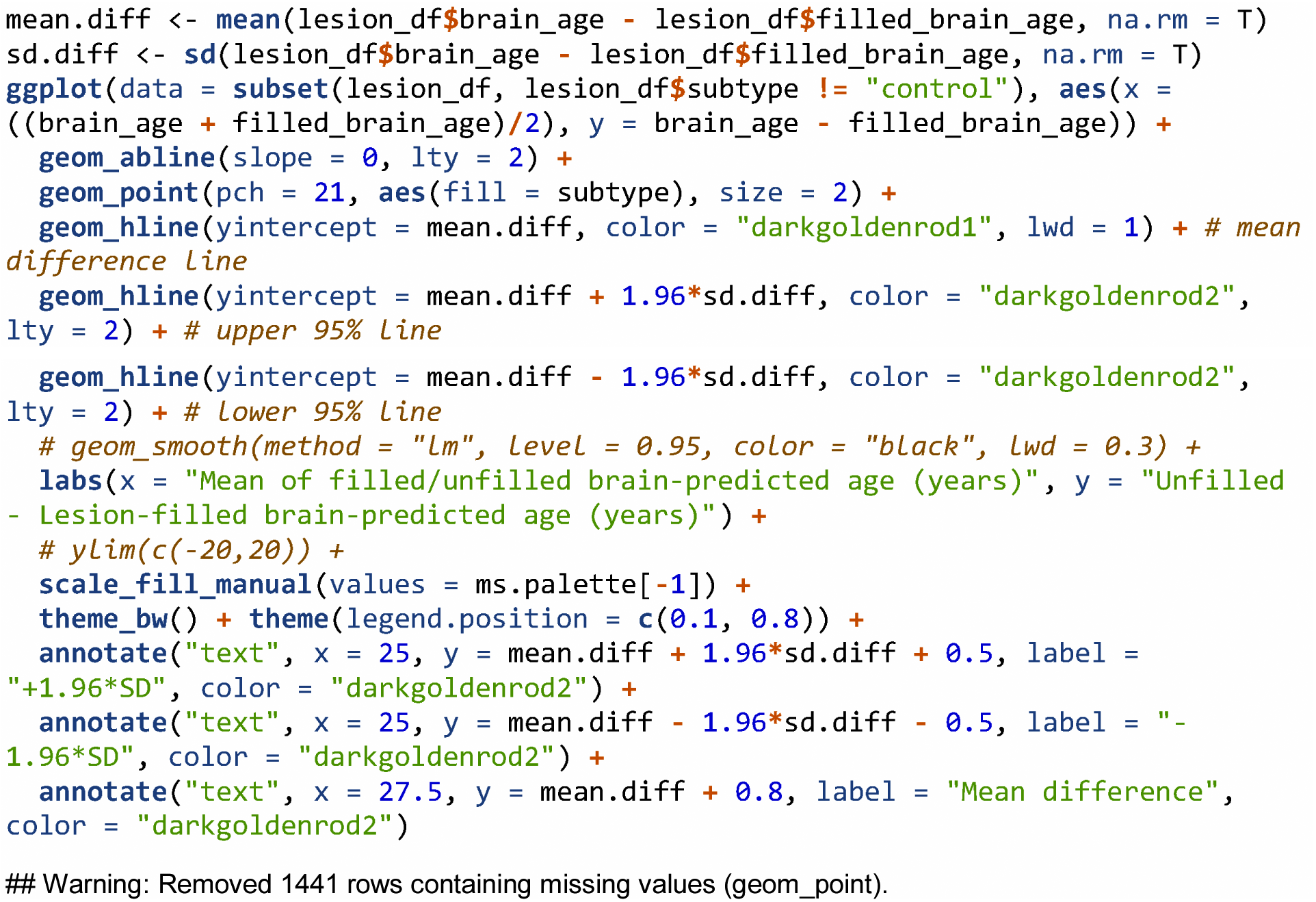

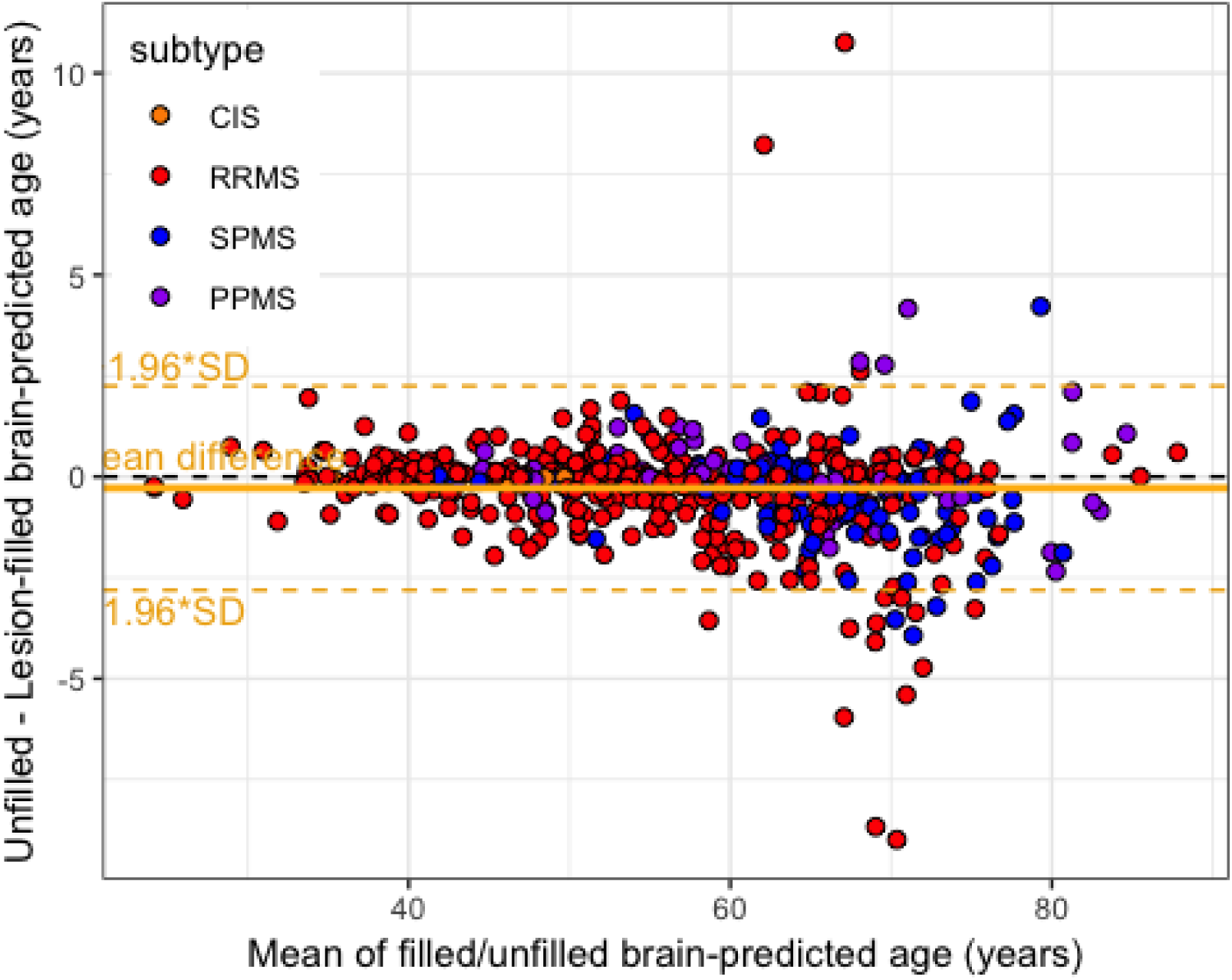

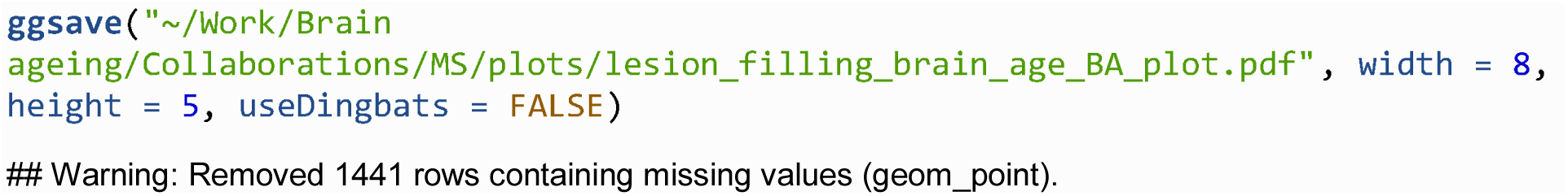

**LME model predicted brain-PAD based on subtype**

This analysis excluded controls.

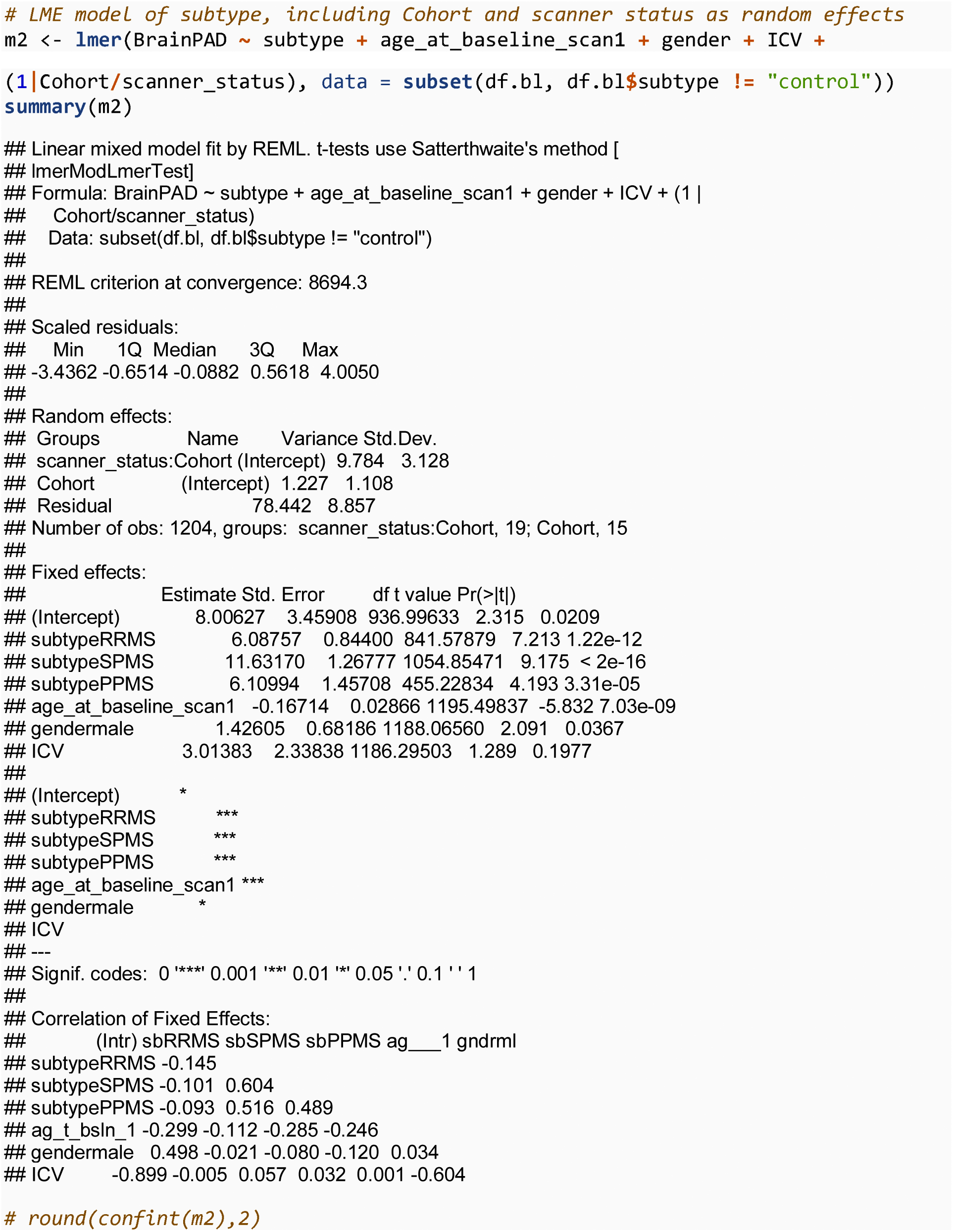

**Brain-PAD estimated marginal means for subtypes**

Generate EMMs for all MS subtypes. LME adjusting for age, gender, ICV, cohort and scanner status.

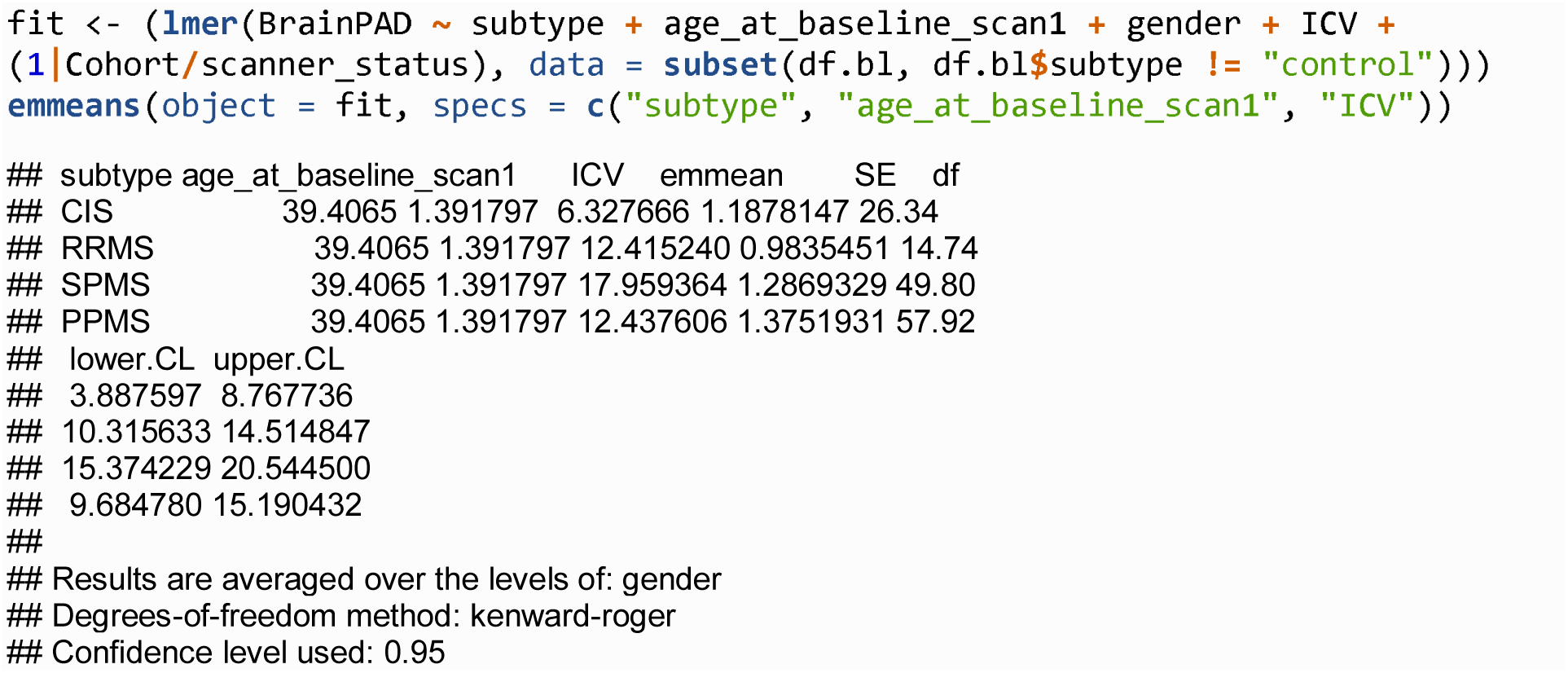

**Brain-PAD boxplot by MS subtype**

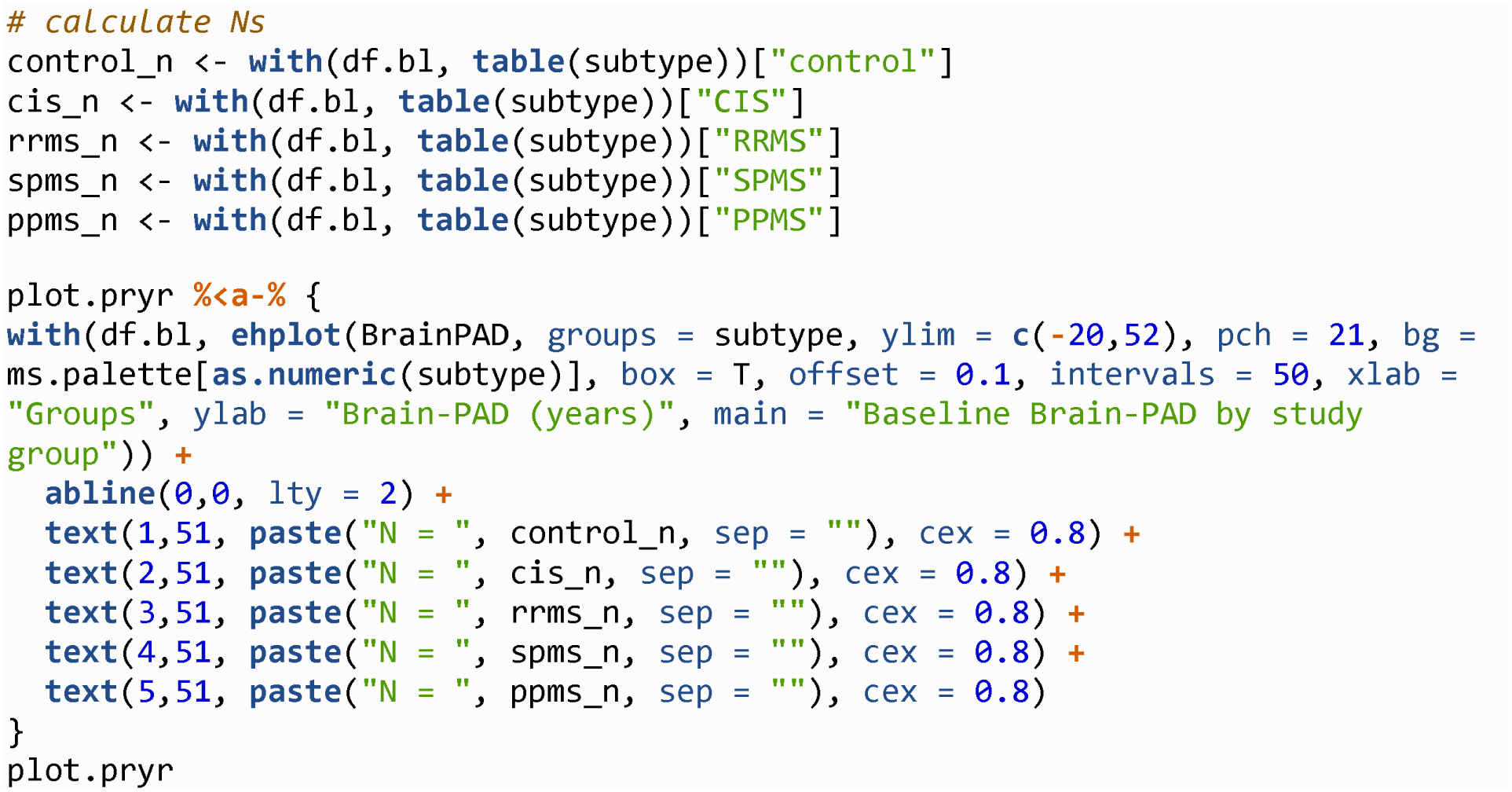

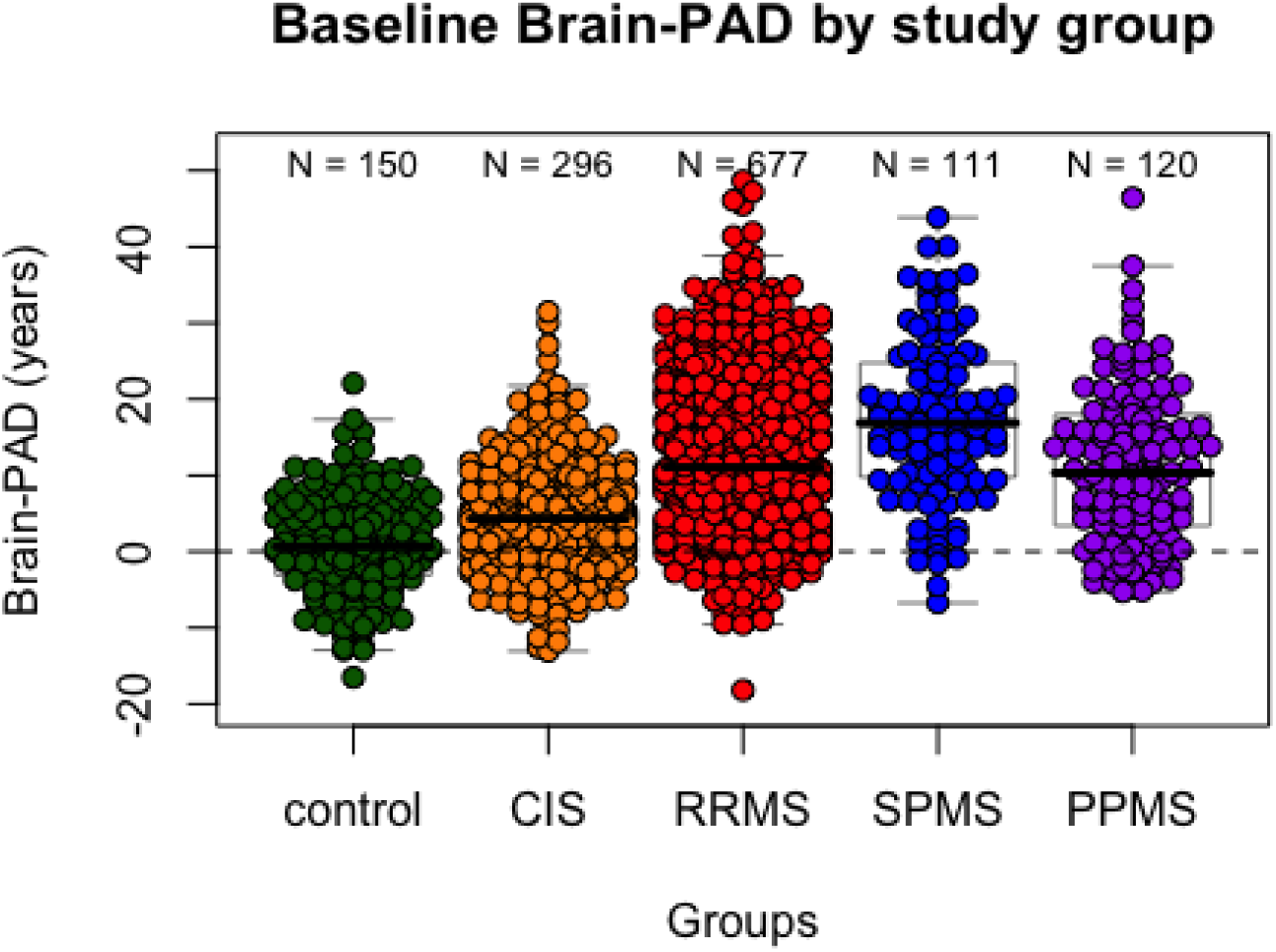

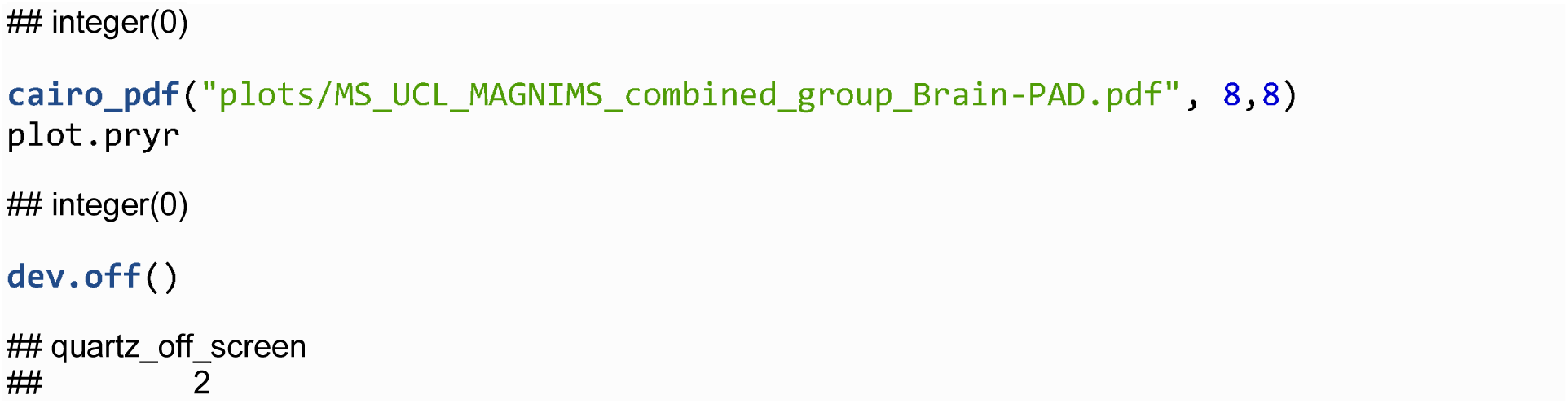

#### Brain-PAD boxplot by MS subtype in cohort UCL3 only

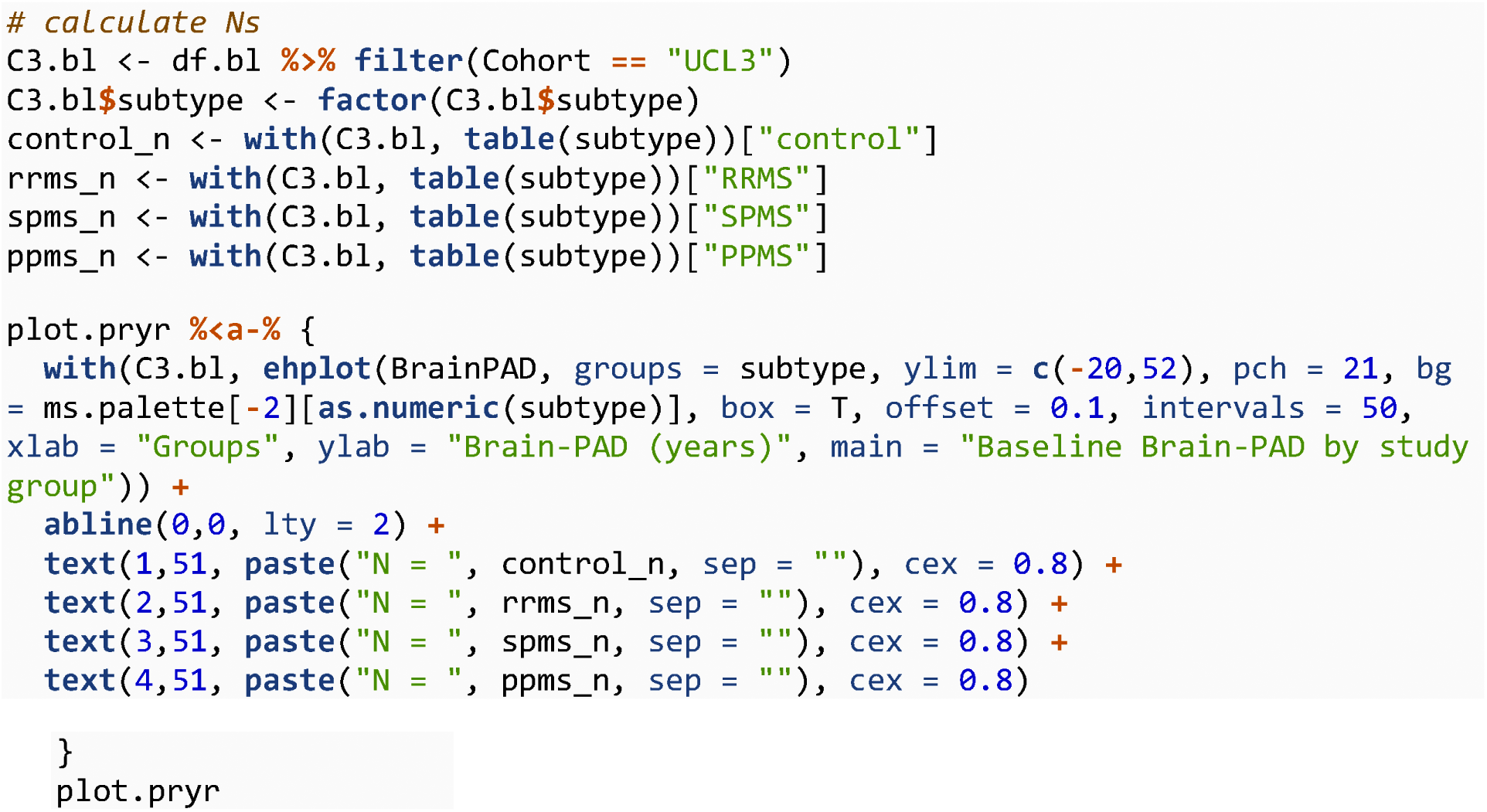

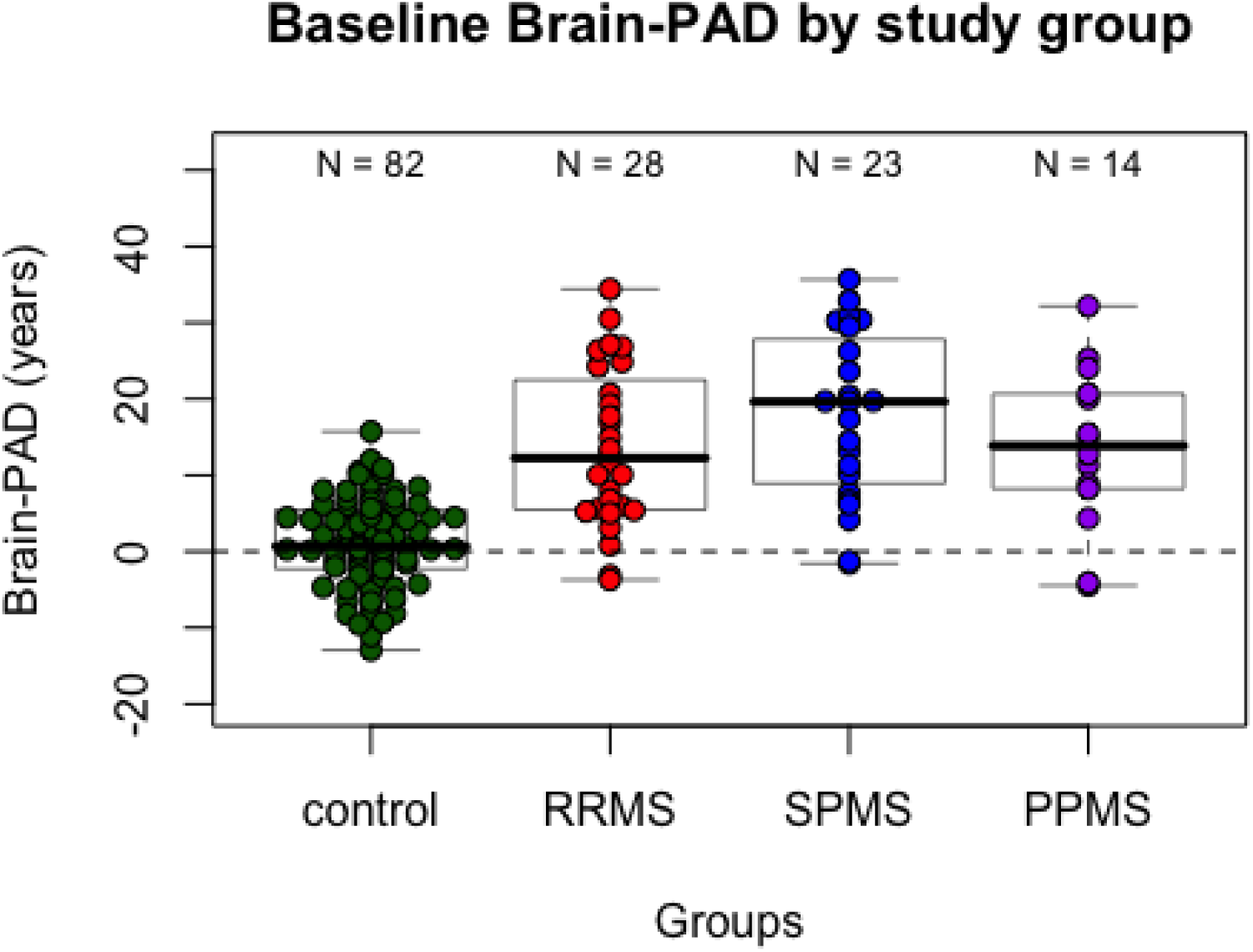

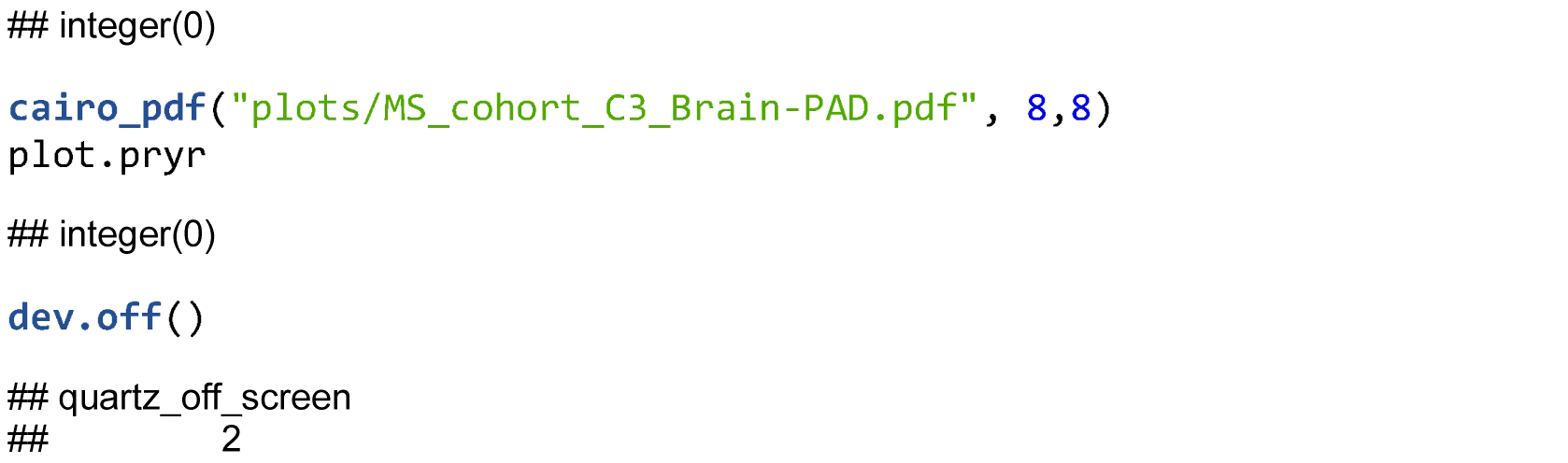

**Post-hoc pairwise brain-PAD comparison of subtypes**

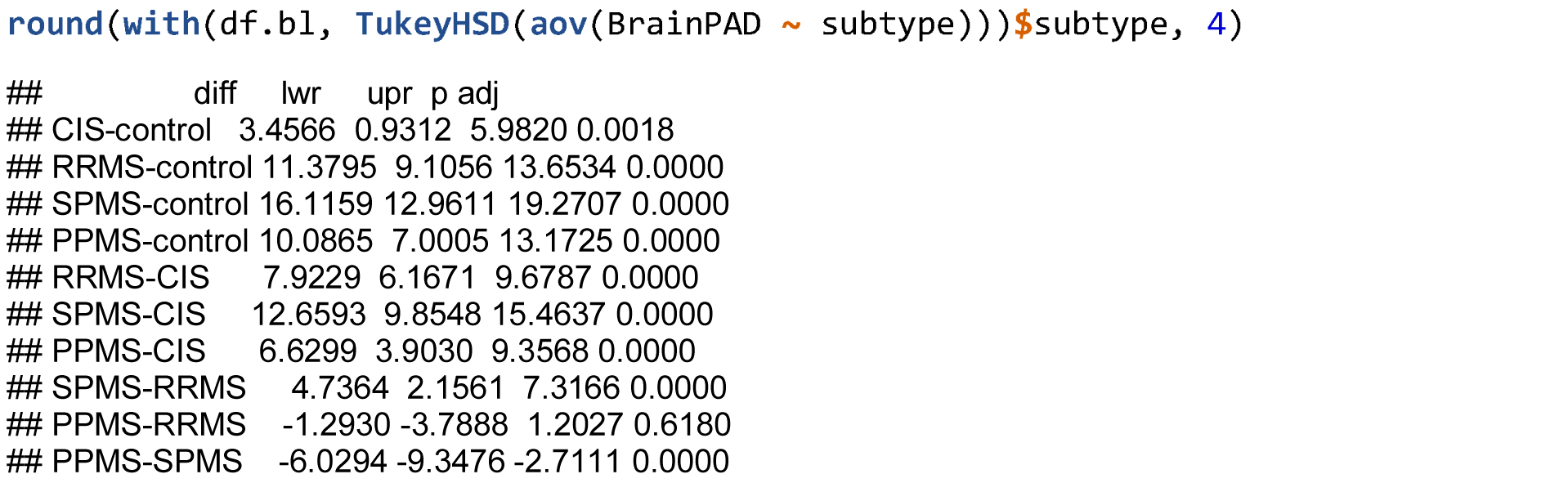

Brain-PAD by subtype descriptive statistics

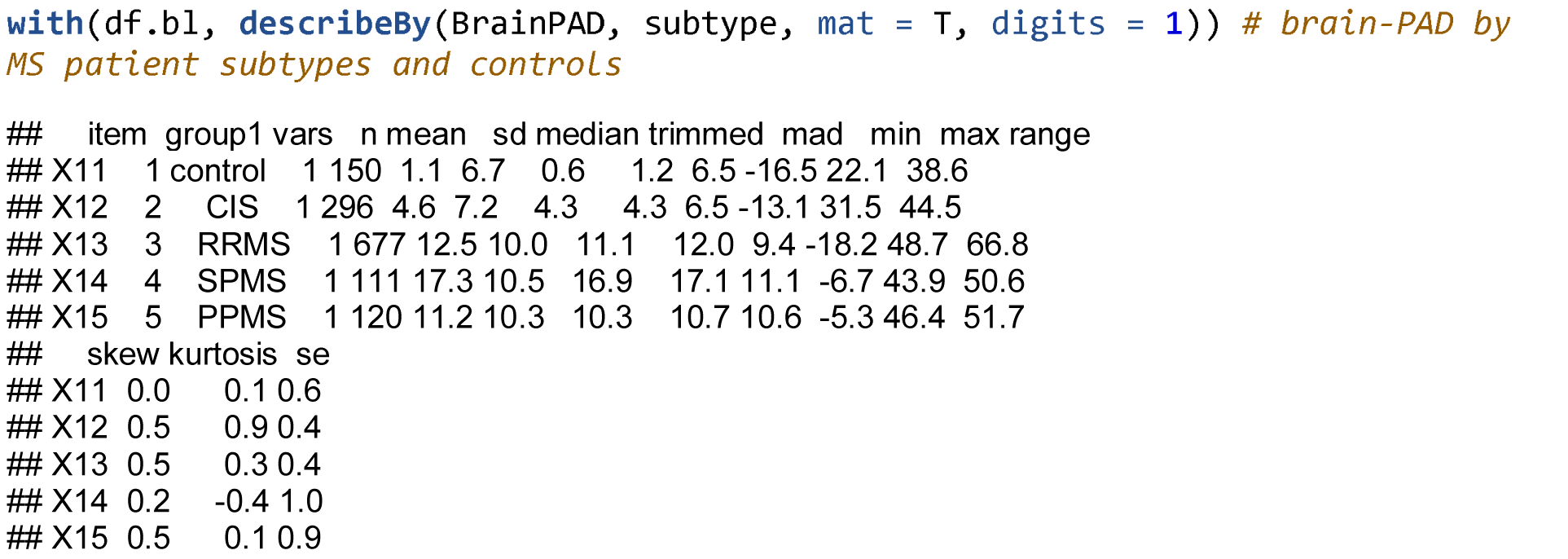

### Correlates of brain-PAD at baseline

EDSS score, an index of disability

LME accounting for fixed effects of age at baseline, gender, ICV and random effects of Cohort and scanner status.

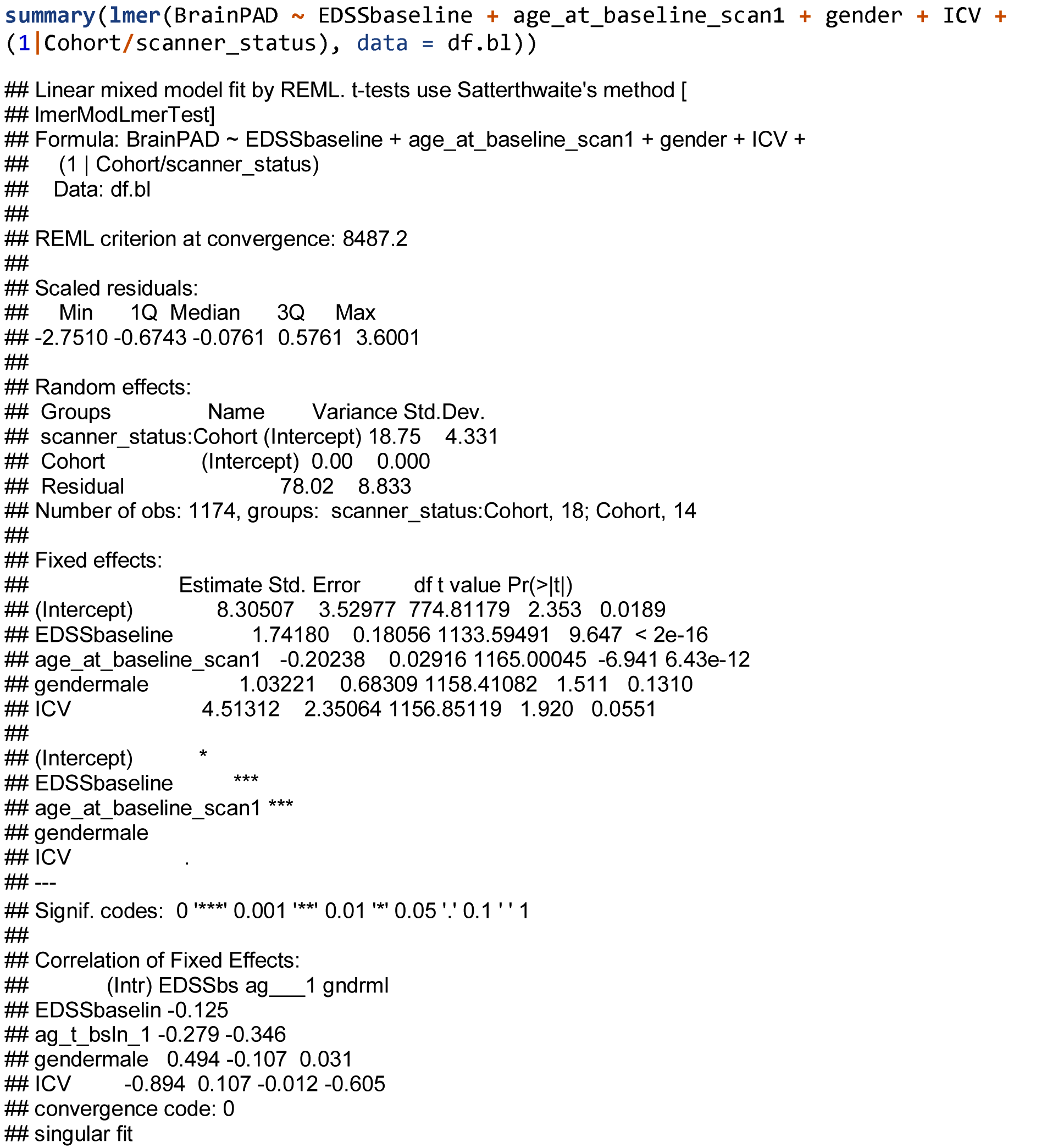

When predicting brain-PAD in a LME model, the effect of EDSS at baseline beta = 1.74, 95% CI = 1.39, 2.09, p = < 2.22e-16.

**Test for interaction between subtype and EDSS on brain-PAD:**

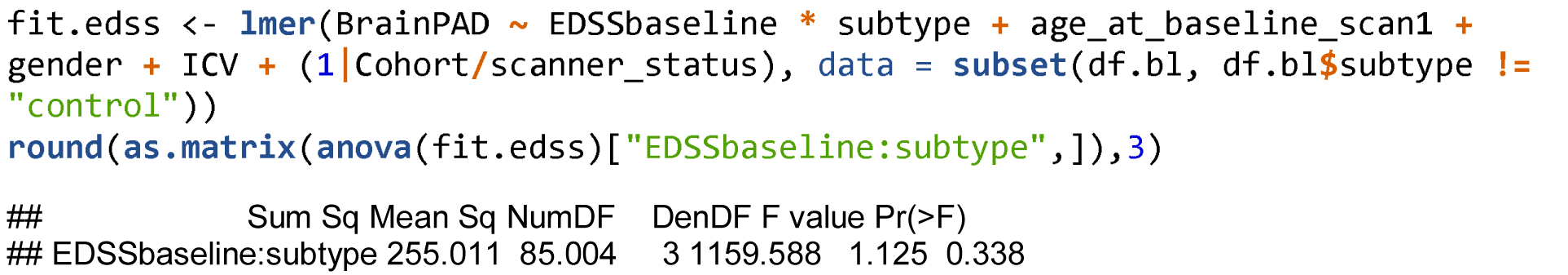

Use simple slopes from jtools to extract adjusted slopes for each subtype.

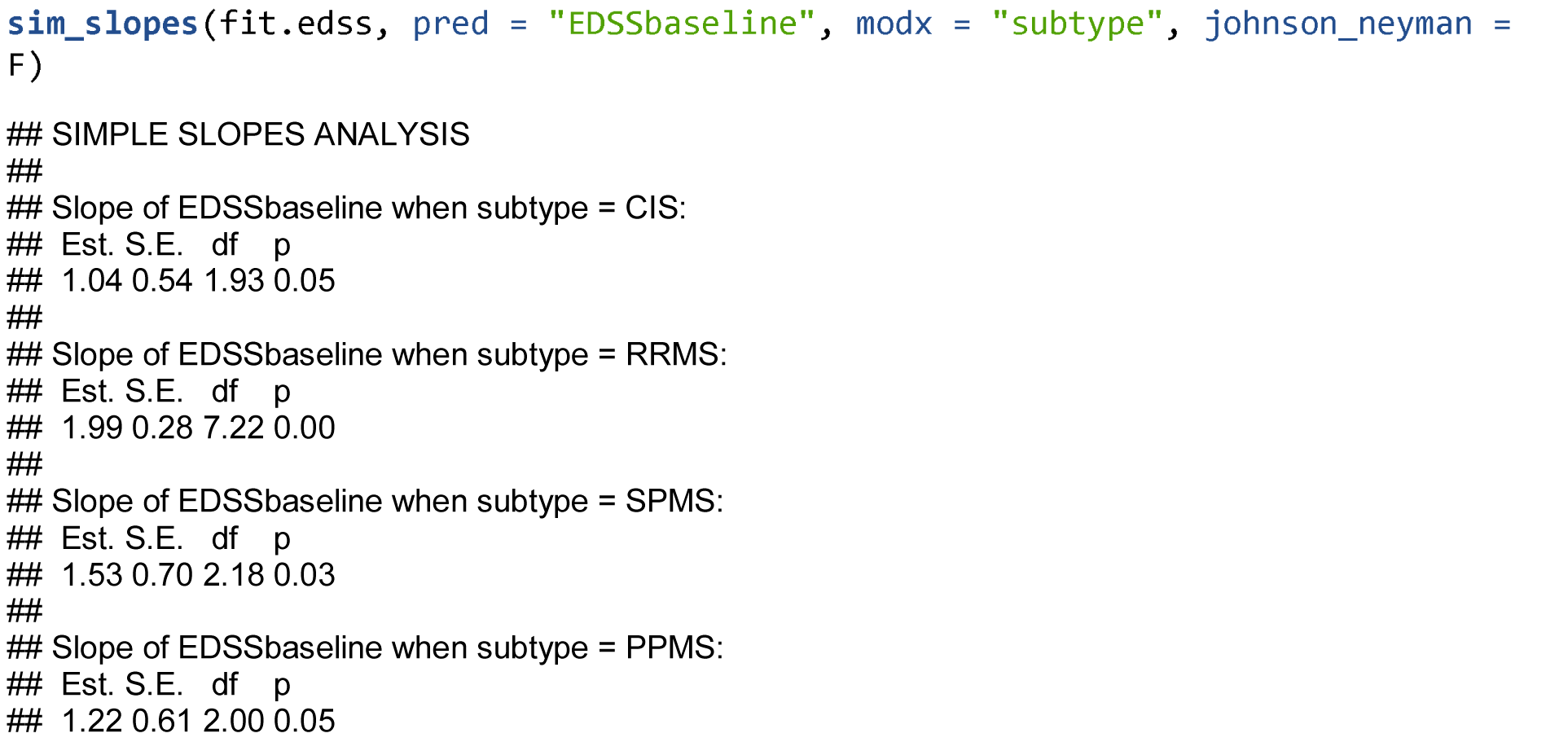

Use interact_plot() from jtools to plot the adjusted slopes per group.

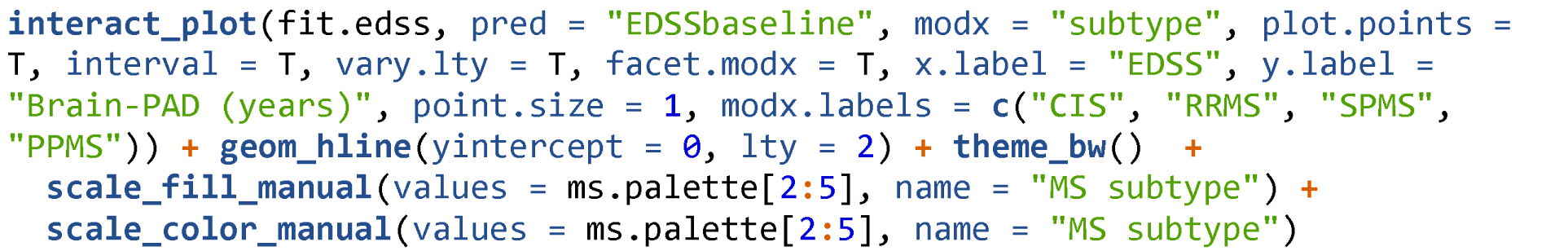

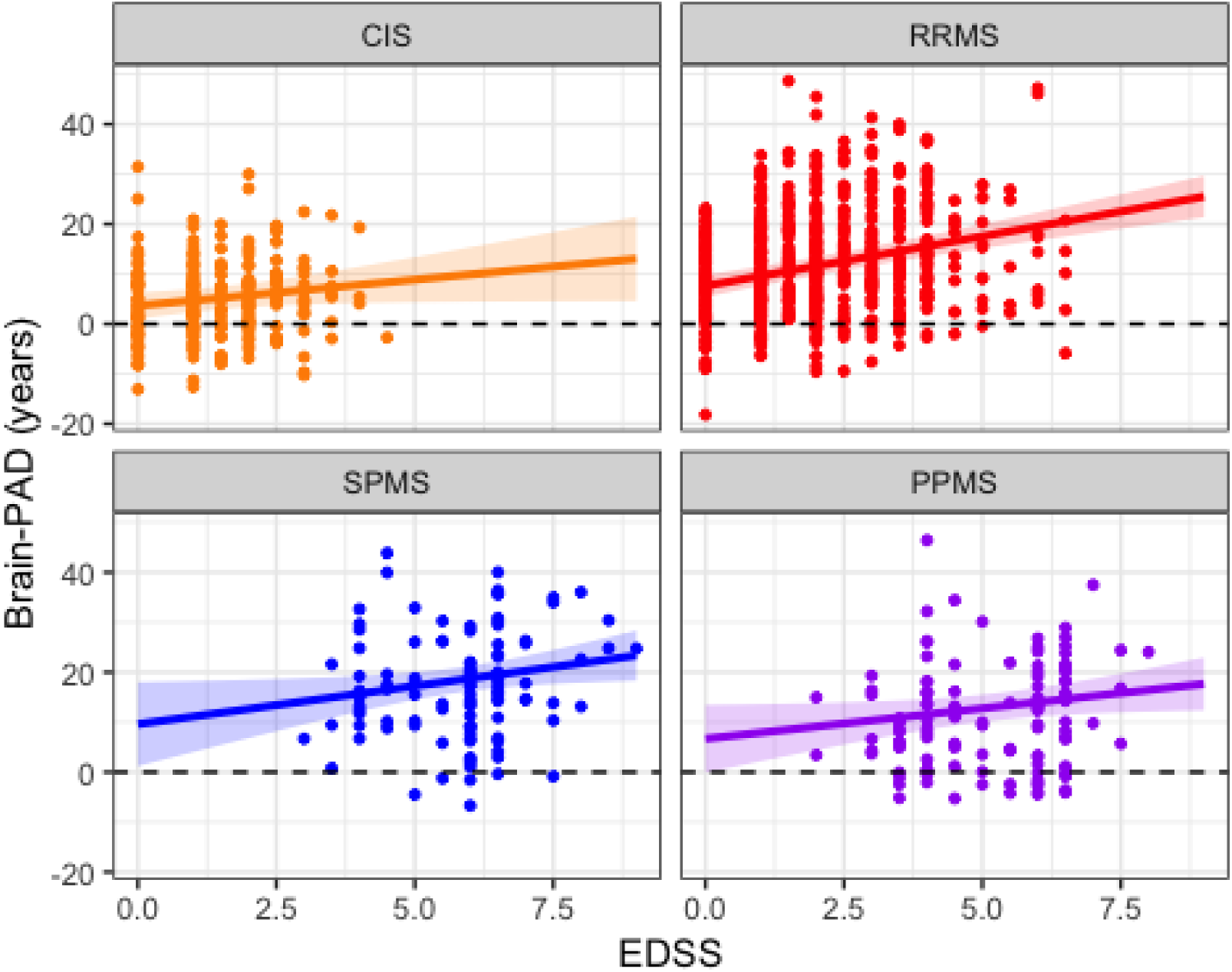

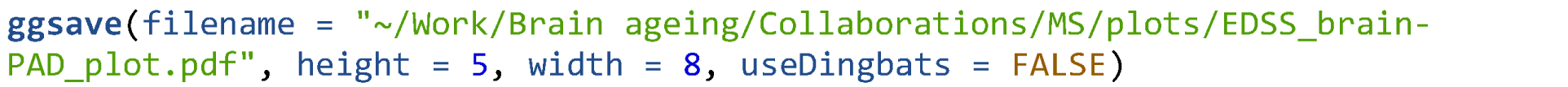

### Age at diagnosis

LME accounting for fixed effects of age at baseline, gender, ICV and random effects of Cohort and scanner status. Exclude CIS patients and healthy controls.

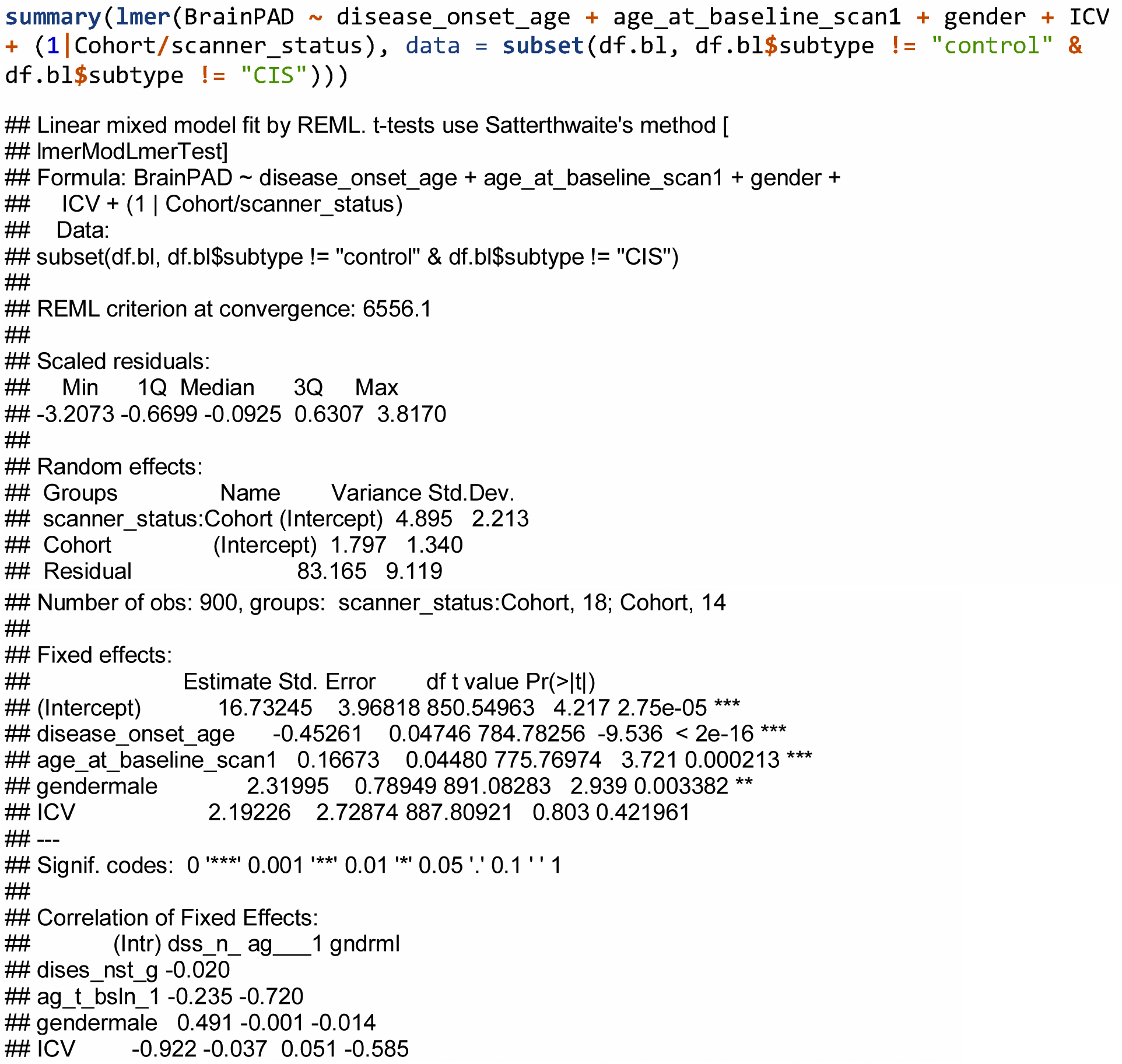

When predicting brain-PAD in a LME model, the effect of age at diagnosis at baseline beta = −0.45, 95% CI = −0.55, −0.36, p = < 2.22e-16.

### Test for interaction between subtype and age at diagnosis on brain-PAD

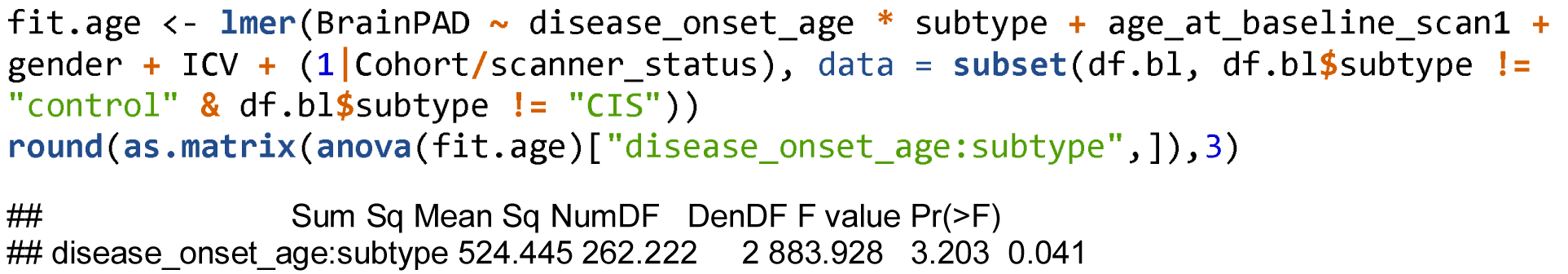

Use simple slopes from jtools to extract adjusted slopes for each subtype.

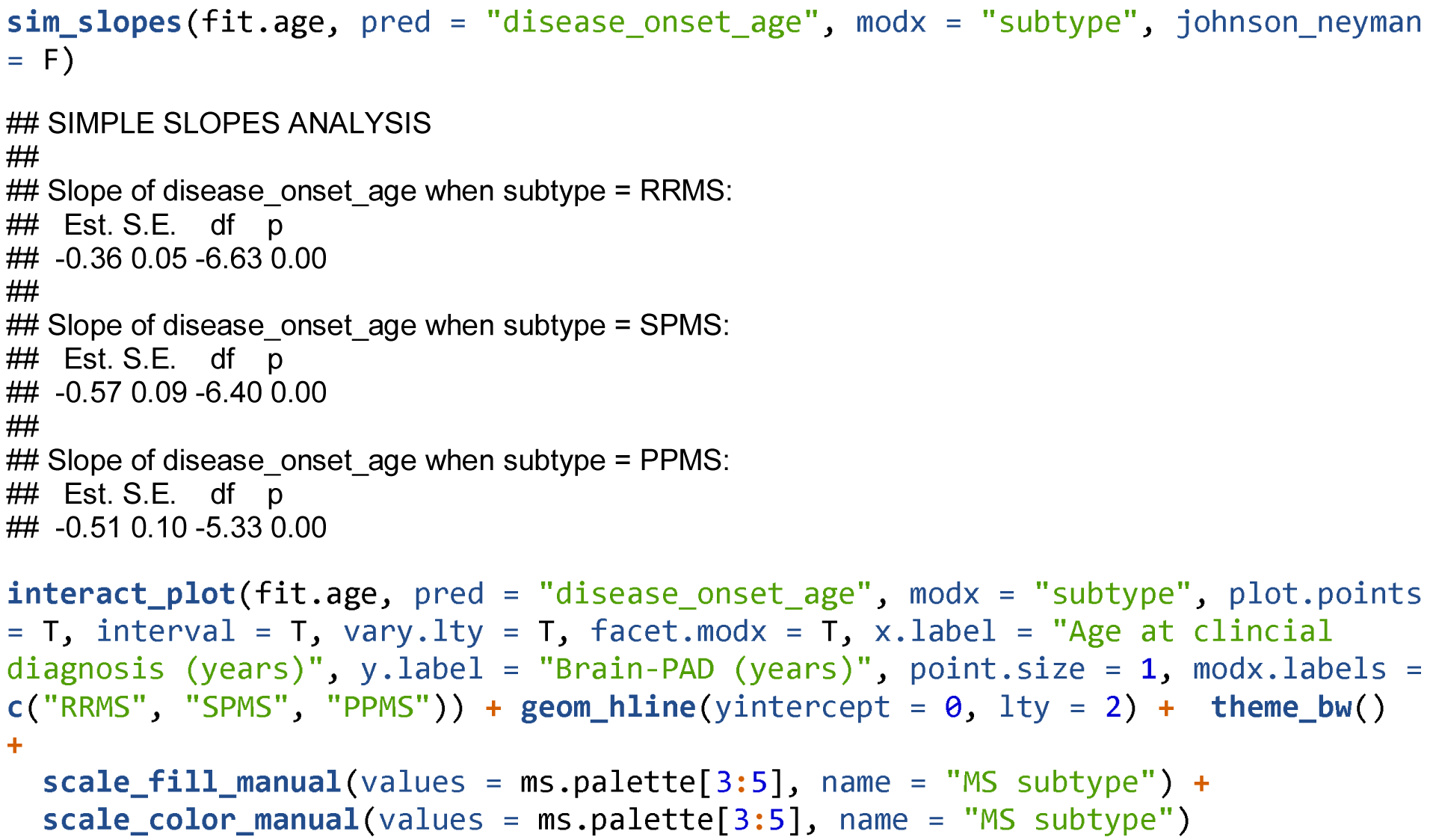

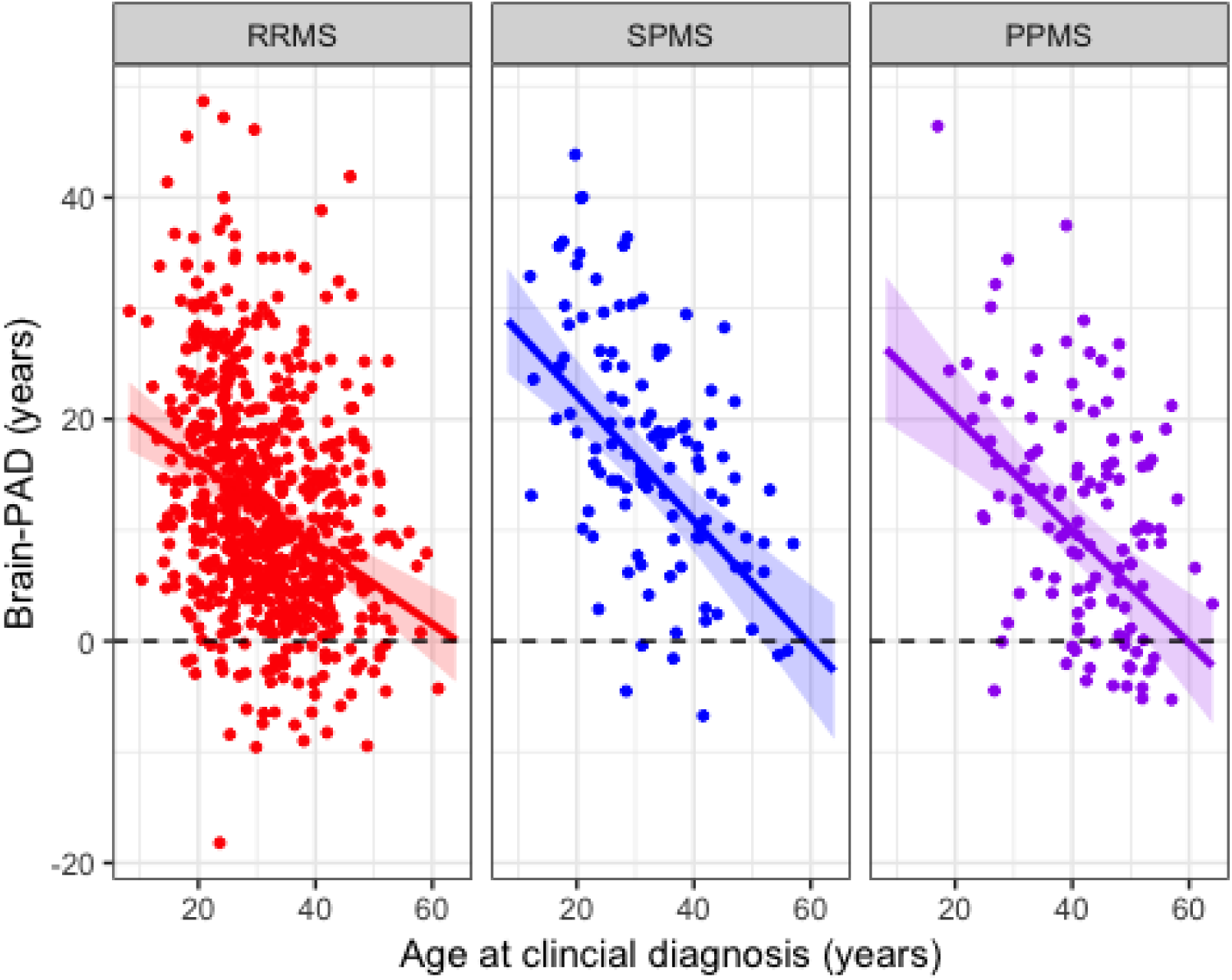

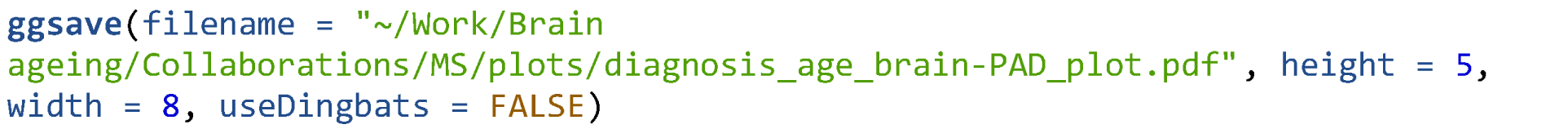

### Time since diagnosis

LME accounting for fixed effects of age at baseline, gender, ICV and random effects of Cohort and scanner status. Exclude controls, CIS patients and anyone with a time since diagnosis = 0.

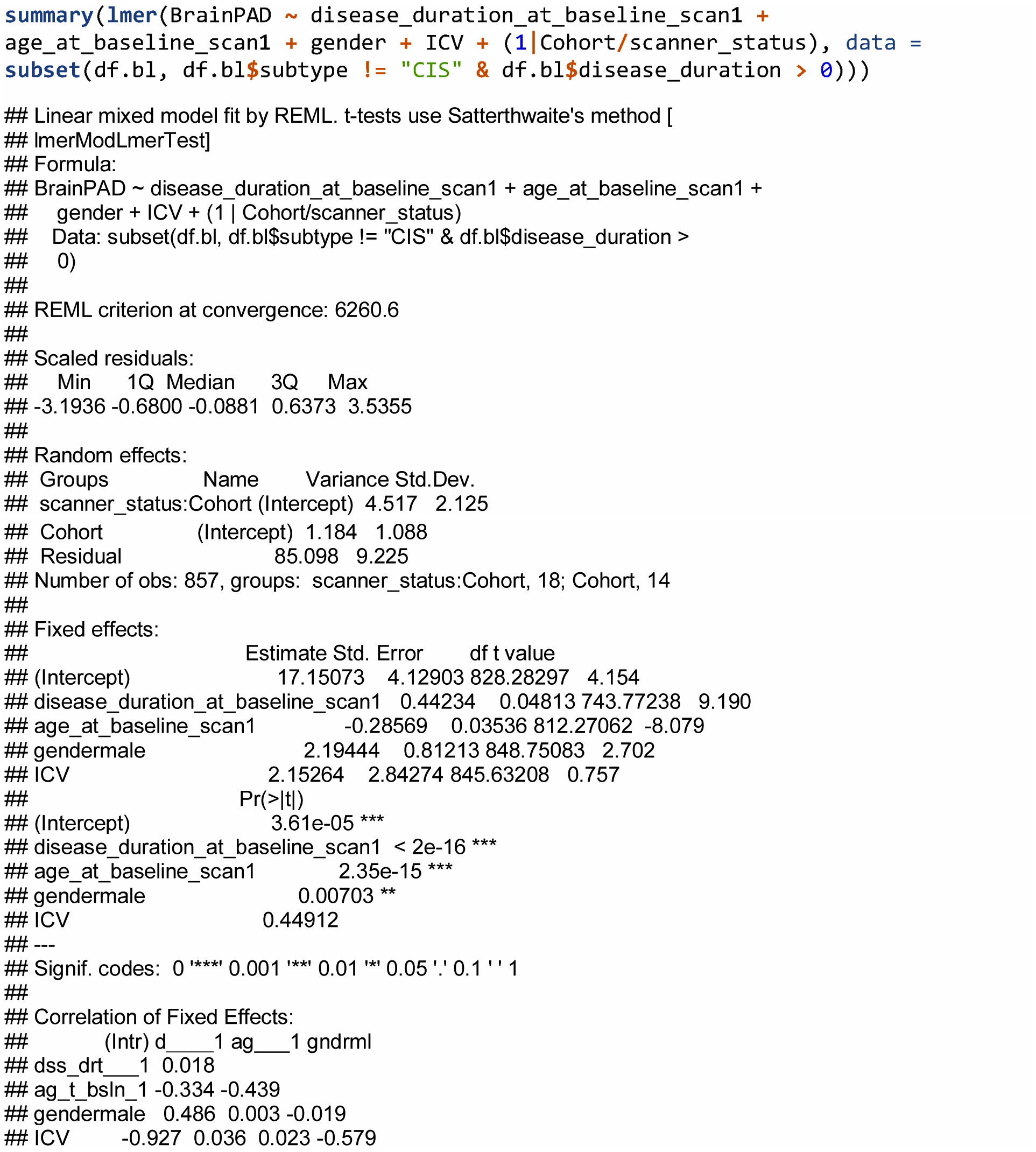

When predicting brain-PAD in a LME model, the effect of time since diagnosis at baseline beta = 0.48, 95% CI = 0.4, 0.57, p = < 2.22e-16.

### Test for interaction between subtype and time since diagnosis on brain-PAD

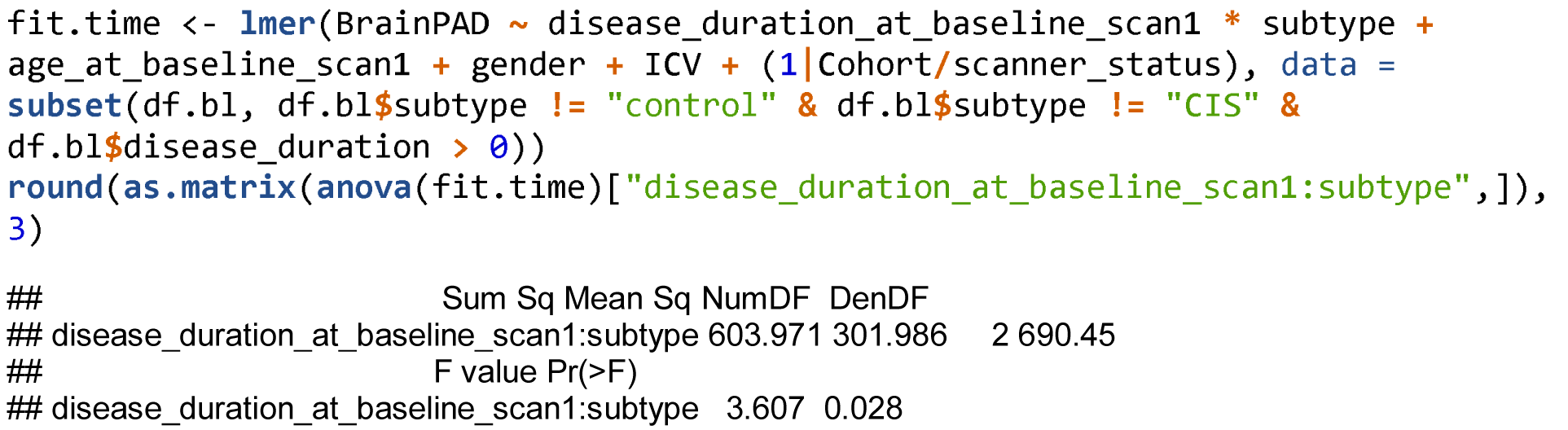

Use simple slopes from jtools to extract adjusted slopes for each subtype.

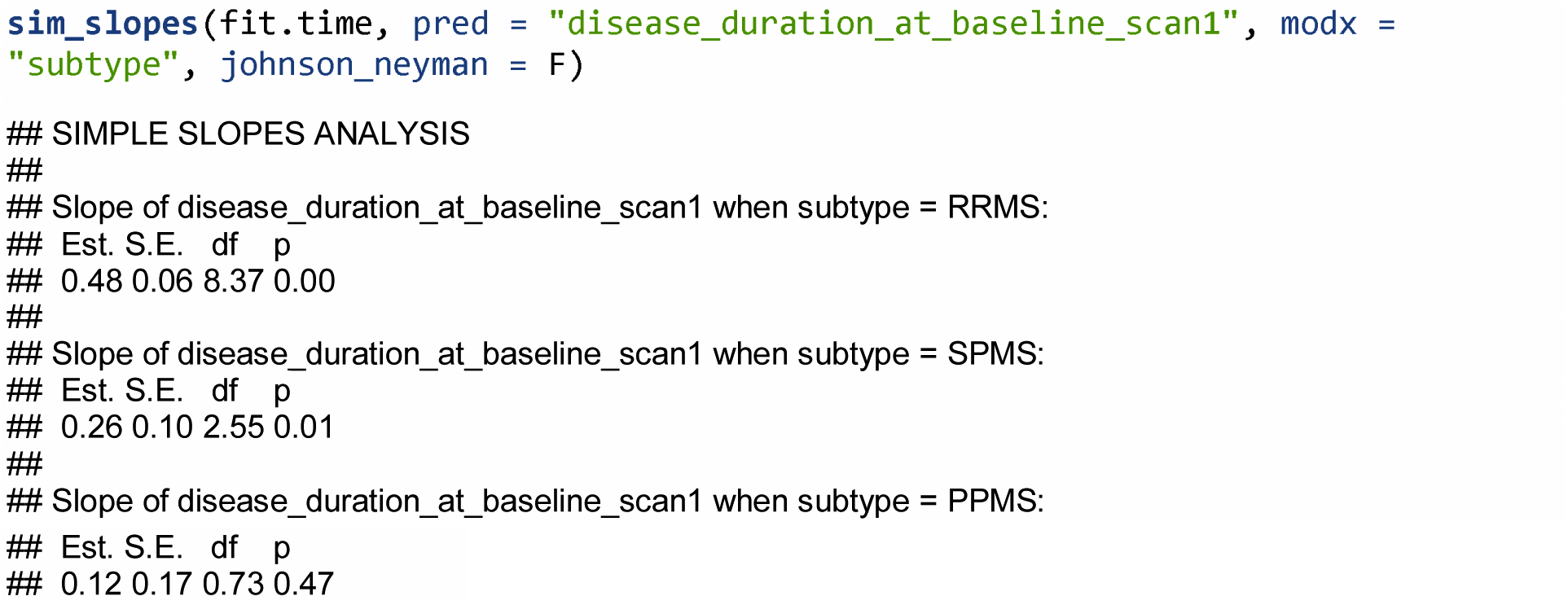

Plot

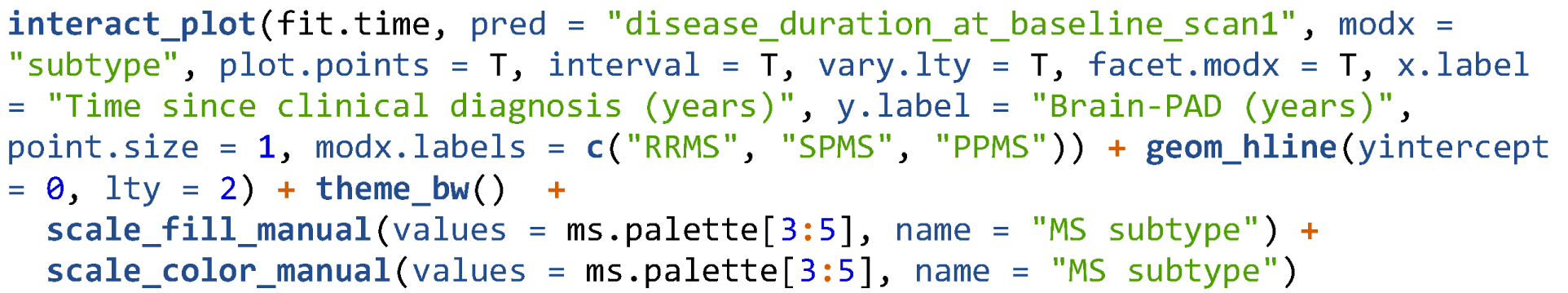

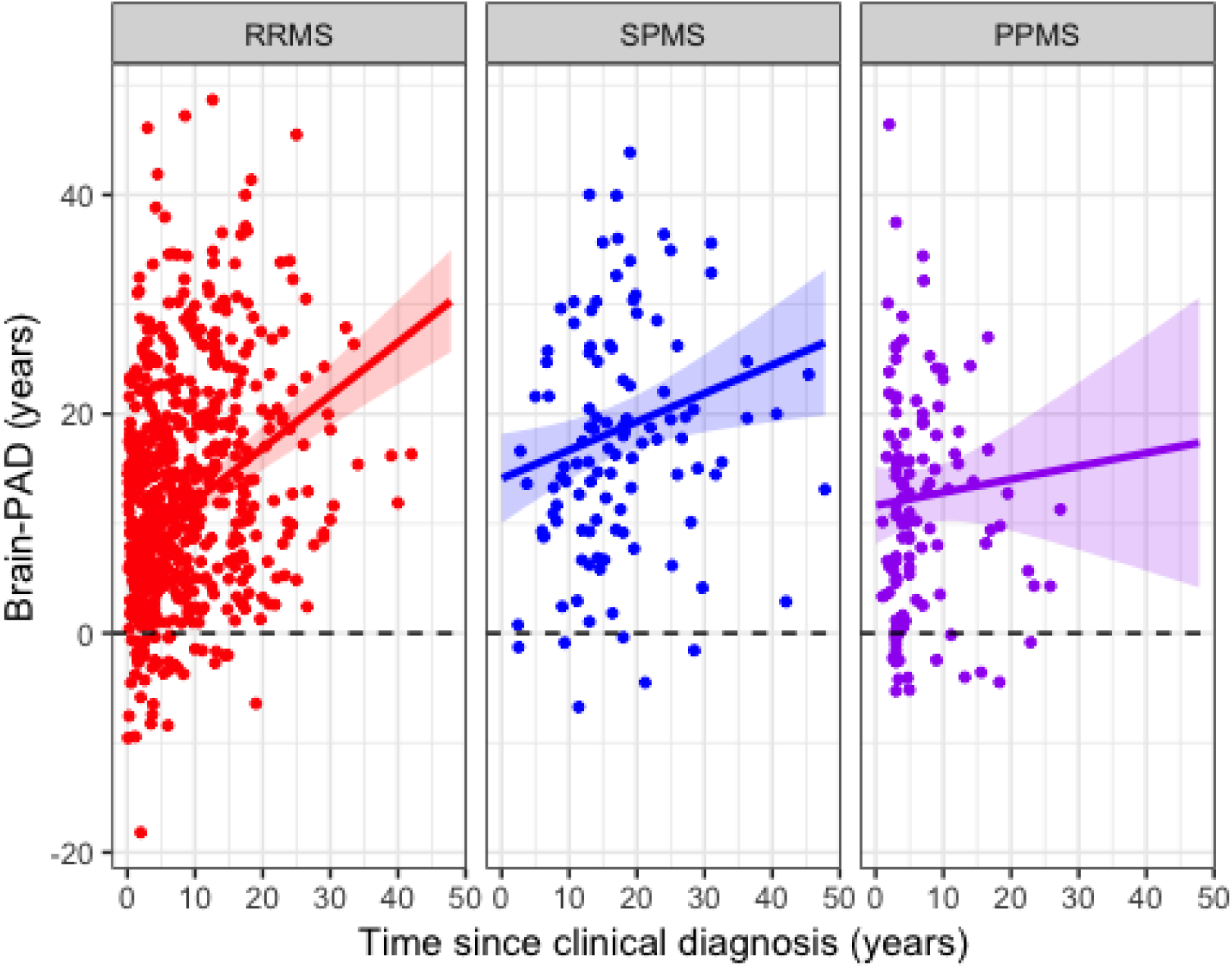

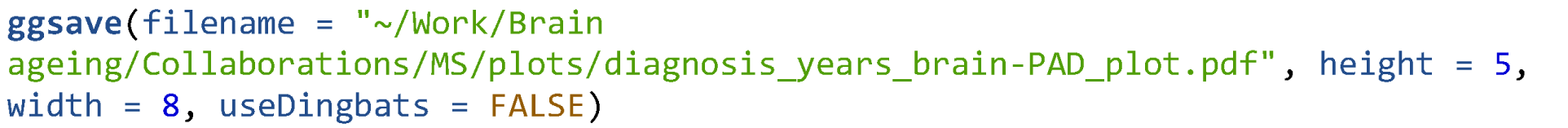

### EDSS progression survival analysis

Based on Arman Eshaghi’s code used in Eshaghi et al., 2018 Annals of Neurology.

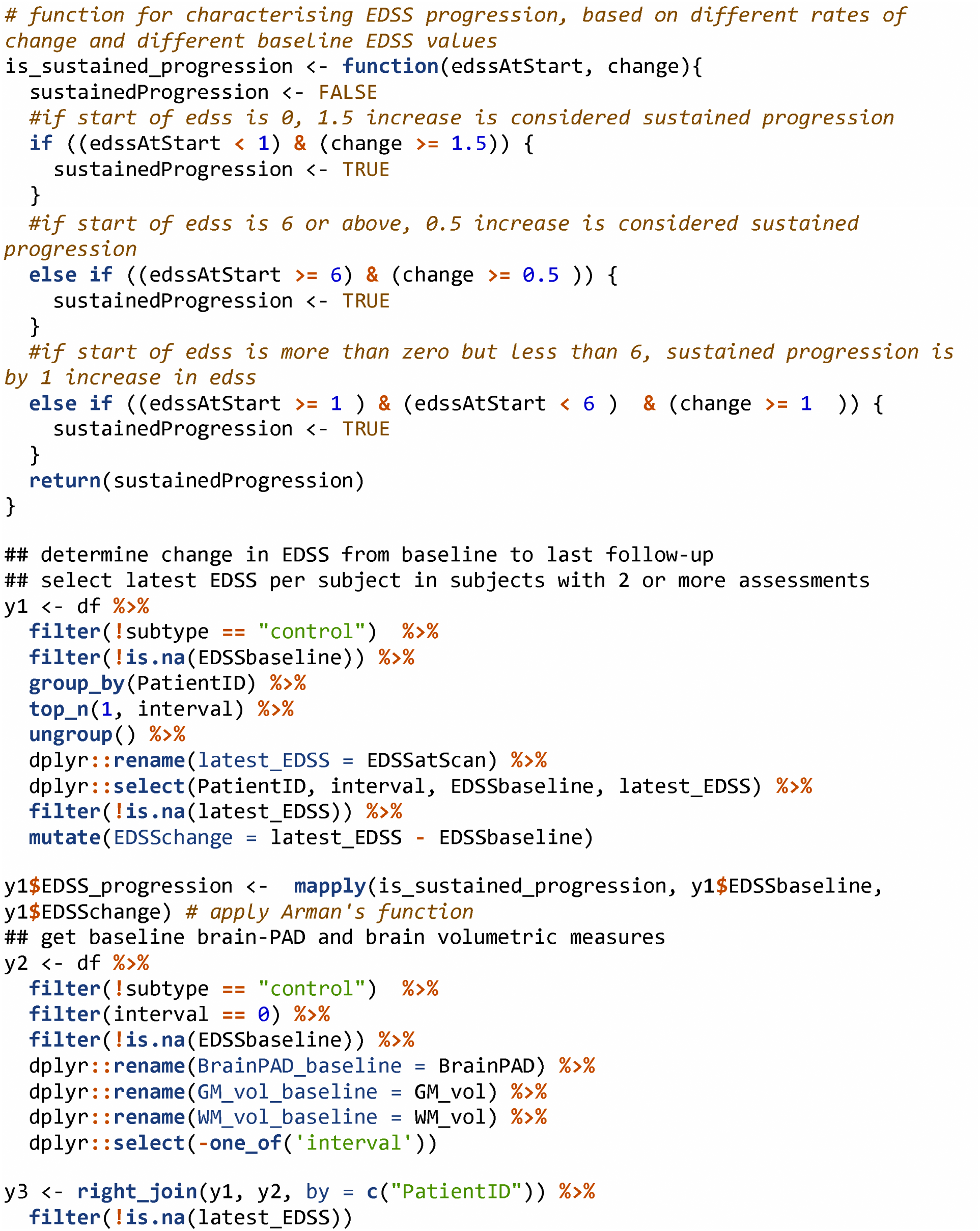

#### Numbers of EDSS progressors

The number of MS patients with >= 2 EDSS scores was 1143.

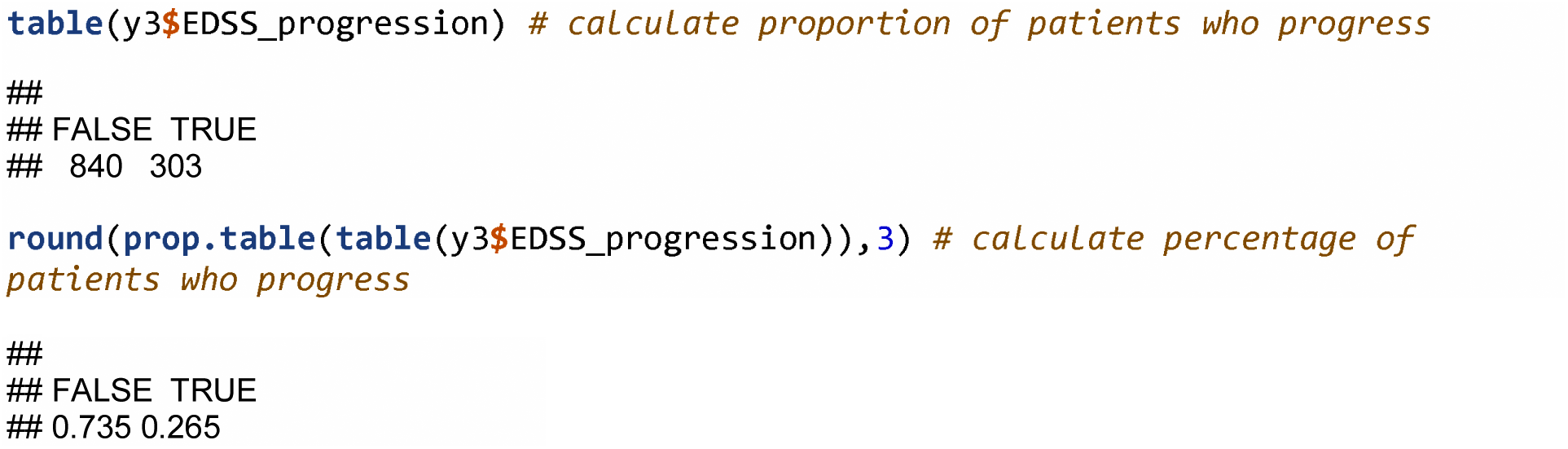

#### Run survival analysis

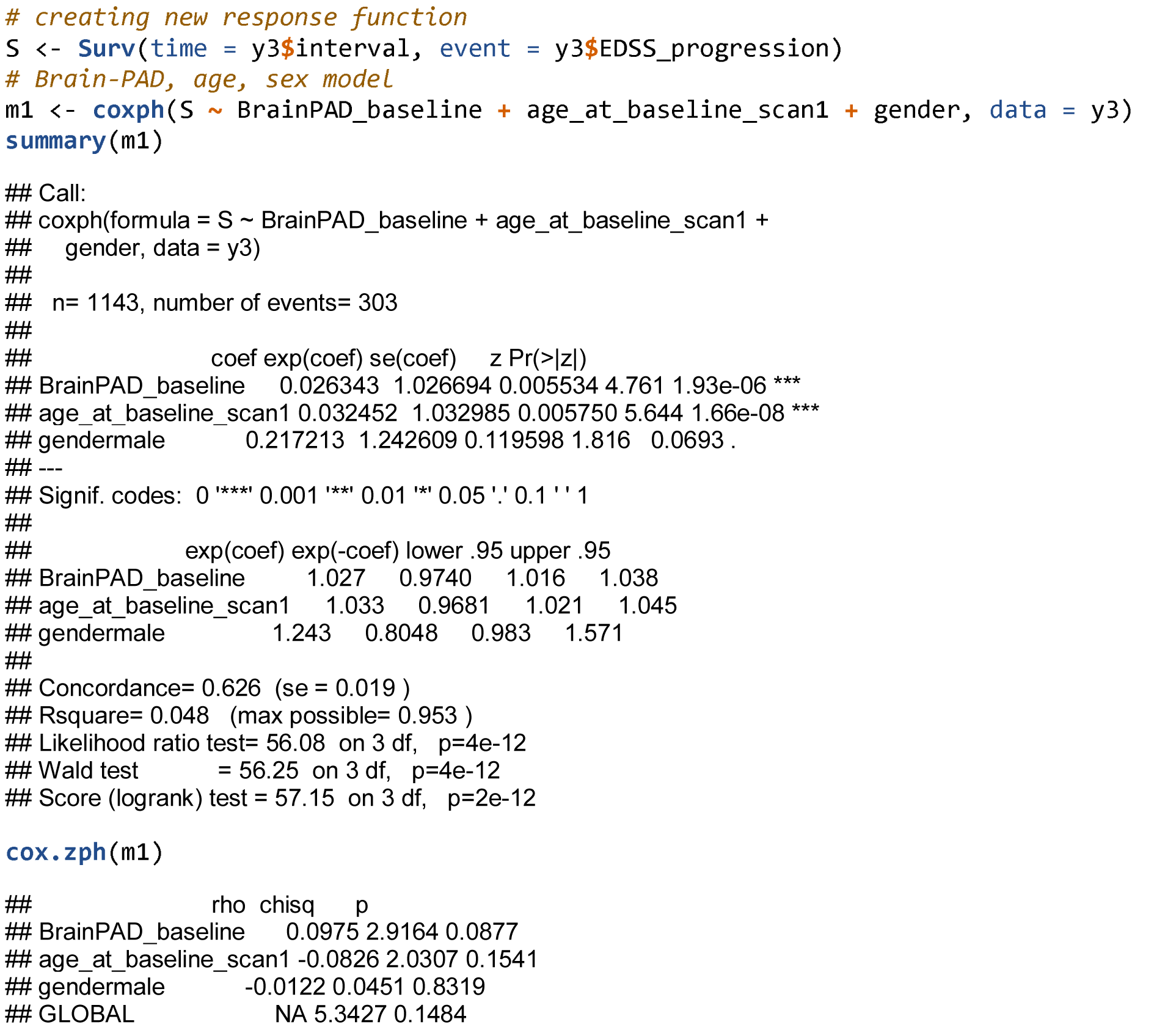

The hazard ratio for brain-PAD on time-to-disease-progression was HR (95% CI) = 1.027, 1.016, 1.038. That means for every additional +1 year of brain-PAD there is a 1.027% increase in the likelihood of EDSS progression. Extrapolated over 5 years of brain-PAD, there is a 1.141 increase in the likelihood of EDSS progression.

#### Time-to-EDSS progression Kaplan-Meier plots

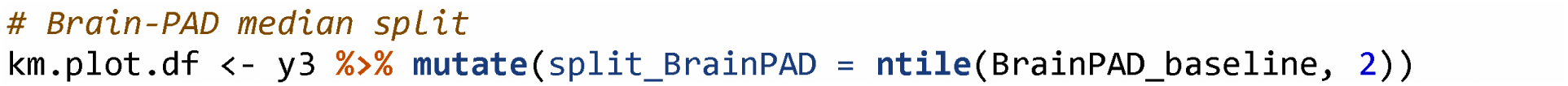

Based on a median split of brain-PAD. The median value = 9.68 years.

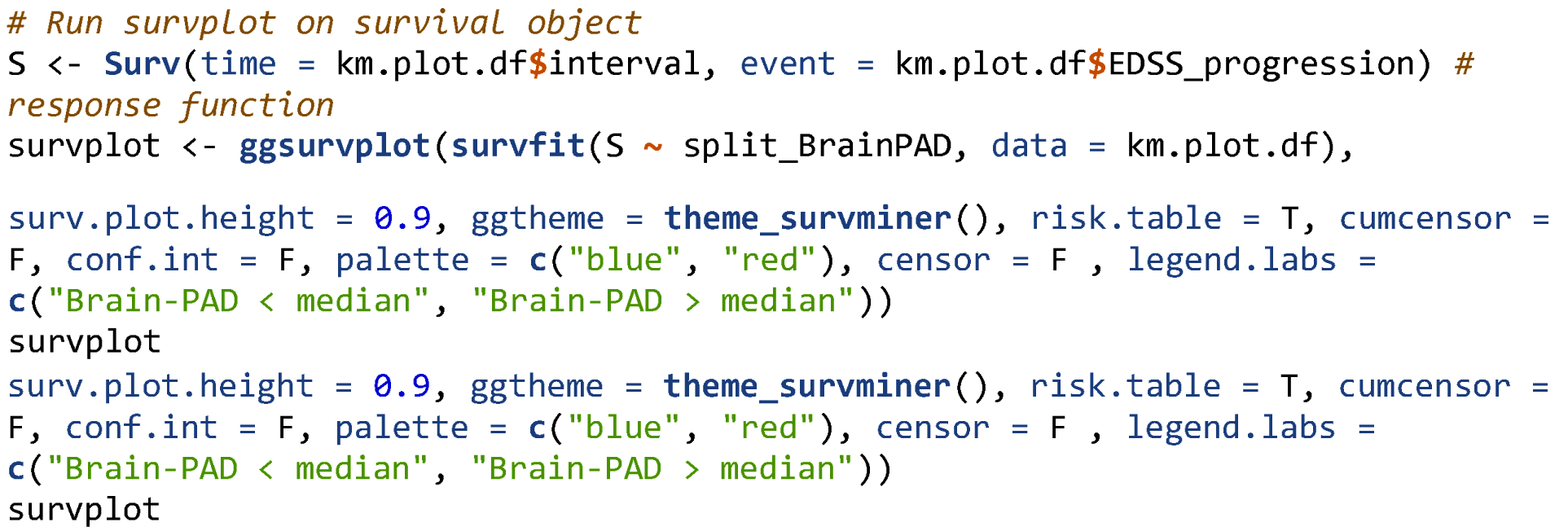

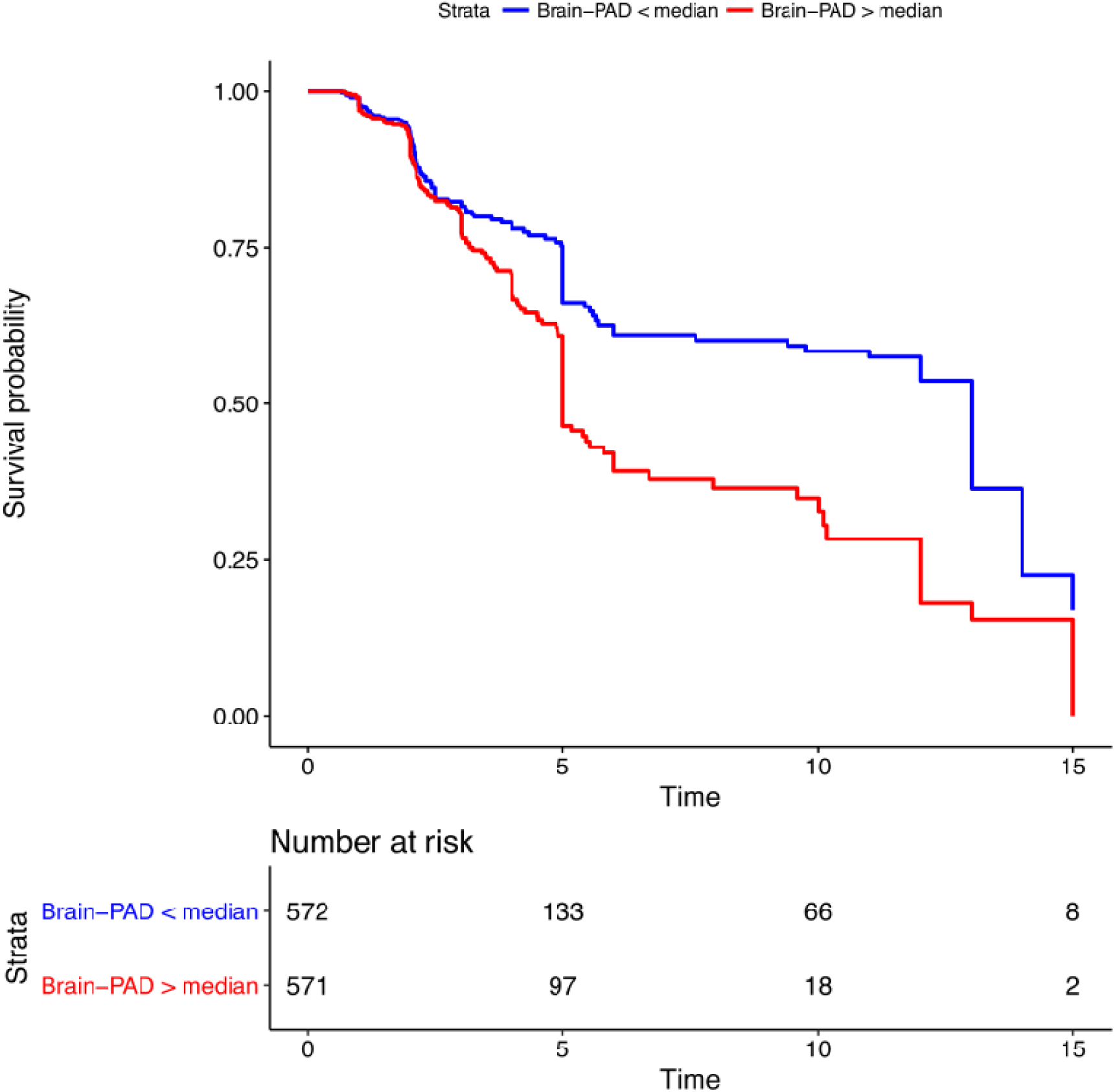

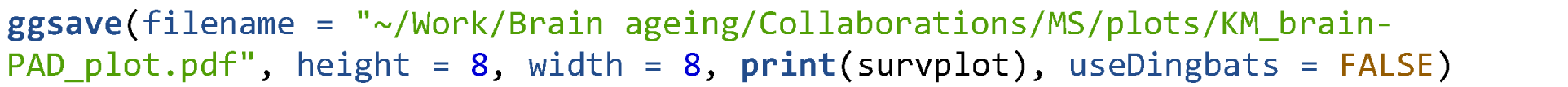

#### Longitudinal brain-age analysis

The total number of people with two or more scans was n = 1266.

**determine change in brain-PAD from baseline to last follow-up**

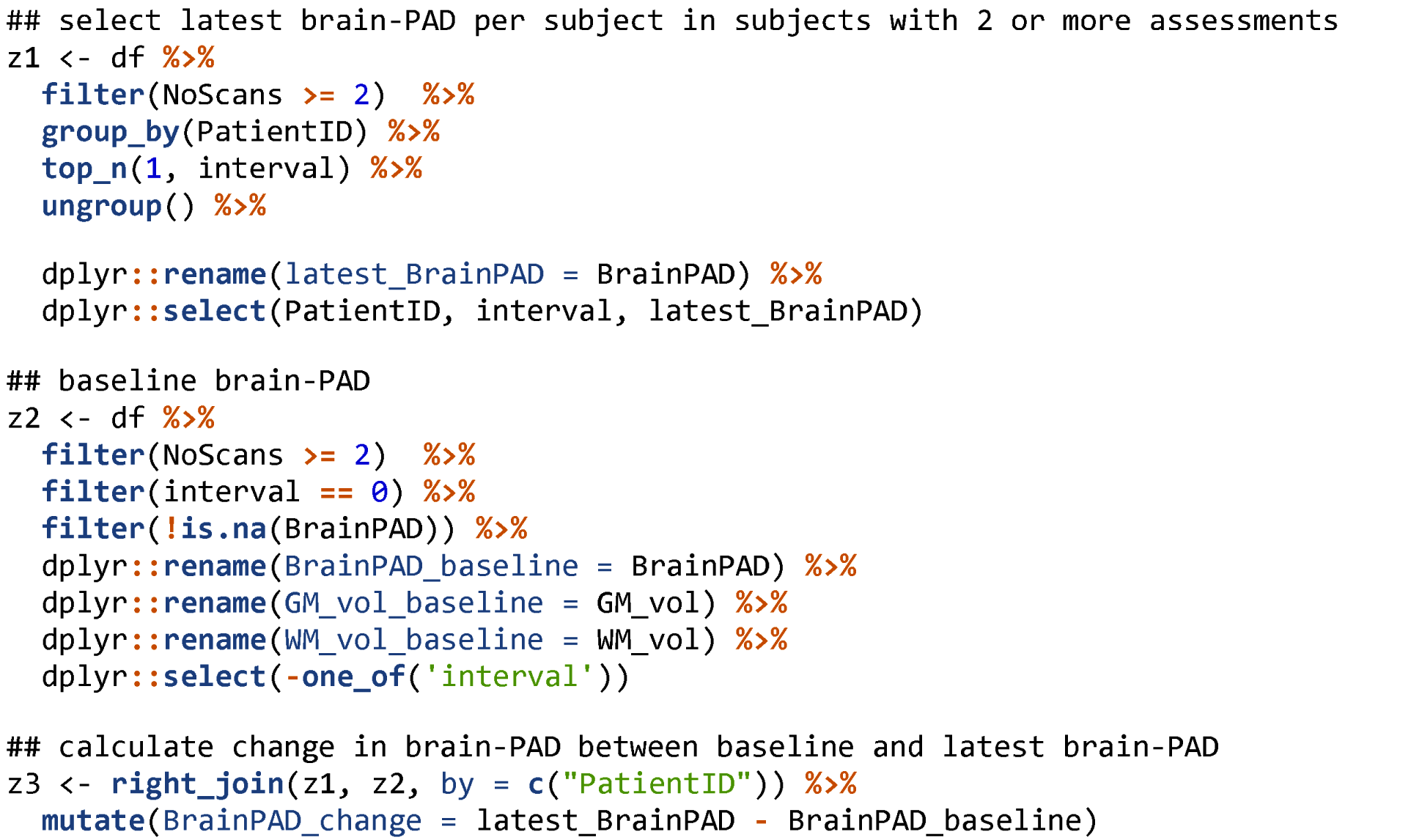

### Mean annualised rates of change in brain-PAD per group

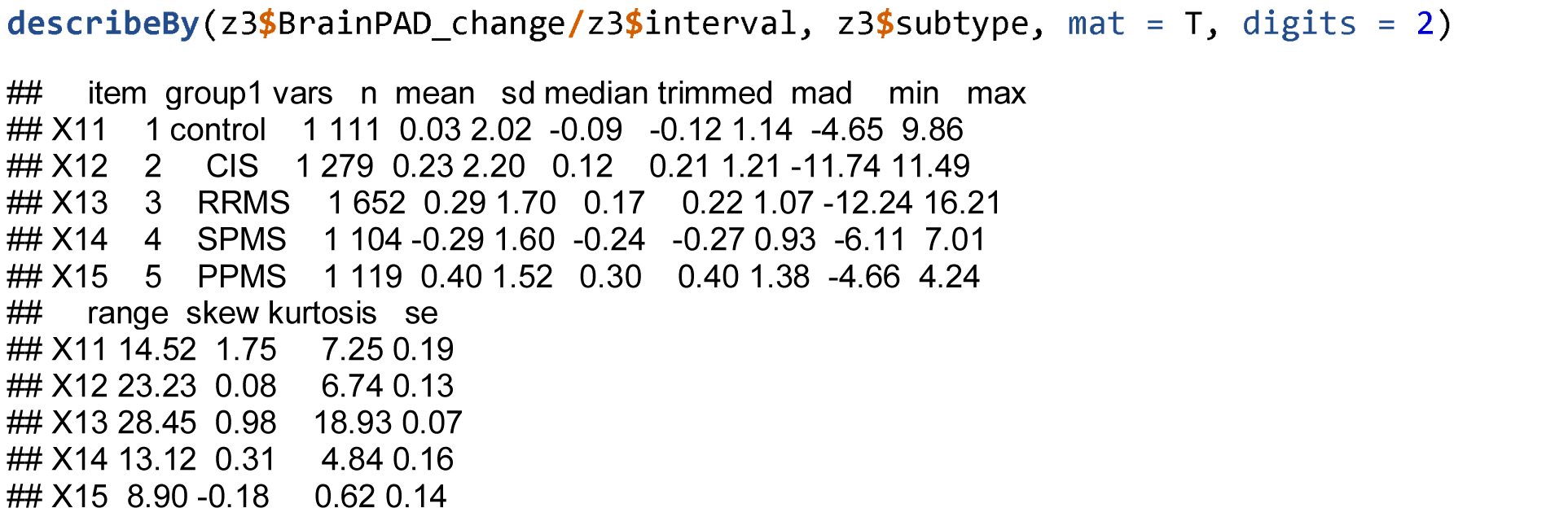

**determine change in EDSS from baseline to last follow-up**

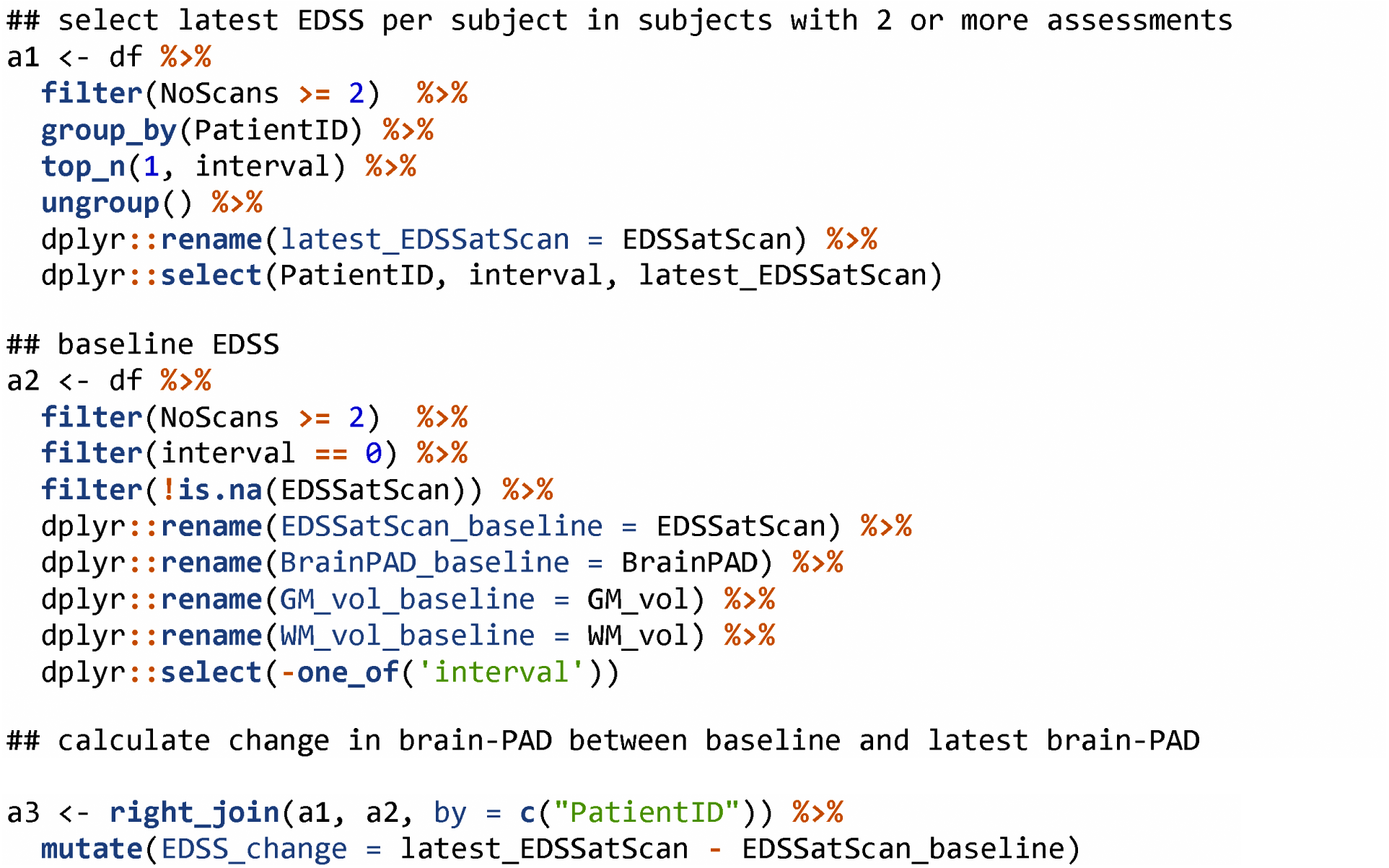

### Mean annualised rates of change in EDSS per group

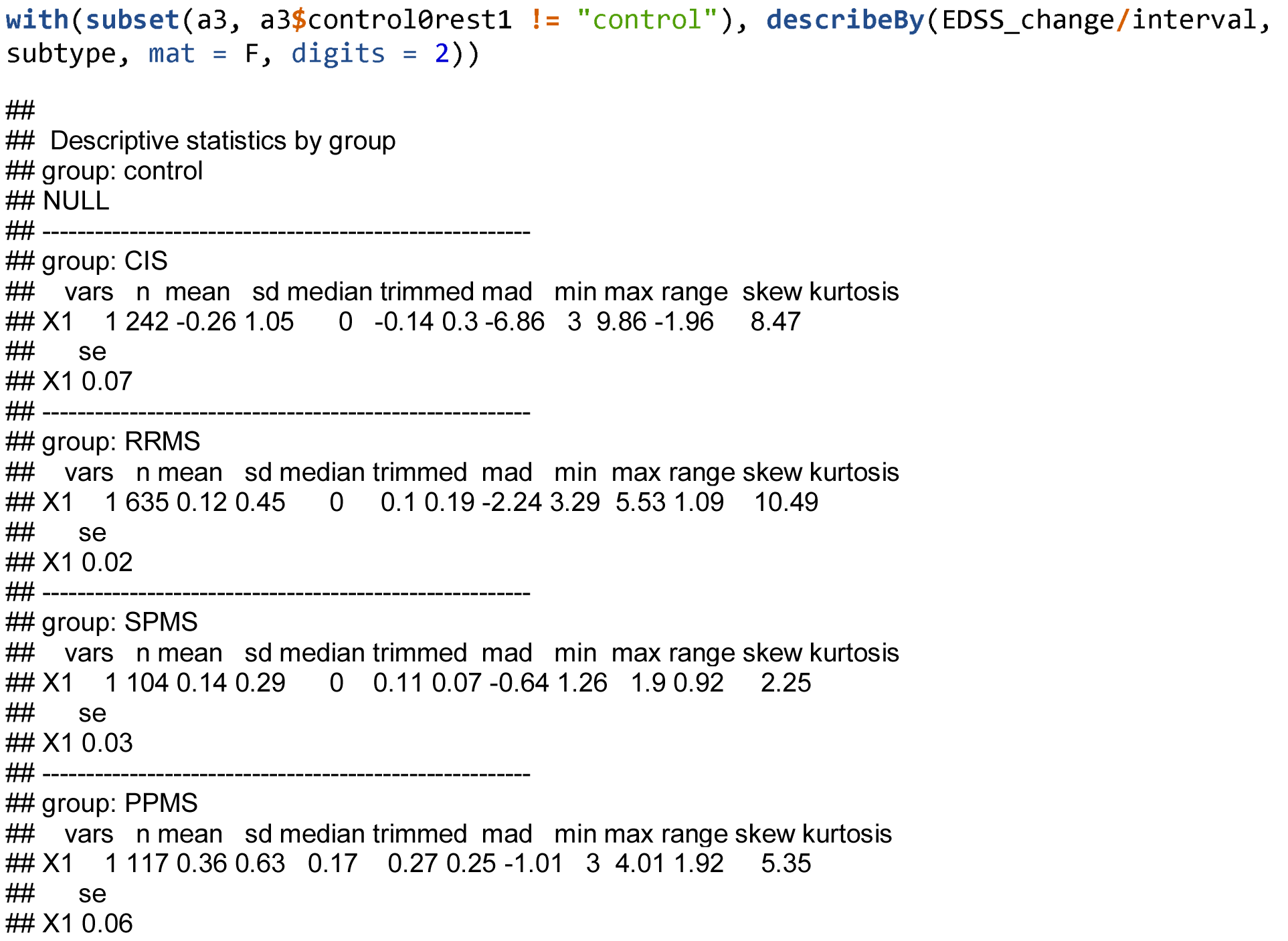

### Correlation between annualised EDSS change and brain-PAD change

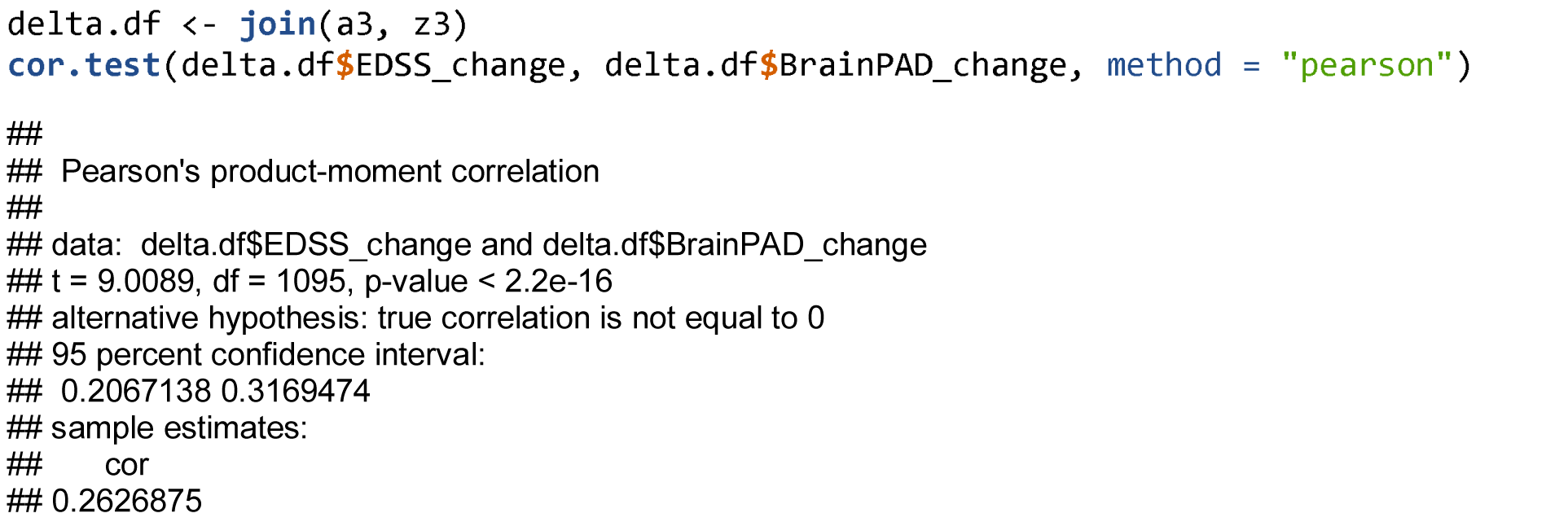

### Interaction between subtype and EDSS change

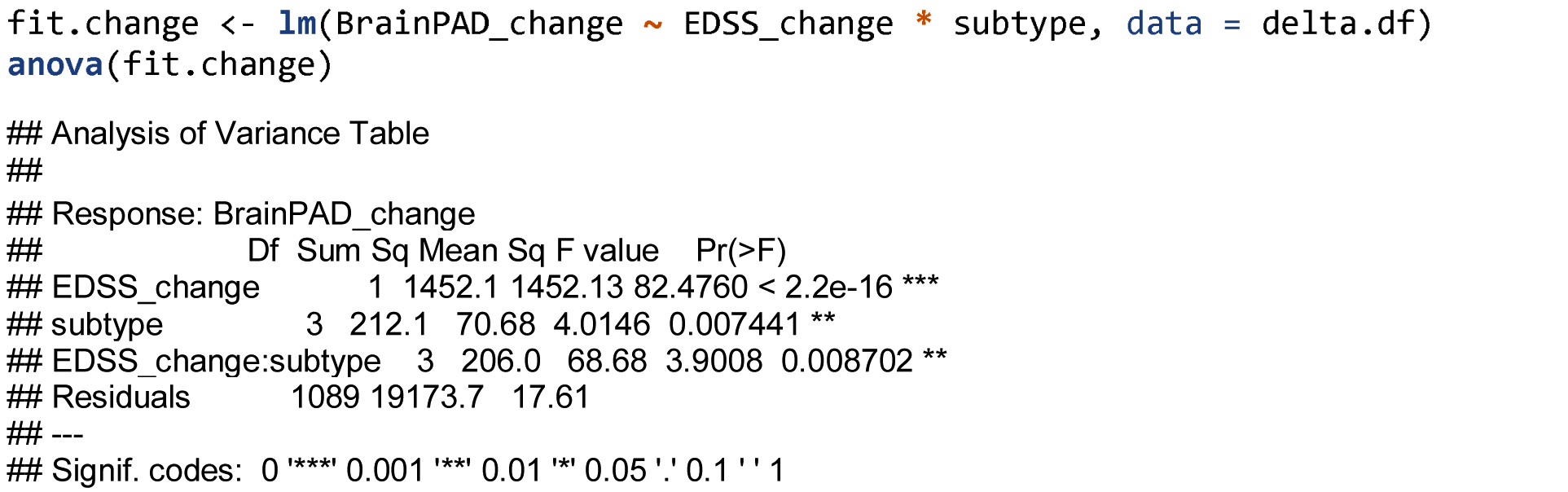

Use jtools package to get slopes from the model, per subtype.

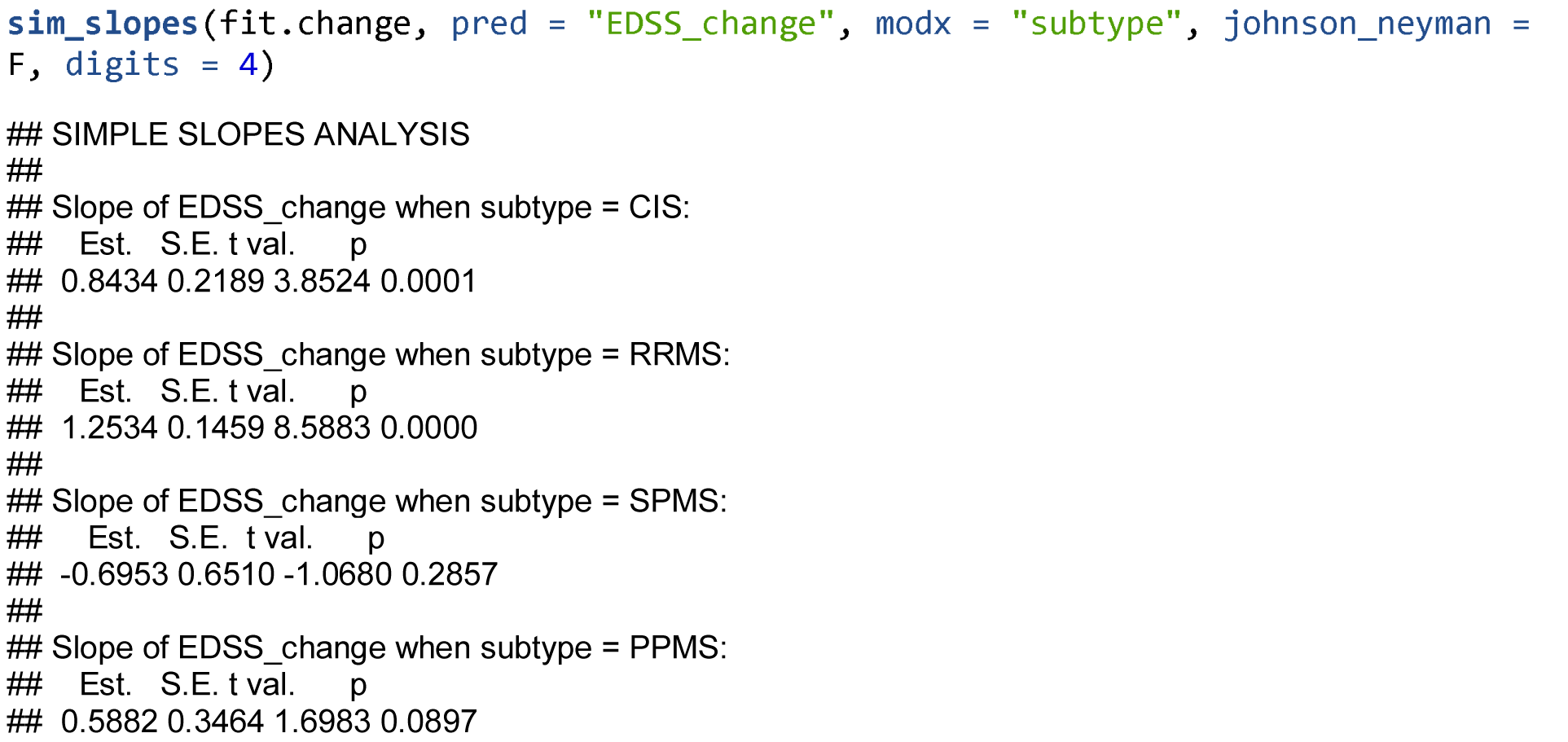

Plot

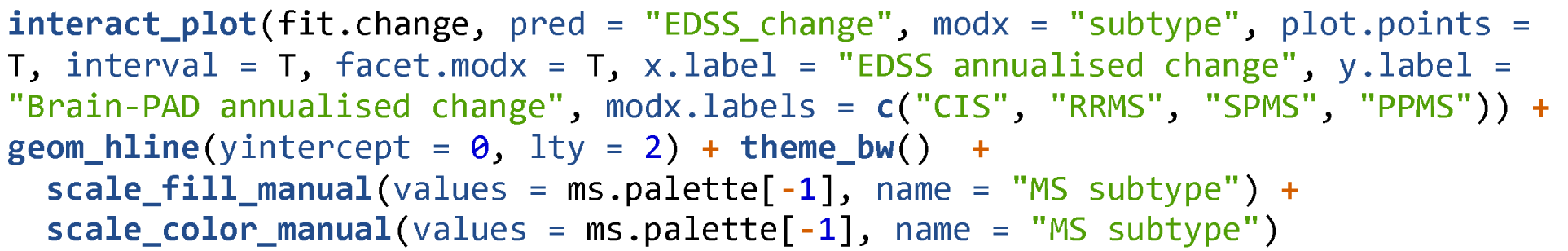

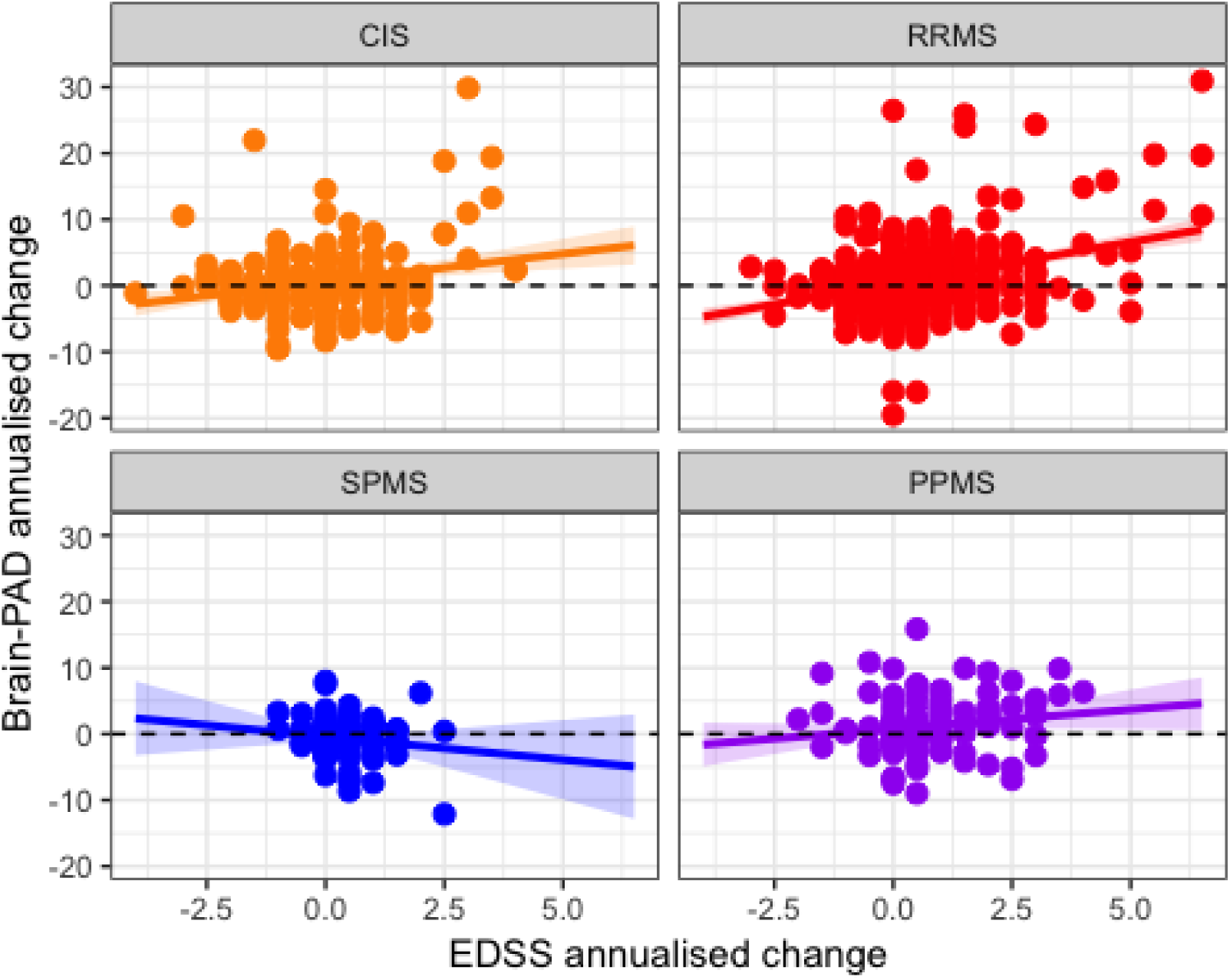

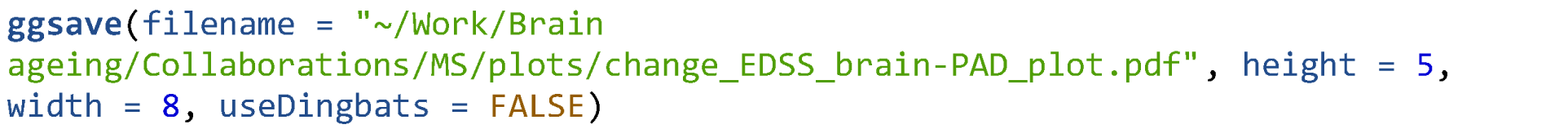

Correlate baseline brain-PAD with the number of follow-up scans completed in the n=104 with >1 scan.

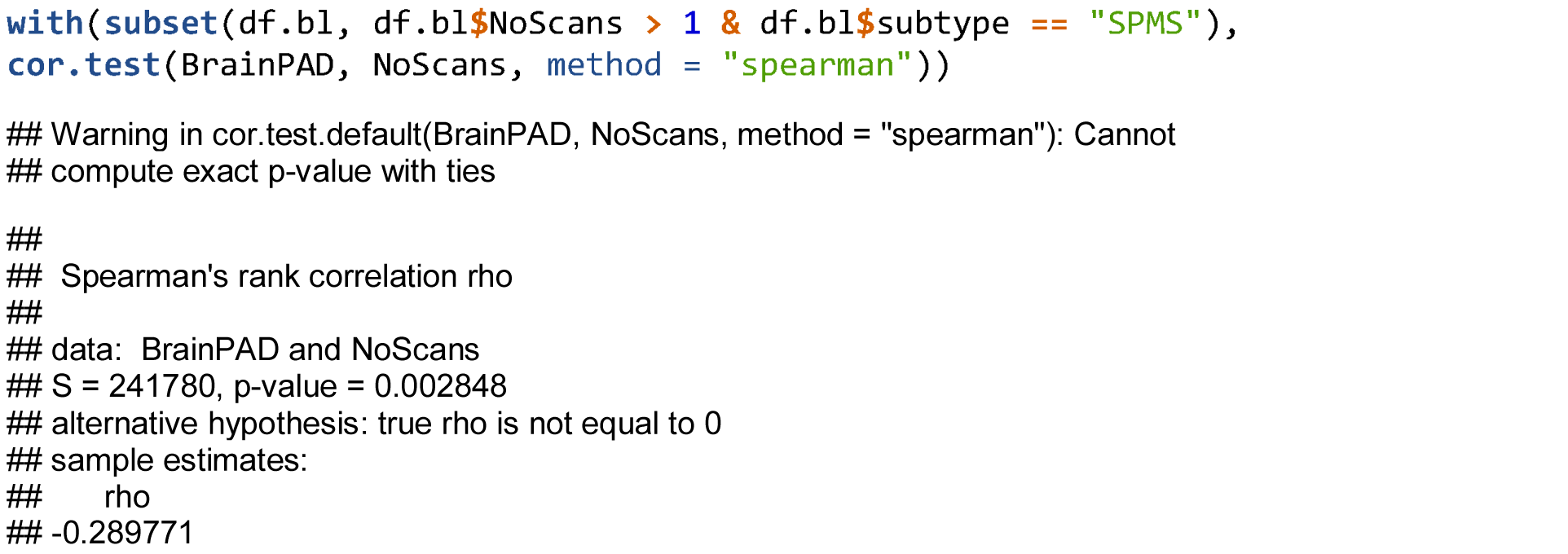

### Longitudinal brain-predicted age trajectories

Interaction between group and time

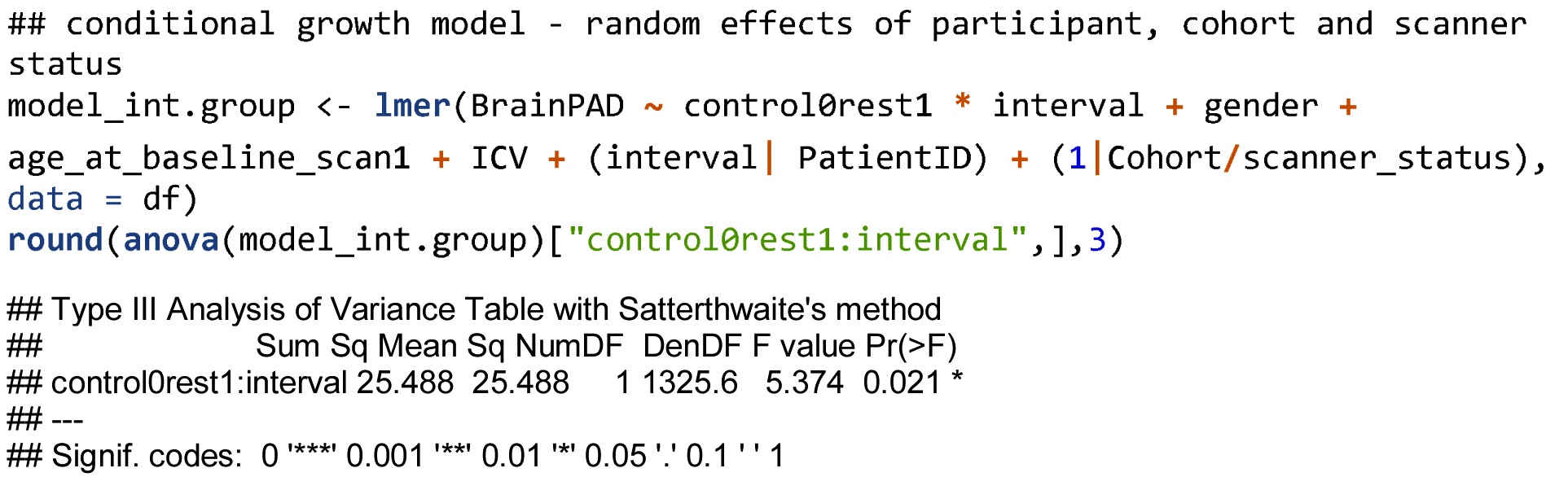

### Brain-PAD change EMMs, using annualised difference between baseline and final follow-up

Generate EMMs for all groups and MS subtypes. LME adjusting for age, gender, ICV, cohort and scanner status.

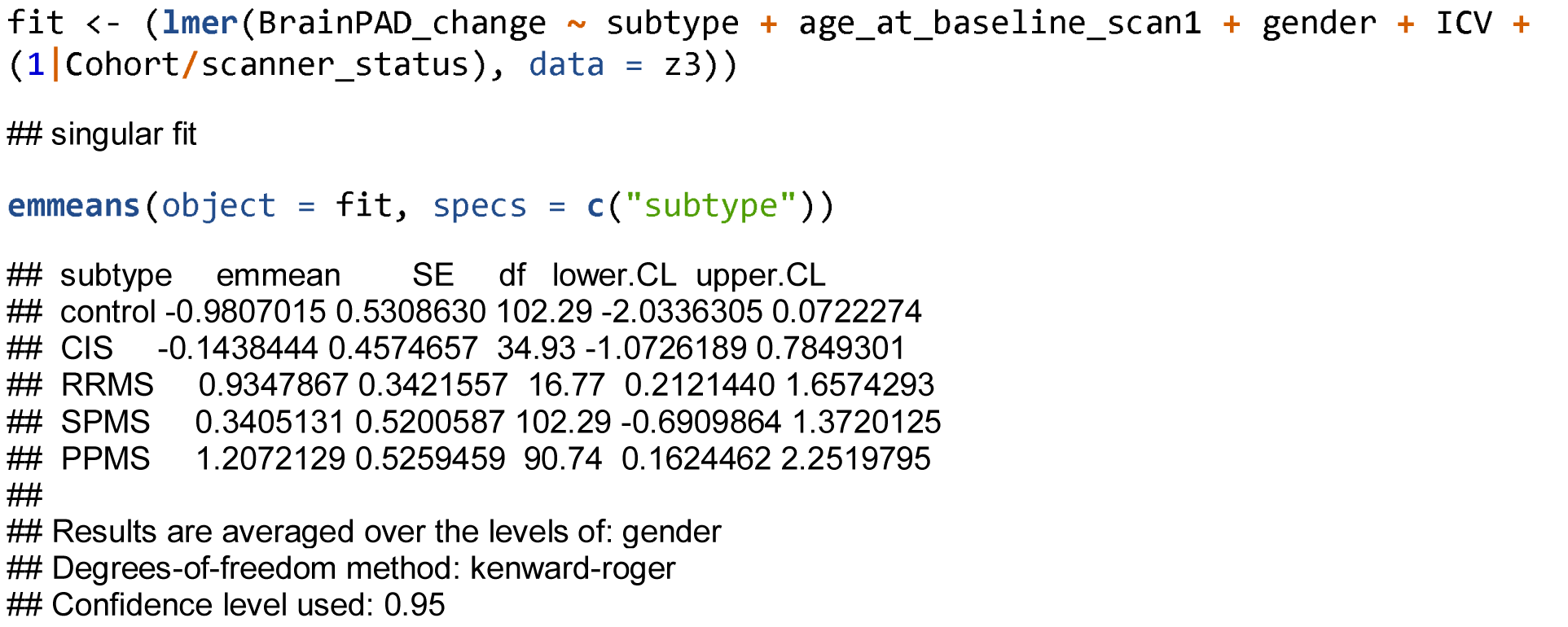

Generate EMMs for HCs and MS/CIS combined. LME adjusting for age, gender, ICV, cohort and scanner status.

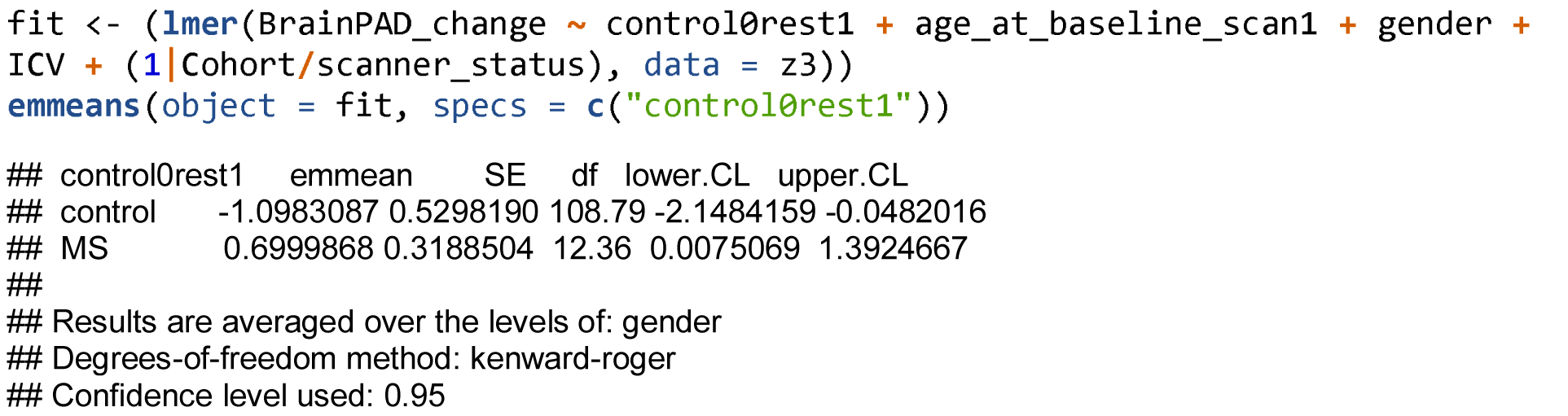

### Slopes per group

Controls

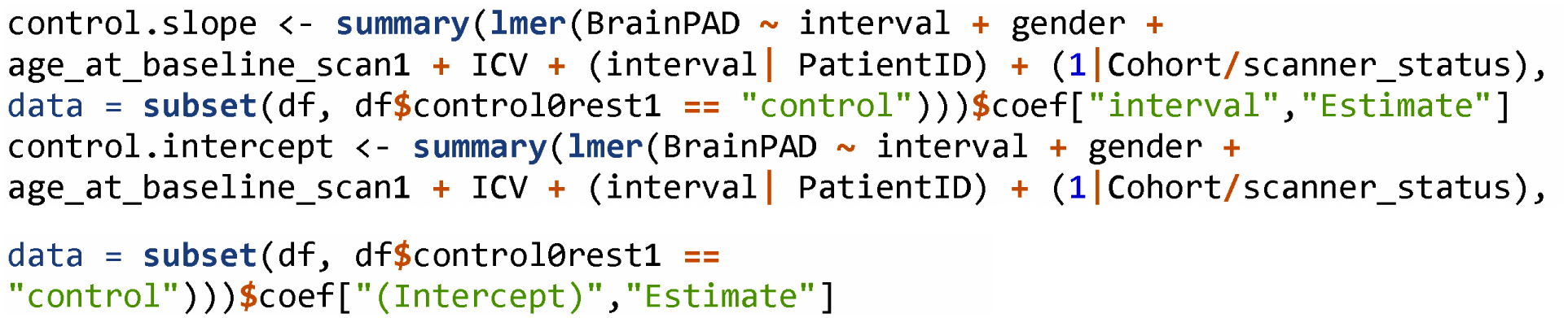

MS patients

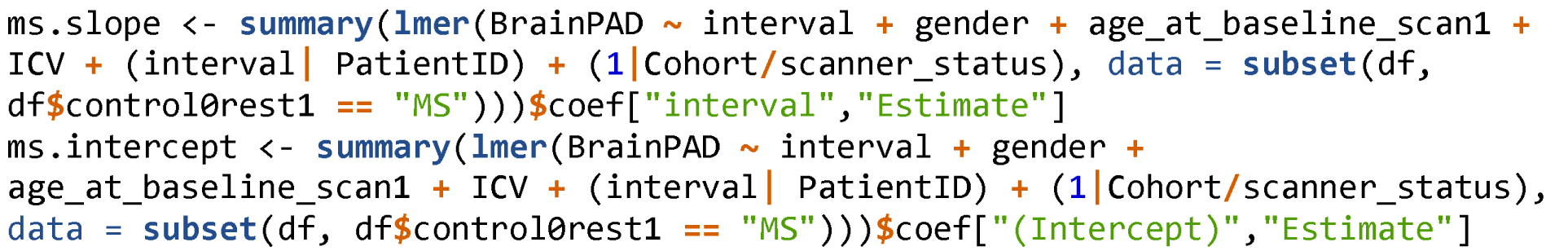

**Longitudinal brain-PAD by interval plots**

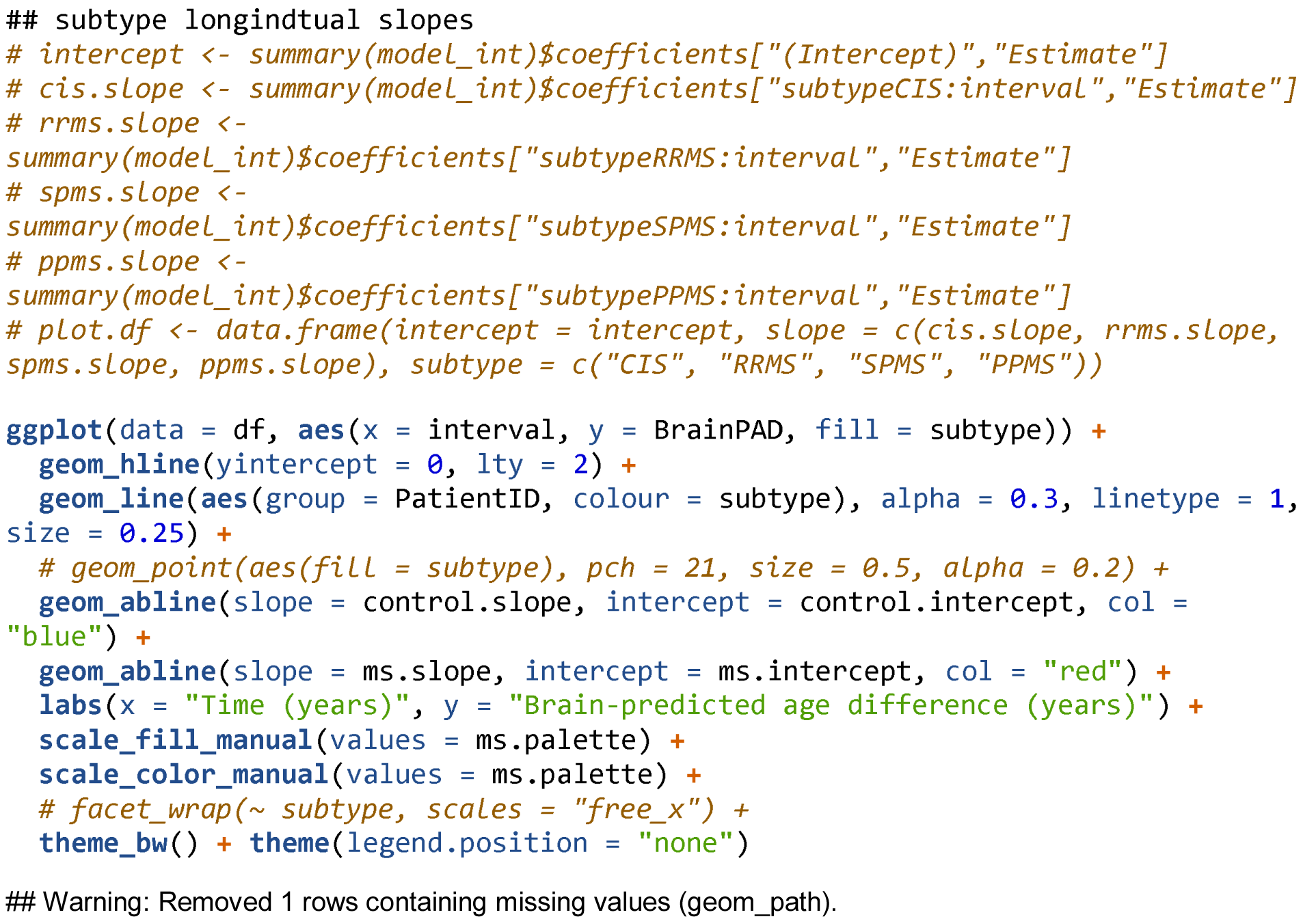

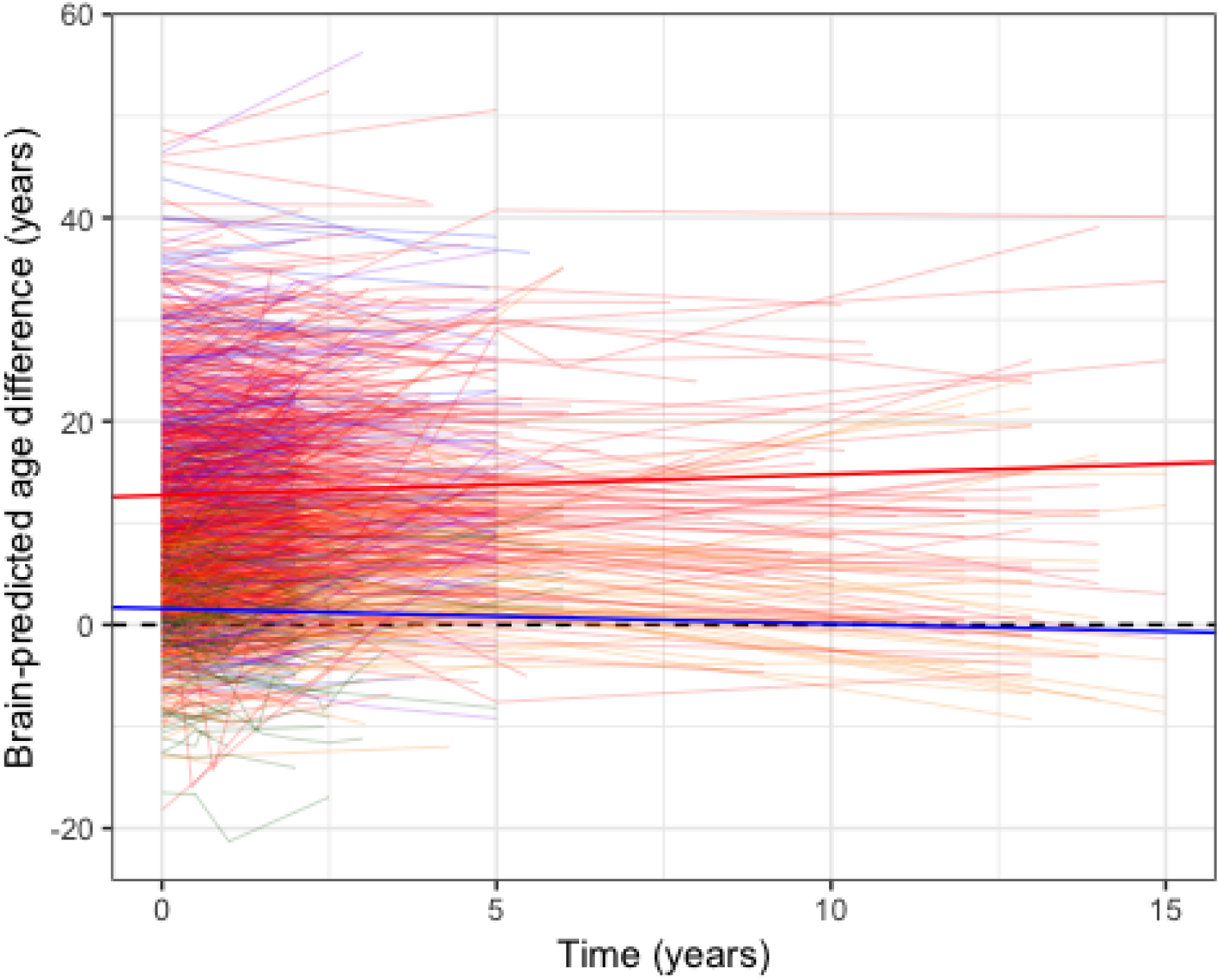

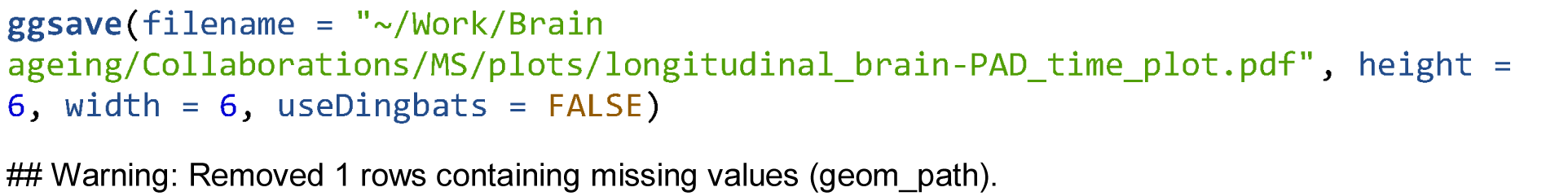

## Supplementary analysis

### Hierarchical partitioning of brain-PAD

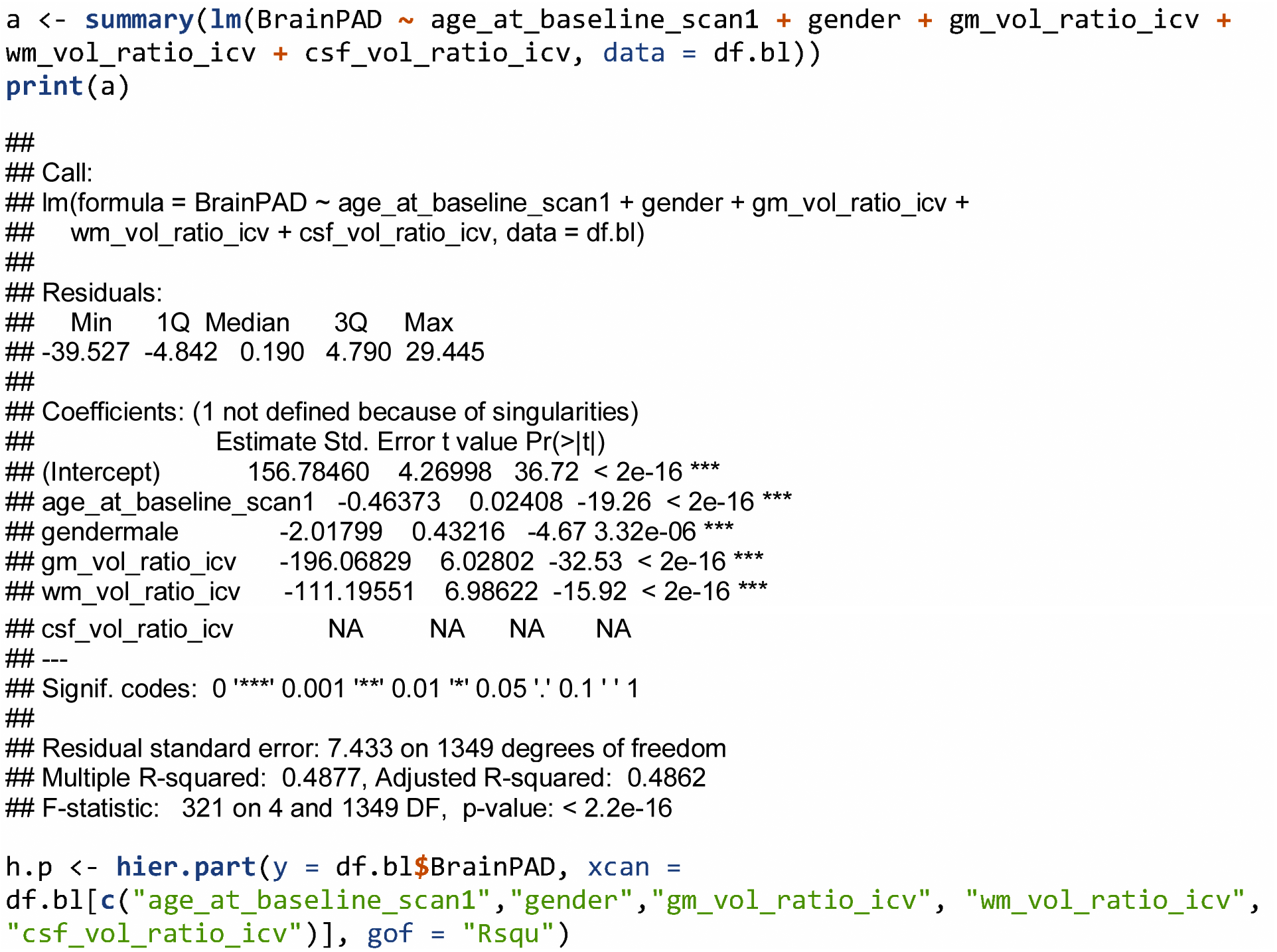

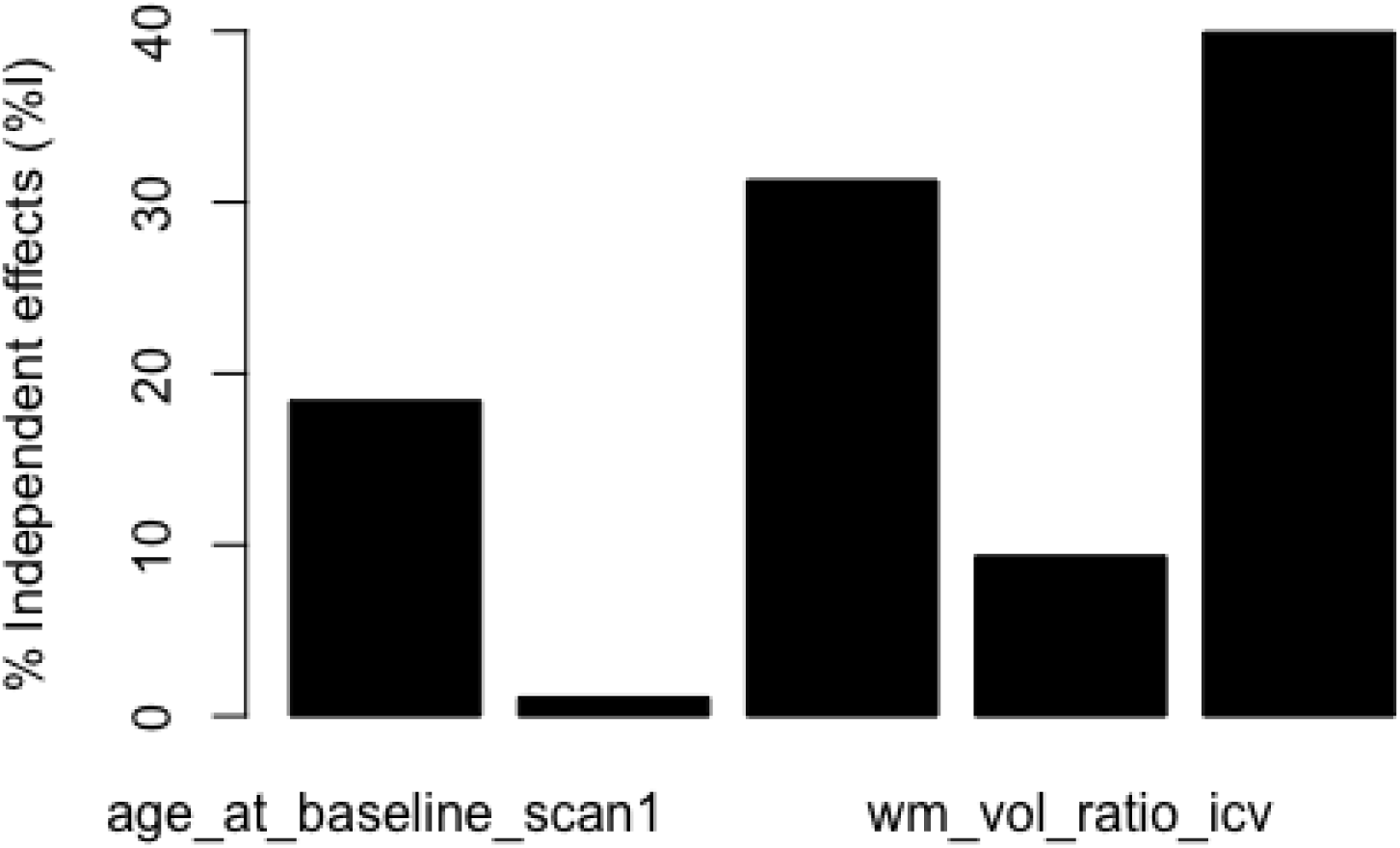

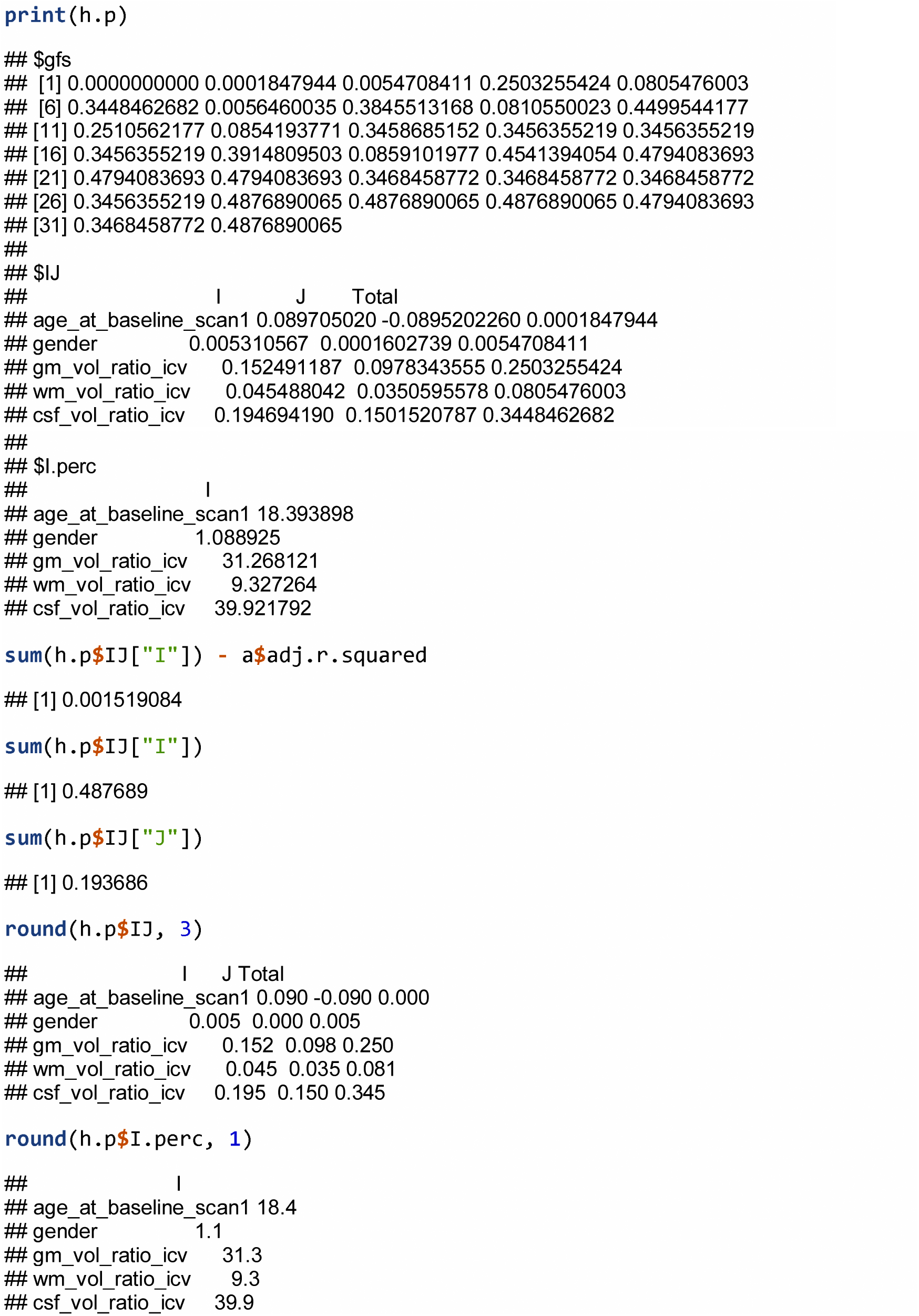

## References

1. Leray E, Yaouanq J, Le Page E, et al. Evidence for a two-stage disability progression in multiple sclerosis. Brain : a journal of neurology 2010; 133(Pt 7): 1900–13.

2. Scalfari A, Lederer C, Daumer M, Nicholas R, Ebers GC, Muraro PA. The relationship of age with the clinical phenotype in multiple sclerosis. Mult Scler 2016; 22(13): 1750–8.

3. Koch M, Mostert J, Heersema D, De Keyser J. Progression in multiple sclerosis: further evidence of an age dependent process. J Neurol Sci 2007; 255(1-2): 35–41.

4. Scalfari A, Neuhaus A, Daumer M, Ebers GC, Muraro PA. Age and disability accumulation in multiple sclerosis. Neurology 2011; 77(13): 1246–52.

5. Tutuncu M, Tang J, Zeid NA, et al. Onset of progressive phase is an age-dependent clinical milestone in multiple sclerosis. Mult Scler 2013; 19(2): 188–98.

6. Oost W, Talma N, Meilof JF, Laman JD. Targeting senescence to delay progression of multiple sclerosis. J Mol Med (Berl) 2018; 96(11): 1153–66.

7. Franceschi C, Garagnani P, Morsiani C, et al. The Continuum of Aging and Age-Related Diseases: Common Mechanisms but Different Rates. Frontiers in Medicine 2018; 5(61).

8. Mattson MP, Arumugam TV. Hallmarks of Brain Aging: Adaptive and Pathological Modification by Metabolic States. Cell Metabolism 2018; 27(6): 1176–99.

9. Cole JH. Neuroimaging Studies Illustrate the Commonalities Between Ageing and Brain Diseases. BioEssays 2018; 40(7): 1700221.

10. Cole JH, Franke K. Predicting Age Using Neuroimaging: Innovative Brain Ageing Biomarkers. Trends in neurosciences 2017; 40 12: 681–90.

11. Franke K, Gaser C. Longitudinal changes in individual BrainAGE in healthy aging, mild cognitive impairment, and Alzheimer’s Disease. GeroPsych: The Journal of Gerontopsychology and Geriatric Psychiatry 2012; 25(4): 235–45.

12. Gaser C, Franke K, Klöppel S, Koutsouleris N, Sauer H, for the Alzheimer’s Disease Neuroimaging Initiative. BrainAGE in mild cognitive impaired patients: predicting the conversion to Alzheimer’s disease. PloS one 2013; 8(6): e67346.

13. Cole JH, Ritchie SJ, Bastin ME, et al. Brain age predicts mortality. Molecular psychiatry 2018; 23: 1385–92.

14. Cole JH, Leech R, Sharp DJ, for the Alzheimer’s Disease Neuroimaging Initiative. Prediction of brain age suggests accelerated atrophy after traumatic brain injury. Ann Neurol 2015; 77(4): 571–81.

15. Cole JH, Underwood J, Caan MWA, et al. Increased brain-predicted aging in treated HIV disease. Neurology 2017; 88(14): 1349–57.

16. Cole JH, Annus T, Wilson LR, et al. Brain-predicted age in Down Syndrome is associated with β-amyloid deposition and cognitive decline. Neurobiology of aging 2017; 56: 41–9.

17. Pardoe HR, Cole JH, Blackmon K, Thesen T, Kuzniecky R. Structural brain changes in medically refractory focal epilepsy resemble premature brain aging. Epilepsy Research 2017; 133: 28–32.

18. Eshaghi A, Prados F, Brownlee Wallace J, et al. Deep gray matter volume loss drives disability worsening in multiple sclerosis. Ann Neurol 2018; 83(2): 210–22.

19. Polman CH, Reingold SC, Banwell B, et al. Diagnostic criteria for multiple sclerosis: 2010 revisions to the McDonald criteria. Ann Neurol 2011; 69(2): 292–302.

20. Lublin FD, Reingold SC, Cohen JA, et al. Defining the clinical course of multiple sclerosis: the 2013 revisions. Neurology 2014; 83(3): 278–86.

21. Kurtzke JF. Rating neurologic impairment in multiple sclerosis: an expanded disability status scale (EDSS). Neurology 1983; 33(11): 1444–52.

22. Schrouff J, Rosa MJ, Rondina JM, et al. PRoNTo: Pattern recognition for neuroimaging toolbox. Neuroinformatics 2013; 11(3): 319–37.

23. Battaglini M, Jenkinson M, De Stefano N. Evaluating and reducing the impact of white matter lesions on brain volume measurements. Hum Brain Mapp 2012; 33(9): 2062–71.

24. Degenhardt A, Ramagopalan SV, Scalfari A, Ebers GC. Clinical prognostic factors in multiple sclerosis: a natural history review. Nature reviews Neurology 2009; 5(12): 672–82.

25. Gooch CL, Pracht E, Borenstein AR. The burden of neurological disease in the United States: A summary report and call to action. Ann Neurol 2017; 81(4): 479–84.

26. Uher T, Vaneckova M, Krasensky J, et al. Pathological cut-offs of global and regional brain volume loss in multiple sclerosis. Mult Scler 2017: 1352458517742739.

27. Rocca MA, Battaglini M, Benedict RH, et al. Brain MRI atrophy quantification in MS: From methods to clinical application. Neurology 2017; 88(4): 403–13.

28. Montalban X, Gold R, Thompson AJ, et al. ECTRIMS/EAN Guideline on the pharmacological treatment of people with multiple sclerosis. Mult Scler 2018; 24(2): 96–120.

29. Kaufmann T, van der Meer D, Doan NT, et al. Genetics of brain age suggest an overlap with common brain disorders. bioRxiv 2018.

30. Cole JH, Poudel RPK, Tsagkrasoulis D, et al. Predicting brain age with deep learning from raw imaging data results in a reliable and heritable biomarker. NeuroImage 2017; 163C: 115–24.

